# Inferring number of populations and changes in connectivity under the n-island model

**DOI:** 10.1101/2020.09.03.282251

**Authors:** Armando Arredondo, Beatriz Mourato, Khoa Nguyen, Simon Boitard, Willy Rodríguez, Camille Noûs, Olivier Mazet, Lounès Chikhi

## Abstract

Inferring the demographic history of species is one of the greatest challenges in populations genetics. This history is often represented as a history of size changes, thus ignoring population structure. Alternatively, structure is defined a priori as a population tree and not inferred. Here we propose a framework based on the IICR (Inverse Instantaneous Coalescence Rate), which can be estimated using the PSMC method of Li and Durbin (2011) for a single diploid individual. For an isolated population, the IICR matches the population size history, which is how the PSMC outputs are generally interpreted. However, it is increasingly acknowledged that the IICR is a function of the demographic model and sampling scheme. Our automated method fits observed IICR curves of diploid individuals with IICR curves obtained under piecewise-stationary symmetrical island models, in which we assume a fixed number of time periods during which gene flow is constant. We infer the number of islands, their sizes, the periods at which connectivity changes and the corresponding rates of connectivity. Validation with simulated data showed that the method can accurately recover most of the scenario parameters. Our application to a set of five human PSMCs yielded demographic histories that are in agreement with previous studies using similar methods and with recent research suggesting ancient human structure. They are in contrast with the widely accepted view of human evolution consisting of one ancestral population branching into three large continental and panmictic populations with varying degrees of connectivity and no population structure within each continent.

## 1. Introduction

Reconstructing the demographic history of populations from the analysis of genomic data is one of the greatest challenges of population geneticists (Goldstein and Chikhi, 2002; Hey and Machado, 2003; Beaumont, 2004; Li and Durbin, 2011). It is an important and challenging statistical problem, but it is also central to our understanding the evolutionary history of species. Indeed, the demographic history that we assume or infer for a particular population or species implicitly or explicitly provides the null model against which selected loci could in theory be identified (Cavalli-Sforza, 1966; Rogers and Harpending, 1992; Beaumont and Nichols, 1996; Wakeley, 1999; Goldstein and Chikhi, 2002). In a period of global environmental change, the reconstructed demographic history should allow evolutionary biologists to correlate changes in population size or connectivity with past environmental changes or species association and interactions (Goossens et al., 2006; Quéméré et al., 2012; Salmona et al., 2017; Hecht et al., 2018, 2020).

In other words, we expect to increase our understanding of the environmental (including species interaction) and anthropogenic factors that have influenced genomic diversity in various species. Understanding how past climatic events or human activities have influenced genomic diversity may become particularly illuminating for conservation biologists regarding the likely long-term consequences of ongoing climate change and human actions.

However, to understand how the past influenced the present patterns of genomic diversity one major question is whether our conclusions or inferences may fundamentally change depending on the family of models assumed and the questions asked (Wakeley, 1999; Beaumont, 2004; Chikhi et al., 2010; Mazet et al., 2016; Chikhi et al., 2018; Rodríguez et al., 2018; Poelstra et al., 2020). Currently, the solutions to this complex inferential problem have been to either ignore population structure (i.e., assume that differentiation between geographic locations can be neglected and panmixia assumed) and infer population size changes, or to assume simplified tree models with *a priori* fixed numbers of populations (i.e., the population tree used in these studies is assumed, not inferred). Additionally, in the case of human evolutionary history, the branches of the assumed tree may represent predefined continental populations. Such models thus assume panmixia over large geographic regions and long periods (Gutenkunst et al., 2009; Mazet et al., 2016; Noskova et al., 2019; Scerri et al., 2019). Panmictic and tree models are useful approximations, and in the last decades they have proven their utility in building stories of human expansions and population splits (Gutenkunst et al., 2009; Li and Durbin, 2011). However, most tree models assume the existence of clear splitting events similar to those separating species. Also, some tree models assume, as in most species trees, that there is no gene flow between branches (even when they represent populations or continents). The latter assumption may then require the inference of admixture events (Kuhlwilm et al., 2016) in order to explain the presence of outlier or divergent alleles that cannot be explained by the tree models if they exclude population structure and gene flow.

Methods can also be classified by the type of data used. Currently, most genomic methods use the allele frequency spectrum (AFS) (Gutenkunst et al., 2009; Excoffier et al., 2013; Liu and Fu, 2015) or the AFS combined with other summary statistics (Boitard et al., 2016). The AFS can be computed from RAD-Seq data for many non-model species or from full genome for a still limited number of species. We stress though that this research field is very active and new methods are regularly proposed that go beyond the simplified classifications proposed here. For instance, a recent method implemented in the diCal2 software allows to infer complex demographic histories from full genomes (Steinrücken et al., 2019).

Authors have focused on panmictic models to infer population size changes (Li and Durbin, 2011; Schiffels and Durbin, 2013) but we note that this is not always the case. For instance, the MSMC methods developed in Schiffels and Durbin (2013); Wang et al. (2020) also allow inferring migration history between a pair of populations considered isolated from the others, which allows to estimate the splitting time between these populations and to build assumptions about admixture events involving only one of them.

Here, we propose to use a different strategy based on the IICR (Inverse Instantaneous Coalescence Rate) introduced by Mazet et al. (2016). We propose to perform demographic inference under the piecewise stationary *n*-island model (Rodríguez et al., 2018), based on the symmetrical *n*-island model (Wright, 1931), using the IICR or estimates of it. The IICR, as defined by Mazet et al. (2016) for a sample of size two, is equivalent to the full distribution of coalescence times for that sample (i.e., the distribution of *T_2_*). Simulations by Chikhi et al. (2018); Rodríguez et al. (2018) under various models of population structure suggest that it is also a highly informative statistical summary of the genomic data on the demographic history. This makes the IICR a good choice for use in inference applications particularly due to its sensitivity to population structure or fluctuations in migration rates (i.e., changes in connectivity). In figure 1, we present an example from Mazet et al. (2016) where a spurious bottleneck effect due to population structure is so strong that it hides the increase in size of all demes in the recent past. This example stresses how it could be misleading to interpret the IICR as a change in population size, but also how structure can profoundly impact genomic patterns of diversity.

**Figure 1:**
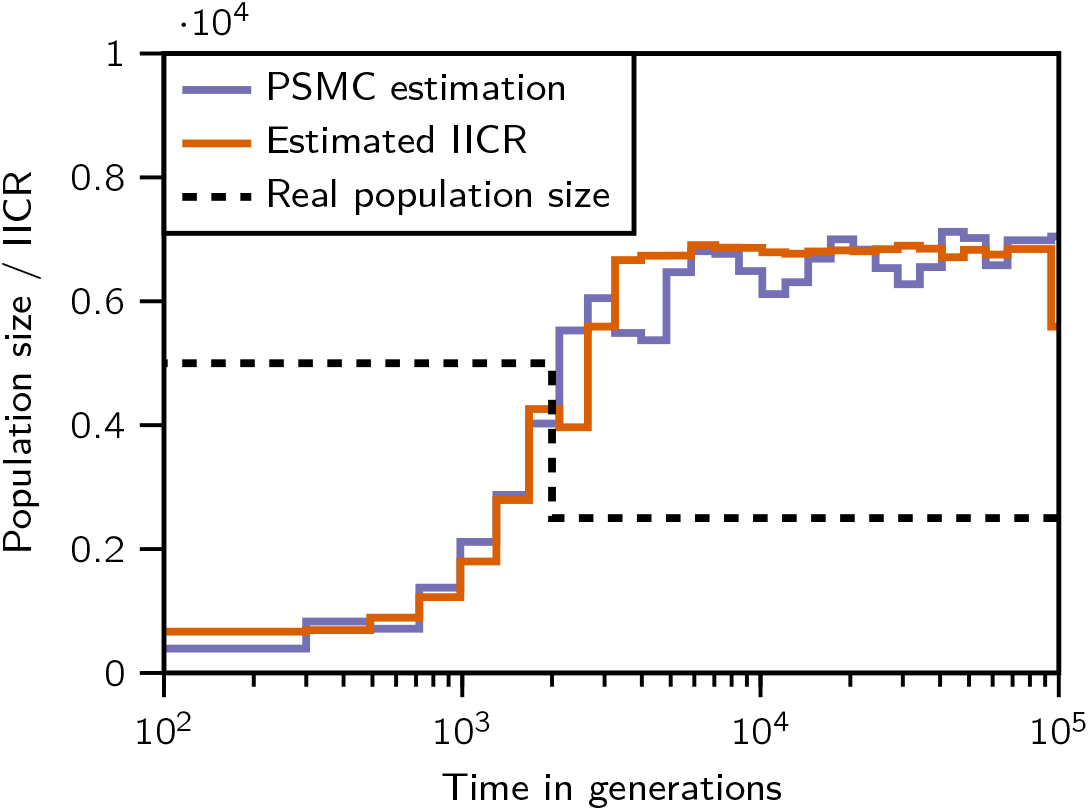
A reading of this IICR curve as a size change function would indicate that the population has decreased in size in the recent past, where in fact the opposite is true and the population has experienced a doubling of size in the recent past.

The approach presented in the present study differs from the approaches mentioned above in several ways. First, we aim at *inferring* the number of populations rather than setting it *a priori*. Second, we ask whether it is possible to date and quantify changes in connectivity (i.e., gene flow) rather than changes in population size. For that we use the piecewise stationary *n*-island model in which continuous gene flow happens between populations at a constant rate during specific periods (called *components*) but is allowed to change between periods (see below and Rodríguez et al. (2018)). This model differs from tree models in that we do not estimate parameters such as splitting times which may or may not be meaningful or appropriate for various species (Scerri et al., 2019) depending on their actual (unknown) demographic history. Our approach could be extended to more complex models. We acknowledge the limitations of the *n*-island model as it ignores spatial distances and other complexities of real species (Chikhi et al., 2018), but our choice for the current study is also guided by simplicity and computational considerations. See Rodríguez et al. (2018) for a more complex scenario involving two piecewise stationary *n*-island models connected by a tree structure. That study also shows that the IICR can be efficiently computed for the piecewise stationary *n*-island, where the number of *components* thus corresponds to the number of piecewise periods of constant gene flow. This means that it is possible to predict the shape of the IICR for any reasonably large number of islands and any rate of gene flow for a fixed number of *components*. It is thus theoretically possible to increase our understanding of ancient patterns of connection and disconnection by using a single genome and the corresponding IICR under the *n*-island model. Given that such changing patterns of connectivity may have been crucial in the recent evolutionary history of many species (Goldstein and Chikhi, 2002; Quéméré et al., 2012; Mazet et al., 2016; Salmona et al., 2017; Scerri et al., 2018; Steinrücken et al., 2019; Fenderson et al., 2020), particularly in the context of Pleistocene climate change and habitat fragmentation, the work presented here may represent an interesting endeavour.

To use the IICR as a summary of genomic information we first assume that an IICR curve can be obtained, which we will use as the *target* for demographic inference. With simulated data (sequences or *T*_2_ values) this target curve can be obtained under any pre-defined coalescent model that could be expressed with a simulation tool (e.g., the ms program). With real genome-wide sequence data, the curve can be estimated with the PSMC method of Li and Durbin (2011). We then try to identify a piecewise stationary *n*-island model that generates an IICR that is identical or similar to the target IICR (or PSMC curve). The similarity between the two IICR curves is quantified with a distance metric defined below. We use a genetic algorithm to explore the parameter space (number of populations, migration rate within a time component, and timing of these changes assuming a fixed number of components for each independent analysis) and minimize that distance. We compute the IICR under the non-stationary structured coalescent (NSSC) of Rodríguez et al. (2018). The inferential method is first validated on simulated data across a wide range of parameter values, spanning many values for the number of islands, the number of demographic events which define the model components and the values of the migration rates during these components. We compare estimated and simulated parameter values and also infer what we call connectivity graphs which are a visual representation of the times at which gene flow changed and the magnitude of these changes. We limit ourselves to validating the method with data simulated under piecewise stationary *n*-island models due to the very large parameter space explored here. We then apply our approach to human genomic data using five published PSMC curves (Prado-Martinez et al., 2013), allowing in each case the number of components to vary between analyses, and compare the inferred histories and connectivity graphs between individuals and with previously inferred scenarios by Rodríguez et al. (2018) and Noskova et al. (2019).

Altogether our simulation results are promising as they show that the IICR is a highly informative summary statistic. We find that most parameters are extremely well estimated when the IICR is well estimated and the number of *components* is less than *five or six* depending on the timing of events. For scenarios with many components our results suggest that the connectivity graphs can be used to identify meaningful models (see §4). For the human data we find that the inferred scenarios are similar and consistent with previous scenarios by Rodríguez et al. (2018) across most individuals.

Beyond human data we find that a crucial issue is the estimation of the IICR from genomic data. Indeed, the stochasticity generated during the estimation of the IICR in very ancient times, and possibly recent times, with humps that are difficult to interpret, may lead to the inference of events that may never have taken place.

## 2. Methods

### 2.1. The structured coalescent and the IICR

The theoretical framework we use for modelling structure is based on the finite Herbots model of the structured coalescent (Herbots, 1994), where we have *n* populations or demes that are assumed to behave as haploid Wright-Fisher models of size *N_i_* = 2*s_i_N* genes, where *s_i_* is the relative deme size and *N* is large. Migration occurs between demes as in each generation a proportion *q_ij_* of lineages migrates from deme *i* to deme *j*. Herbots denoted by *m_i_* the proportion of the population of deme *i* that was received from other demes in any given generation, such that 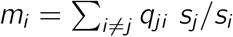. She also showed that measuring time in units of 2*N* generations and making *N* go to infinity in such a way that the number of migrants stays bounded, the model converges to a continuous-time Markov process with a known transition rate matrix *Q* that is a good approximation of this discrete time model. In this transformation, *q_ij_* goes to zero in such a way that the product 2*Nq_ji_ s_j_*/*s_i_* converges and has limit *M_ij_*/2. Thus we can express the transition rates in *Q* in terms of *n, s_i_* and *M_ij_*. In the case of the symmetrical island model (Wright, 1931), all the migration rates *M_ij_* between any pair of islands *i* and *j* are equal, so we use the notation *M* = (*n* – 1)*M_ij_* to denote the migration rate perceived by any given island. In addition to this base model, we use an extension presented in Rodríguez et al. (2018) which allows to introduce demographic events that change the rate *M* or relative deme size *s* at certain points in time (see §2.2). We note however that throughout the manuscript we will only focus on symmetrical models with constant size (see §4 for extensions to symmetrical models with population size changes).

For the demographic histories under these models, we study the IICR of a sample of size 2 (see §2.2.2), and we use it as a statistic for demographic inference. We do this by comparing the IICR of many hypothetical demographic scenarios to a target IICR curve. This target IICR may be simulated, or it may be obtained from diploid individuals via full genome studies (Prado-Martinez et al., 2013). In such cases, these target IICRs are themselves inferred demographies under the assumptions of a particular model. For example, the PSMC method (Li and Durbin, 2011) uses the population size change model, where a single panmictic population varies in size according to a function *N*(*t*) = *N*(0)λ(*t*) (see Tavaré (2004)). It was shown by Mazet et al. (2016) that the IICR of a sample of size 2 under this model is exactly the λ(*t*) relative size changing function, and it relates to the distribution of the time to coalescence *T*_2_ as:

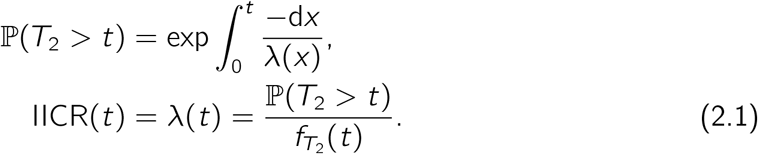

The IICR is not tied to any particular model, structured or otherwise. It is defined using the distribution of the coalescent times of a sample of size two. It can be approximated to arbitrary numerical precision under the assumptions of the NSSC (Rodríguez et al., 2018); it can also be computed empirically by simulating a sample of coalescent times (Chikhi et al., 2018); or it can be read from full sequence genomic data using the appropriate methods (Li and Durbin, 2011). Such sequences can be real or simulated.

### 2.2. The piecewise stationary n-island coalescent

#### 2.2.1. The parameter space

We first define the parameter space, as this directly determines the family of demographic histories that we can explore and infer from. In this work we limit our scope to piecewise-stationary symmetrical *n*-island models. The piecewise stationarity refers to the fact that, although migration rate is constant and identical between any pair of islands, this rate may be different between consecutive time periods (components), and there is a fixed number *γ* that represents the number of demographic events. To say that there are γ changes of gene flow thus means that there are *γ* + 1 components or periods of constant gene flow. Likewise, the deme size, which is the same for all islands, may in theory change through time in the general model presented in Rodríguez et al. (2018). In the present study we focus on models with constant population size but we present here a more general model where deme sizes could change between components. In this more general case, the parameter space for defining any given model includes the number of islands *n*, the times *t_i_* for the demographic events, and the values of both the migration rate *M_i_* and the local deme size *s_i_* at each new demographic period (see Rodríguez et al. (2018) for a more general introduction to these types of models). As noted above, there are *c* = *γ* + 1 *components*, between which the migration rate *M_i_* and local deme size *s_i_* can change. The number of islands is inferred as one of the model parameters, but it does not change through time, so it remains constant across components. We thus assume no extinction, no population split and no creation of new populations, but we try to find the number of islands that best explains the data, and this is not fixed a priori.

To formalize the notion of parameter space, given a fixed integer γ of demographic events to consider (*γ* ⩾ 0) and a collection of bounds *B* in the form of:

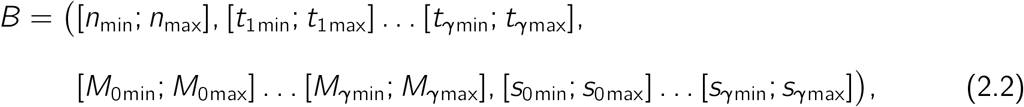

we define the parameter space Φ_*γ,B*_ as:

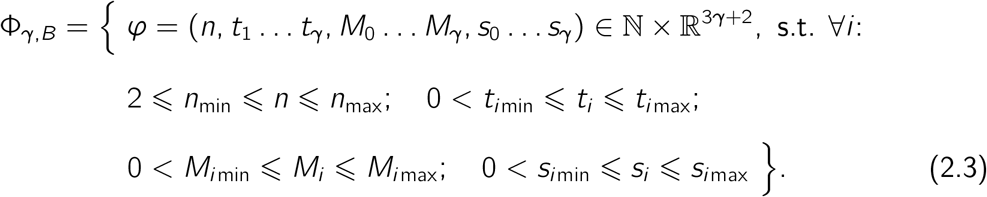

We define bounds for each variable because we use a constrained optimization algorithm, for which all parameters must be bounded (see §2.3). Also, since we focus on the case where there are no deme size changes, we enforce this by using *B*, as making *s*_*i* min_ = *s*_*i* max_ = 1 for all 0 ⩽ *i* ⩽ *γ* effectively fixes all deme sizes to 1.

**Figure 2:**
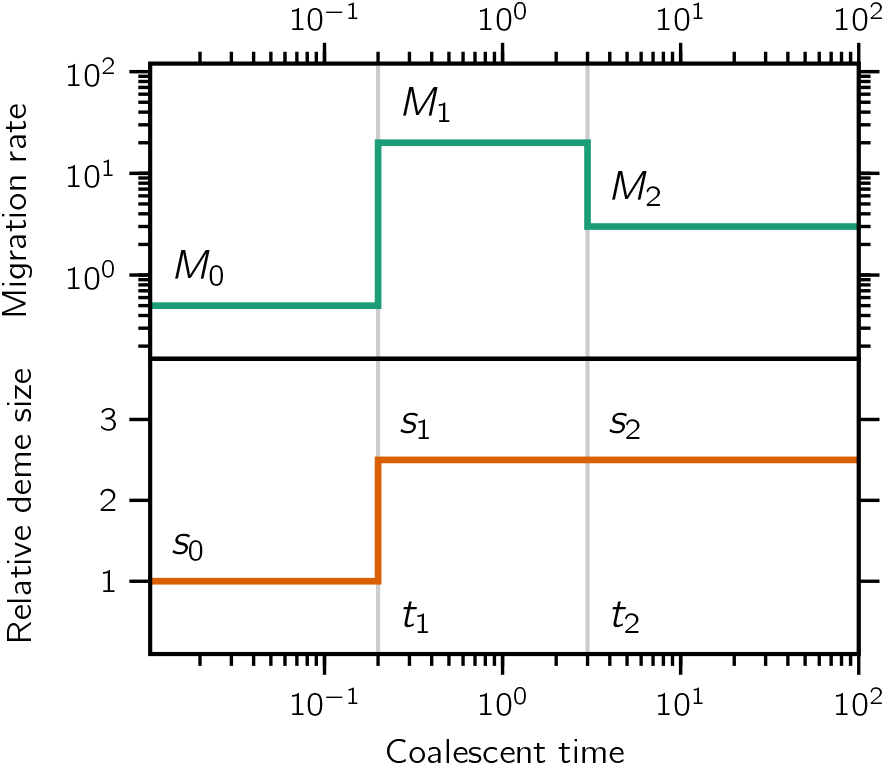
Connectivity and population size graphs. Visual representation of the *t_i_, M_i_* and *s_i_* of a demographic model with *c* = 3 components. We refer to the upper part of the figure as a connectivity graph. The lower part represents population size changes of the general model. In the present study the s, are identical and equal to 1 and will not be represented with connectivity graphs. The number of islands is inferred but constant, and it is not shown in this figure (although it is usually shown in a separate figure or panel).

#### 2.2.2. Computing the IICR

Given any demographic scenario from Φ_*γ,B*_, the associated coalescent process is an instance of the NSSC of Rodríguez et al. (2018). Since our main object of interest regarding these scenarios is the IICR, we proceed with a brief overview of how to perform its computation for any given *φ* ∈ Φ_*σ,B*_.

Considering again the c components which constitute the model, we note that during component *i*, taking 0 ⩽ *i* ⩽ *γ*, the underlying coalescent process *X_t_* is being governed by an *n*-island model which has transition rate (see Rodríguez et al. (2018) for details and a more general approach):

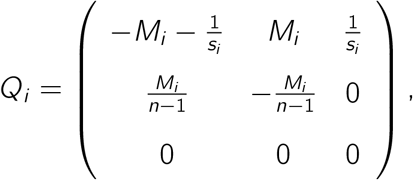

where the three states in the matrix *Q_i_* are sorted, both by row and column, as:

1. the two sampled lineages are in the same deme (configuration ‘same’);
2. the two sampled lineages are in different demes (configuration ‘diff.’);
3. the sampled lineages have coalesced (absorption).

Additionally, the rates in this matrix are active during the time interval [*t_i_*; *t*_*i*+1_), where we understand *t*_0_ and *t*_*γ*+1_ to be 0 and +∞ respectively.

We know from that the probability distribution of *T*_2_ is given by the cumulative effects of the exponential functions e^*tQ_i_*^ (see Hobolth et al. (2019) for an extensive review). More formally, given any *t* > 0, let *i* be the largest index such that *t_i_* ⩽ *t*, for which case we have:

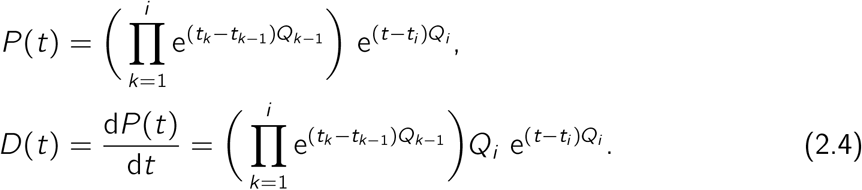

Since component (*p, q*) in matrix *P*(*t*) gives the probability 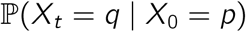, then the random variable *T*_2,same_ of the time until coalescence of two lineages sampled in the *same* deme (initial state 1) has distribution *F*_same_(*t*) = *P*(*t*)_(1,3)_ and density *f*_same_(*t*) = *D*(*t*)_(1,3)_, and for the case where we sample in two *different* demes (initial state 2), then *T*_2,diff_. would have its distribution given by *F*_dff_.(*t*) = *P*(*t*)_(2,3)_ and its density by *f*_diff_.(*t*) = *D*(*t*)_(2,3)_.

The factor matrices in (2.4) can be computed in several ways in the general case (see Herbots (1994) or Hobolth et al. (2019)), but considering this particular instance of size 3 × 3, they may also be computed directly given an arbitrary Δ*t* and rate matrix *Q*:

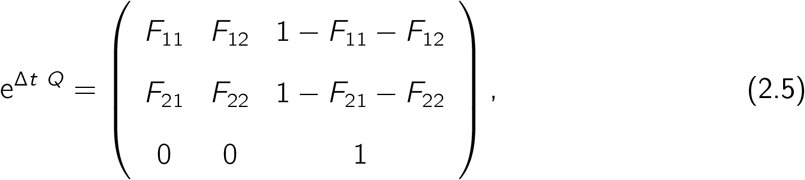

where:

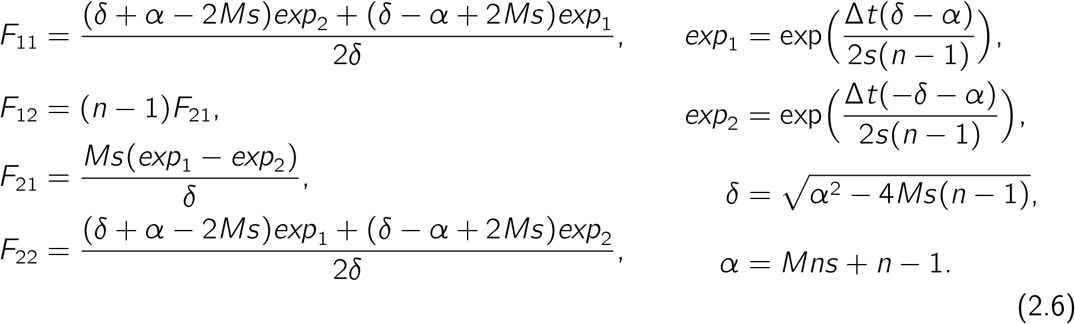

With this we can efficiently compute either IICR functions:

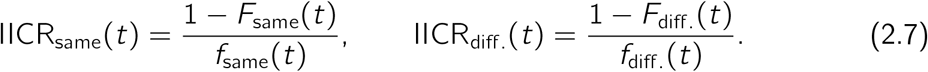

#### 2.2.3. The scaled IICR

Note that in the previous definitions (and in accordance with the definitions in §2.1), the functions in (2.5) receive the time *t* in units of 2*N* generations, and return values in units of *N* generations per coalescence, so these IICR functions are dimensionless in the sense that they operate in a *relative* frame of reference.

In order to get the IICR in an absolute frame of reference, which is useful for example when comparing it with PSMC inferences, we need to properly re-scale both the time and the IICR values themselves by a reference deme size *N* which specifies how many haploid genes correspond to a local deme size of 1. Given any positive value for *N*, the scaling would be carried out as follows:

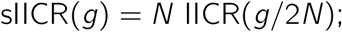

where sIICR(*g*) refers to the *scaled* IICR, and IICR(*t*) to the *unscaled* (dimensionless) one. Note that we use *g* for generations as the variable name for sIICR to further stress the difference. The returned values of the sIICR functions may be interpreted in units of *generations per coalescences*. The parameter space for the sIICR can be thought of as a simple one-dimensional addition to Φ_*γ,B*_:

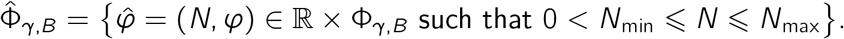

It is important to point out that, in contrast with the parameter space for the unscaled IICR (Φ_*γ,B*_) in which there is a one-to-one correspondence between a parameter tuple and the corresponding IICR curve, there are only 3*γ* + 3 independent degrees of freedom in 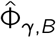, even though there are 3*γ* + 4 parameters. This notion is formalized by the following lemma.

##### Lemma 1.

*Given any* 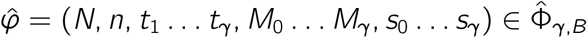, then the parameter tuple:

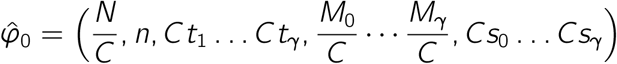

is such that:

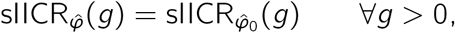

*where C is any rescaling factor for which the coordinates of* 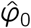 *are within the bounds B*.

A proof for this lemma can be found in §1 of the supplementary materials.

The implication of Lemma 1 is that when trying to infer all the parameters of 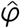 simultaneously, the only parameter for which we may get an absolute estimate is *n*, as the rest of them can only be distinguished up to an unknown re-scaling factor *C*. Note that this un-identifiability issue is different from the one identified in Mazet et al. (2016) regarding the inability to discriminate between panmictic and non-panmictic demographies with a single IICR. However, we stress here that in practice this is not necessarily an issue because it suffices to fix one of the model parameters (for instance, *s*_0_ = 1) to be able to uniquely map any sIICR curve to its parameters. In the case of constant size this is even less of an issue, since *all* deme sizes are fixed to *s_i_* = 1 and thus no further considerations are necessary.

### 2.3. Optimization framework: search algorithm and optimality criteria

In this section we describe the search algorithm used to explore the parameter space and the optimality criteria used to select the structured scenario that best explains a given IICR curve. We first assume that there is a known target IICR curve, either scaled or unscaled. This curve can originate from a PSMC analysis of a real or simulated genome, or it may be obtained after simulating *T*_2_ values or exact IICR curves (following the approach in §2.2.2) from an arbitrarily chosen demographic model. In this section we denote the target IICR exclusively with IICR_0_ for the sake of brevity, but the same constructions are valid for scaled IICRs. We also assume that the underlying coalescence times for these IICRs have distribution *F*_0_ and density *f*_0_.

Given a target IICR and a parameter space Φ_*γ,B*_, we want to find a parameter tuple *φ* in Φ_*γ,B*_ such that the exact IICR curve corresponding to the model defined by *φ* (henceforth denoted by IICR_*φ*_) approximates the target IICR as best as possible. We can express this more formally with the following optimization problem statement:

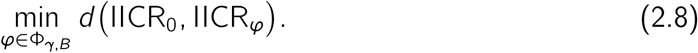

Thus, our tasks are reduced to finding an appropriate definition for the distance function *d*, and solving problem (2.8).

Regarding the distance *d*, a straightforward definition would be:

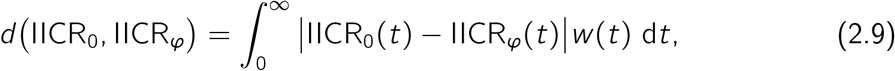

where *w*(*t*) is a weight function that should take into account the natural distribution of the information in an IICR. One reasonable solution for *w* is to take a quantity proportional to the density *f*_0_ of the coalescence rate because it ensures that the integral in (2.9) is finite, and also because it assigns more weight to the temporal periods where the target IICR is expected to be more accurate and reliable since more coalescences are likely to have happened.

We thus consider the family of weight functions:

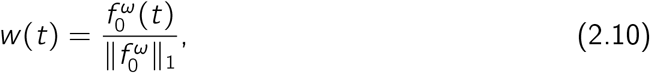

where ║·║_1_ is the *L*^1^-norm and *ω* > 0 is a *weight-shifting* parameter, with the purpose of dampening (if *ω* < 1) or exaggerating (if *ω* > 1) the effect of the weight *f*_0_. Unless otherwise noted, we take *ω* = 1, which corresponds in practice to giving more weight to recent periods of the IICR. The inference may thus ignore the most ancient parts of an IICR or PSMC curve as they are based on much fewer coalescent events.

In practice, we need to consider that all we know about IICR_0_ is a stepwise discretization over a bounded interval of time, so a numerical approximation of the distance (2.9) is required. This includes approximating the density *f*_0_ of the underlying *T*_2_ distribution. Given a division of time into *I* intervals {[*τ*_*j*−1_; *τ_j_*)} for 1 ⩽ *j* ⩽ *I*, where *τ*_0_ = 0 and *τ*_1_ < ∞, we can consider a discrete representation of IICR_0_ in the form of a collection of *I* values {*y_j_*} such that:

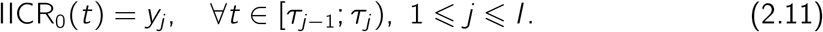

We can use this form to compute a numerical approximation for the integral in (2.9). For instance, a first degree approximation would yield:

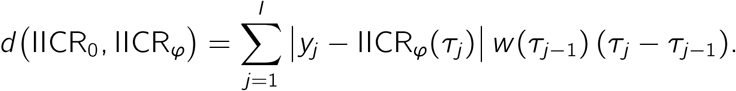

As for the values of *w*(*τ_j_*), notice that from (2.1) we have the identity:

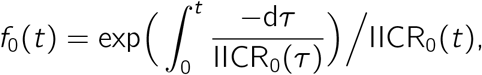

which, using the representation (2.11), can be discretized into:

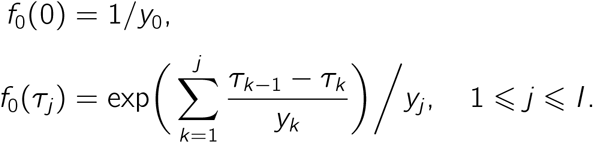

We then have that for any given *ω, w*(*τ_j_*) can be expressed as:

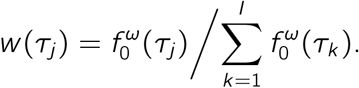

An alternative option for the definition of *d* in (2.8) could be one that takes into account the ultimate visual nature of the curve fitting task. Assuming that the points {*τ_j_*} are somewhat log-distributed and that they will be used for visualization purposes in a horizontally log-scaled plot like Figure 1, then the definition:

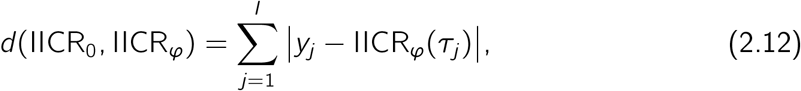

captures the perceived visual difference between the plots of the two curves. We distinguish distance (2.9) from (2.12) by denoting them *d_ω_* and *d_visual_* respectively. We keep both definitions because we found that the weighted family of distances generally performs better than the visual distance under certain validation tests, but also that the *d_visual_* distance might be used to choose the optimal weight parameter in *d_ω_* (see §4).

Regarding the optimization problem (2.8) itself, we use the *Differential Evolution* method (Storn and Price, 1997). As a genetic meta-heuristics, this algorithm maintains and evolves (using mutation and recombination parameters) a population of solutions iteratively. As a global optimization algorithm, it features mechanisms for escaping local *optima* of the parameter space. In §2 of the supplementary materials we explore the potential effects on the inference results of tuning some of the parameters provided by this algorithms implementation. For our validations, we run the method multiple times and we refer to these repetitions as *rounds*. We set a maximum number of allowed rounds, as well as a tolerance *ε* for the distance which controls the minimum number of rounds.

### 2.4. Validation framework

For the purpose of validation we applied our inferential method to target IICRs generated under piecewise stationary *n*-island models of increasing complexity (i.e., number of components) and with known parameter values (*N, n, t_i_, M_i_*) and then compared the inferred parameter values to those actually used.

In what follows we present various ways of generating random demographic scenarios and then computing appropriate IICR curves from them for use in validation.

#### 2.4.1. Sampling the parameter space

Given a parameter space Φ_*γ,B*_ (we only discuss the unscaled case here for brevity, but the same principles apply to a scaled parameter space), we sample demographic scenarios from which we compute the corresponding IICRs. We used two sampling strategies which call continuous and discrete sampling.

##### Continuous sampling

Assuming that we want to realise *L* independent tests, this sampling strategy consists in using uniform or log-uniform distributions for each of the 3*γ* + 3 random variables:

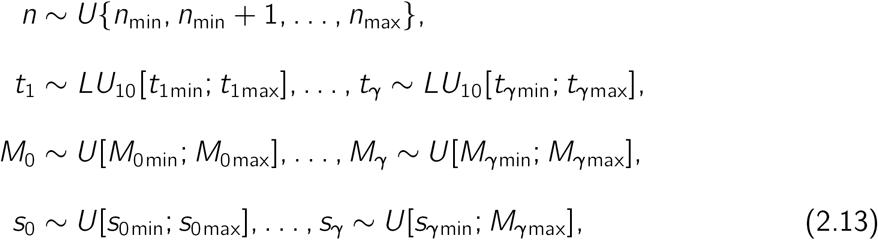

where *U* denotes a uniform distribution (discrete in the case of *n* and continuous for the rest) and *LU*_10_ denotes a log-uniform distribution of base 10. This distribution is used for sampling the times of changes in a logarithmic space in order to take into account the natural distribution of information in an IICR.

In order to construct a valid demographic scenario from these realisations, we need to enforce that the event times are in increasing order. Consider then for every 1 ⩽ *j* ⩽ *L* the sorted times 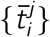 such that:

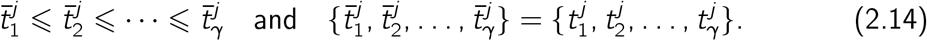

With this, we can define the *L* sampled scenarios as those corresponding to the tuples:

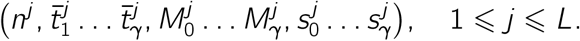

This sampling strategy provides a comprehensive coverage of the parameter space and allows for an easy visualization for the sampled-versus-inferred scenarios (see §3.1). It also makes it very unlikely to sample exactly the same parameter values twice or to sample exactly the same *M_i_* values in two consecutive components. However, it sometimes produces demographic scenarios in which consecutive *t_i_* and/or *M_i_* values may be close to each other, and thus difficult to distinguish. This makes it thus harder on our inferential framework compared to cases where we would chose contrasted scenarios with clearly separated events with major changes in migration rates. In other words, our inferential method was sometimes trying to infer parameters in the case of extremely difficult scenarios as we show below.

In §3.1 we show the results obtained using this sampling method with *L* = 400 scenarios and the following values for the sampling bounds and the slightly larger inference bounds:

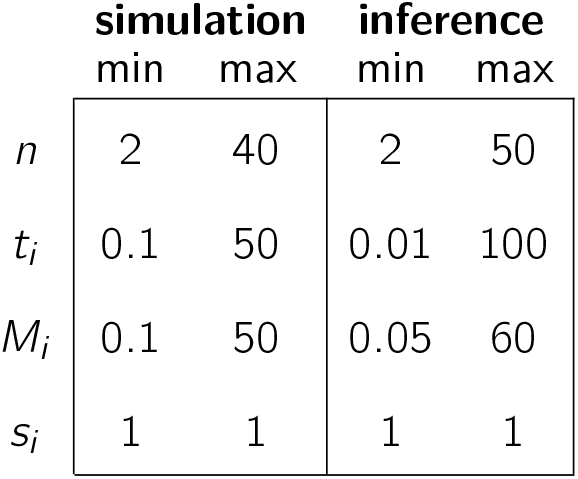

|  | simulation |  | inference |  |
| --- | --- | --- | --- | --- |
|  | min | max | min | max |
| $n$ | 2 | 40 | 2 | 50 |
| $t_i$ | 0.1 | 50 | 0.01 | 100 |
| $M_i$ | 0.1 | 50 | 0.05 | 60 |
| $s_i$ | 1 | 1 | 1 | 1 |

##### Discrete sampling

Here we sampled *L* = 100 independent scenarios from the same parameter space, but using the following set of predefined values:

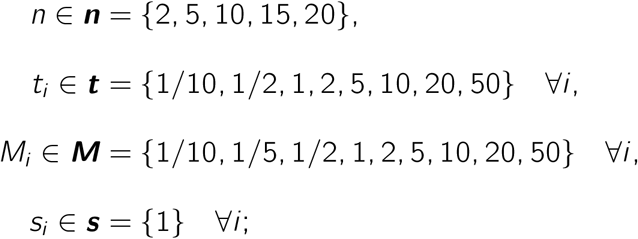

The inference process was, however, done within the continuous space. For instance, under this validation scheme (see §3.2) we only simulated data with 2, 5,10,15, 20 islands but the inference process always allowed *n* to take any value between 2 and 50. The choice of the *L* independent simulated data sets was done using the following procedure. We first considered the following cartesian product of dimension 3γ + 3:

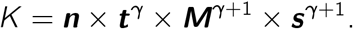

and then uniformly drew *L* tuples from *K* without substitution. We then sorted the sampled event times as in (2.14), hence obtaining a set of *L* demographic scenarios. Note that it is then straightforward to include additional filtering rules when sampling from *K* to discard undesired scenarios, such as one containing two consecutive components with the same migration rate or two consecutive values from ***t***. In our simulations, we drew randomly (without replacement) from the set *K*, rejecting scenarios with identical *M_i_* values in two consecutive components, until we reached *L* accepted scenarios.

#### 2.4.2. The three types of target IICRs

We aim to generate target IICRs from a known demographic model in order to compare the model inferred from this IICR curve to the correct one (see Figure 3). Here we explore three different types of target IICRs (see Figure 4) that could be obtained given a scenario *φ* ∈ Φ_*γ,B*_. Note that all IICRs are discretized so as to be comparable to PSMC plots (see equation 2.11).

**Figure 3:**
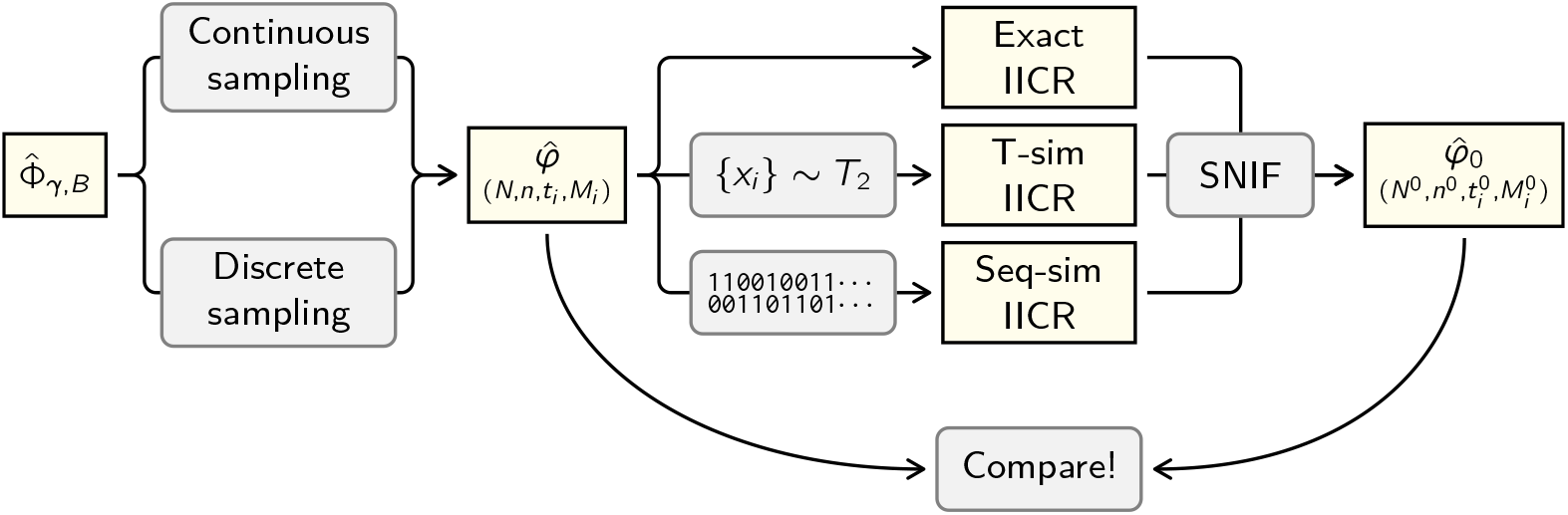
Flowchart of the validation procedures. Starting from a parameter space 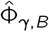 we use one of two sampling methods (§2.4.1) to generate a demographic history 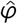 defined (for the scaled case) by the parameters (*N, n, t_i_, M_i_*). We then compute the IICR of that demographic history using one of three methods (§2.4.2) to obtain the target IICR. After that, we run the inference algorithm on this target IICR curve (using wider bounds than those in B) to obtain an estimated (or inferred) demographic history 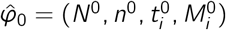, which we then compare to the known 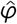 in order to assess the accuracy of the inference methodology (§3).

**Figure 4:**
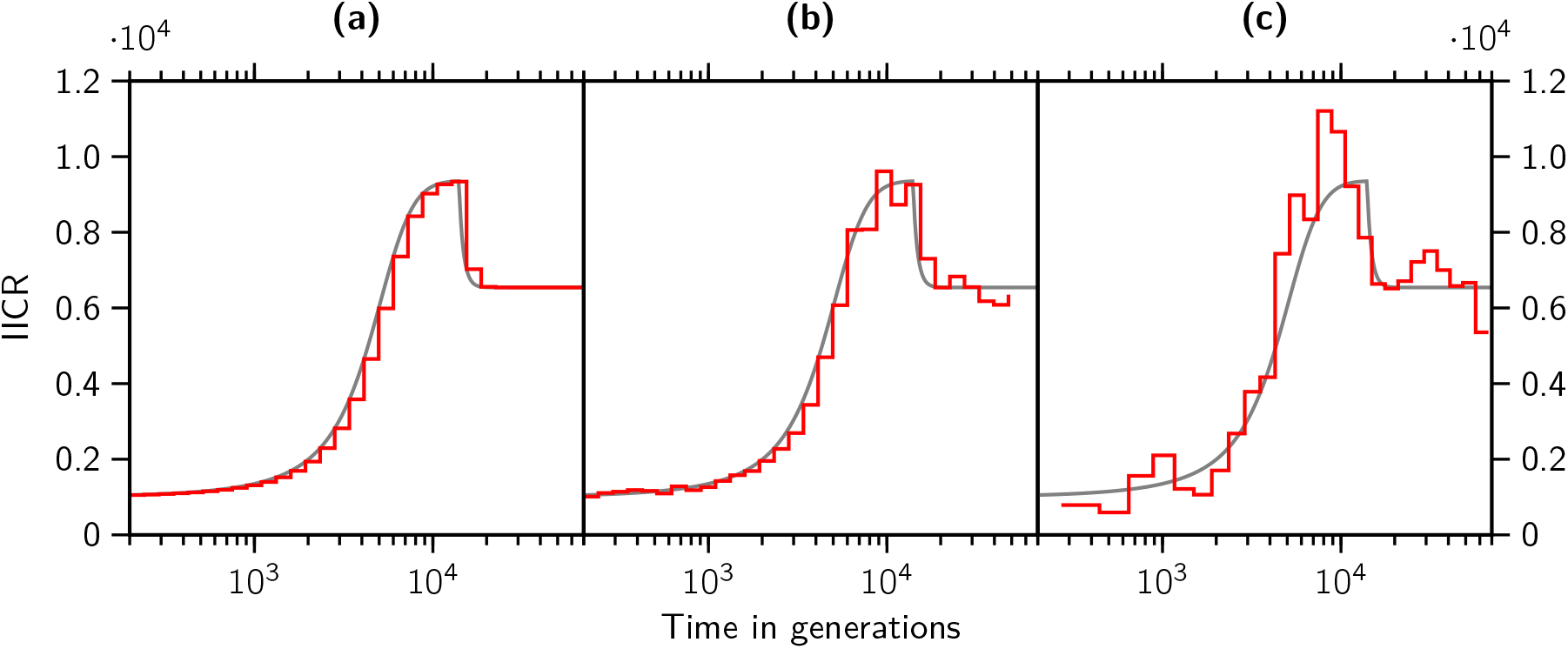
The three ways of obtaining a target IICR. The panels represent different types of discretised target IICRs for the same two-components demographic history with *N* = 10^3^, *n* = 4, *t*_1_ =7, (*M_i_*) = (0.5,1.0) and (*s_i_*) = (1.0,1.0). **(a)** Exact IICR, computed as per §2.2.2 and discretized to 32 intervals. In gray we show a smoother discretization and keep it in the other panels for reference. This is obtained using the approach of Rodríguez et al. (2018). **(b)** T-sim IICR, obtained by simulating 2 × 10^4^ *T*_2_ samples using the `ms` command `ms 2 20000 -T -L -I 4 2 0 0 0 0.5 -eM 3.5 1 -eN 3.5 1` and later scaling the IICR using the value of *N*. This is the approach used in Chikhi et al. (2018) **(c)** Seq-sim IICR, obtained by running PSMC on a genome simulated using the ms command `ms 2 100 -t 100 -r 20 2000000 -p 8 -I 4 2 0 0 0 0.5 -eM 3.5 1 -eN 3.5 1` and later scaling with mutation rate *μ* = 1.25 × 10^-8^. This is obtained by running the PSMC method of Li and Durbin (2011)

Validating our approach on PSMC plots across the parameter space described above would be extremely time consuming as it would require simulating genomes and then running the PSMC method (or other related methods) on these genomes before applying our approach. We stress here that we will only run the PSMC method in the case of the scenarios inferred for the human data so as to integrate the uncertainty due to the PSMC inferential process. This is a crucial issue but here our aim is not to test the PSMC or other inferential methods, but rather build a framework based on the IICR to identify models with changes in connectivity. To clarify this we explain below the different types of IICR that could be computed given a scenario *φ* ∈ Φ_*γ,B*_.

##### Exact IICR

We saw in §2.2.2 how to compute the IICR for any *n*-island model at any time value *t*, but to generate input data we need a discretization as in (2.11), so considering that we take a log-distributed sample of size *I* in the interval [*t*_min_, *t*_max_], we end up with a suitable IICR_0_. Note that even though this IICR has been discretized, its values are exact within machine precision, so it is still an artificial product compared to real data.

For the validations using the exact IICR in §3.1 we chose for the distance tolerance between a target and an inferred IICR a value of *ε* = 10^-10^ for the unscaled IICRs and an equivalent value of *ε* = 10^-7^ for the scaled IICR (since the simulated *N* was always 1000). It should be noted that this value of e is quite small even for double-precision floating-point arithmetic, and thus is only a reasonable choice for validation using exact IICRs (i.e., those where the distance could theoretically be zero).

##### T-sim IICR

The T-sim IICR is obtained by simulating a finite collection of *T*_2_ realizations using ms and then building an empirical IICR as in Mazet et al. (2016), using the Kaplan–Meier estimator (Kaplan and Meier, 1958), with log-distributed times. We stress that ms scales time in units of 4*N* generations whereas our models use a scale of 2*N* generations (see the example in Figure 4), so this must be kept in mind when writing ms commands.

##### Seq-sim IICR

In order to closely follow the pipeline of real genomic data, we simulate genomic sequences with ms and then apply the PSMC method for obtaining the IICR to be used by the inference method. Since simulating genomes and performing PSMC analyses is significantly more time consuming than the other two methods used for obtaining a simulated IICR, we limited ourselves to validating the Seq-sim IICR for the human PSMC based scenarios that we obtain after performing the demographic inference described in §2.5.

### 2.5. Application to humans

We applied our method to the human genomes published in the great apes study by Prado-Martinez et al. (2013). Namely, we used the PSMC files of five sampled humans identified as Dai, French, Karitianan, Sardinian and Yoruban (see Figure 5). For each human PSMC curve we performed demographic inference independently within the following bounds:

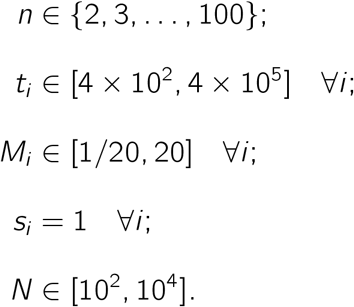

**Figure 5:**
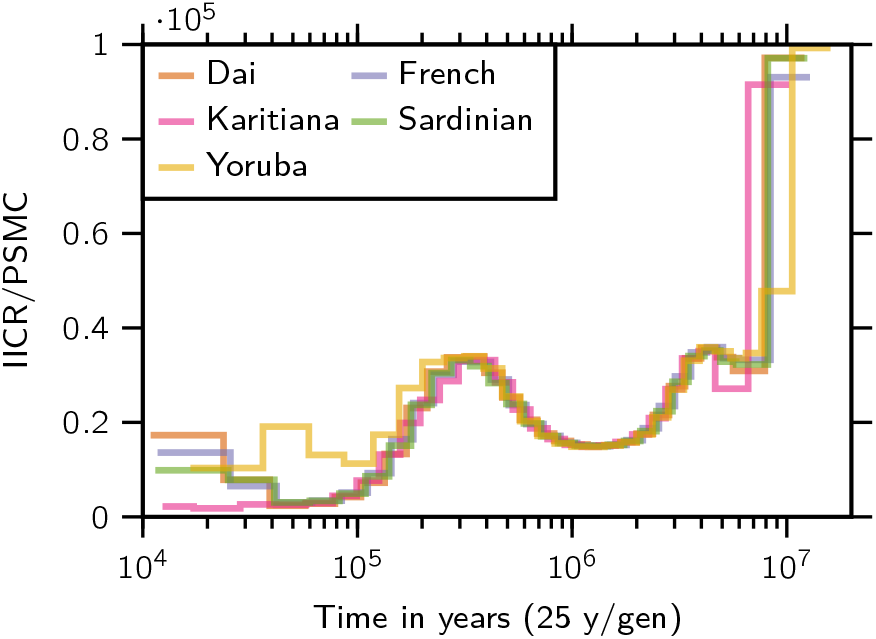
Five human PSMC plots. All five PSMC curves were obtained from the study of Prado-Martinez et al. (2013). The scaling was done as recommended by Li and Durbin (2011) (see equation 2.15 and surrounding explanation). No further processing was necessary since .*psmc* files contain the final result of the effective size or IICR estimate.

The bounds for the *t_i_* are specified in generations, so given a generation time of 25 years, we effectively allowed for the inference of demographic events between 10 thousand and 10 million years ago. Regarding the number of components, we choose *c* ∈ {2,3,4, 5}; i.e., between one and four demographic events in agreement with Mazet et al. (2016) who suggested that a minimum of three events were necessary to explain the two humps, and in agreement with our validation simulations which suggest that inference above five components is difficult.

Some of the analyzed IICR curves exhibit an increase in effective size in the recent past, which could be due to a genuine population growth as noted by Mazet et al. (2016). Given that we choose to specifically rule out changes in deme sizes, we account for this fact by running every inference a second time, ignoring this period of possible recent expansion. This is accomplished using an option that allows to limit the interval where the distance function is computed. In this case, we restricted both this range and the bounds for the *t_i_* to be between 50 thousand and 10 million years ago, thus ignoring any population size change that may have happened in the last 50,000 years. Note that this option is also useful to ignore very ancient sections of the PSMC plots which may be difficult to trust.

Since these inferences are based on real human PSMCs which are unlikely to have been generated by an *n*-island model—as opposed to the inferences described in §2.4 where the underlying model was in fact a piecewise stationary *n*-island coalescent—the default value used for *ω* may not be the most appropriate and we thus performed inferences with *ω* ∈ {1, 0.5, 0.2}. Decreasing values of ω give increasing weight to the most ancient part of the PSMC (see the weighted distances (2.10)). The resulting inferred demographic scenarios are shown in section §3.3.

As a way of validating the inference process using PSMC outputs, we generated 10 Seq-sim IICRs corresponding to the inferred demographic scenarios for the French, Karitianan and Yoruban individuals (the inferred histories for the Dai and Sardinian are similar enough to the other three that they don’t warrant a separate analysis). For each one of these scenarios we simulated nreps = 30 chromosomes of length *L* = 10^8^ base pairs, using the effective size *N* inferred by the method, a per-base per-generation mutation rate of *μ* = 1.25 × 10^-8^ and a scaled recombination rate of *ρ* = *θ*/5 (as in Li and Durbin (2011)), then we run the ms command with *θ* = 4*μLN* using:

~~~
ms 2 nreps -t *θ* -r *ρ* L -p 8 -I …
~~~

where the rest of the command follows according to the inferred demography (see Figure 4 for a reference). After that we prepared a .*psmcfa* file as input for PSMC, always choosing a bin size of *s* = 100. Then we ran the PSMC with the command:

~~~
psmc -N25 -t15 -r5 -p “4+25*2+4+6” …
~~~

following Li and Durbin (2011) on human data. We then applied the final step required to obtain a scaled IICR by applying the following scaling to the data in the resulting *.psmc* file:

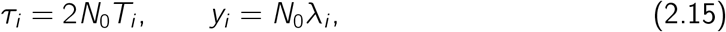

where the *T_i_* and *λ_i_* are the time and size data from the *.psmc* file, and *N*_0_ corresponds to the population size at time *T* = 0 in the panmictic model underlying the PSMC, and is computed as *N*_0_ = *θ*_0_/(4*μs*), where *θ*_0_ is the one given in the *.psmc* file and s is the bin size for preparing the input to PSMC (in our tests we took s = 100). Finally, we used these PSMC curves as targets to determine whether we could indeed infer the parameters used for such complex scenarios.

We also applied our method to genomic data simulated under the scenarios used to describe recent human evolutionary history by Gutenkunst et al. (2009); Noskova et al. (2019). Here, we thus ask the following two questions: if human evolution were indeed closer to such splitting models, would our method infer again an *n*-island model with similar parameters to those inferred from the humans PSMCs? additionally, do these models generate IICR plots that are similar to the human PSMCs?

## 3. Results

In this section we show the results of validating the inference method using target IICRs from known demographic histories; the application of the method to real human data; and the comparison of the obtained results with previously published demographic histories for humans. When performing validation, both the source and the inferred demographic models are known, which allows us to assess the accuracy of the inference *in the parameter space*, i.e., we can compare the values of the inferred parameters to those of the simulated parameters. This is not possible when performing an analysis of real genomic data since the model is unknown. Therefore, we always include comparisons of inferred and observed of IICR curves since for real data, what is observed is an estimated IICR (possibly in the form of a PSMC curve), and as such this is the type of visualization that is always possible to do. The results of the validations are presented in Figures 6 to 8 in the main manuscript and Figures S3 to S29 in the supplementary materials. Another set of figures (Figures 9 to 13 and Figures S30 to S36) present the results of the application to human data.

**Figure 6:**
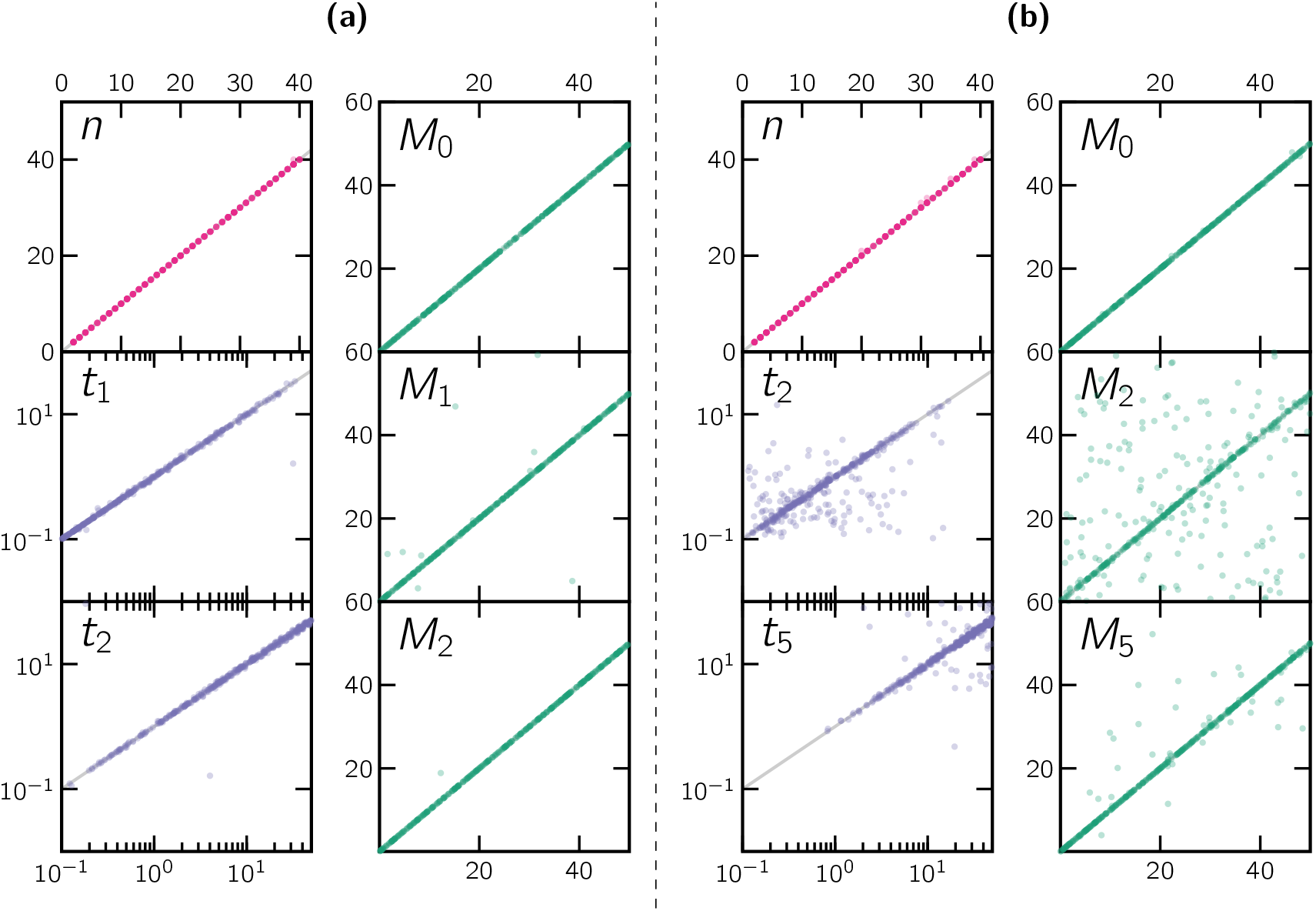
Scatter plots of simulated and inferred parameters. Panel **(a)** corresponds to scenarios with c = 3 components, and **(b)** to scenarios with *c* = 6 components. The different sub-panels represent the simulated (horizontal axis) versus inferred (vertical axis) parameter values for all the parameters (or a representative selection of parameters in the case of panel **(b)**) of *L* = 400 unscaled simulated scenarios.

**Figure 7:**
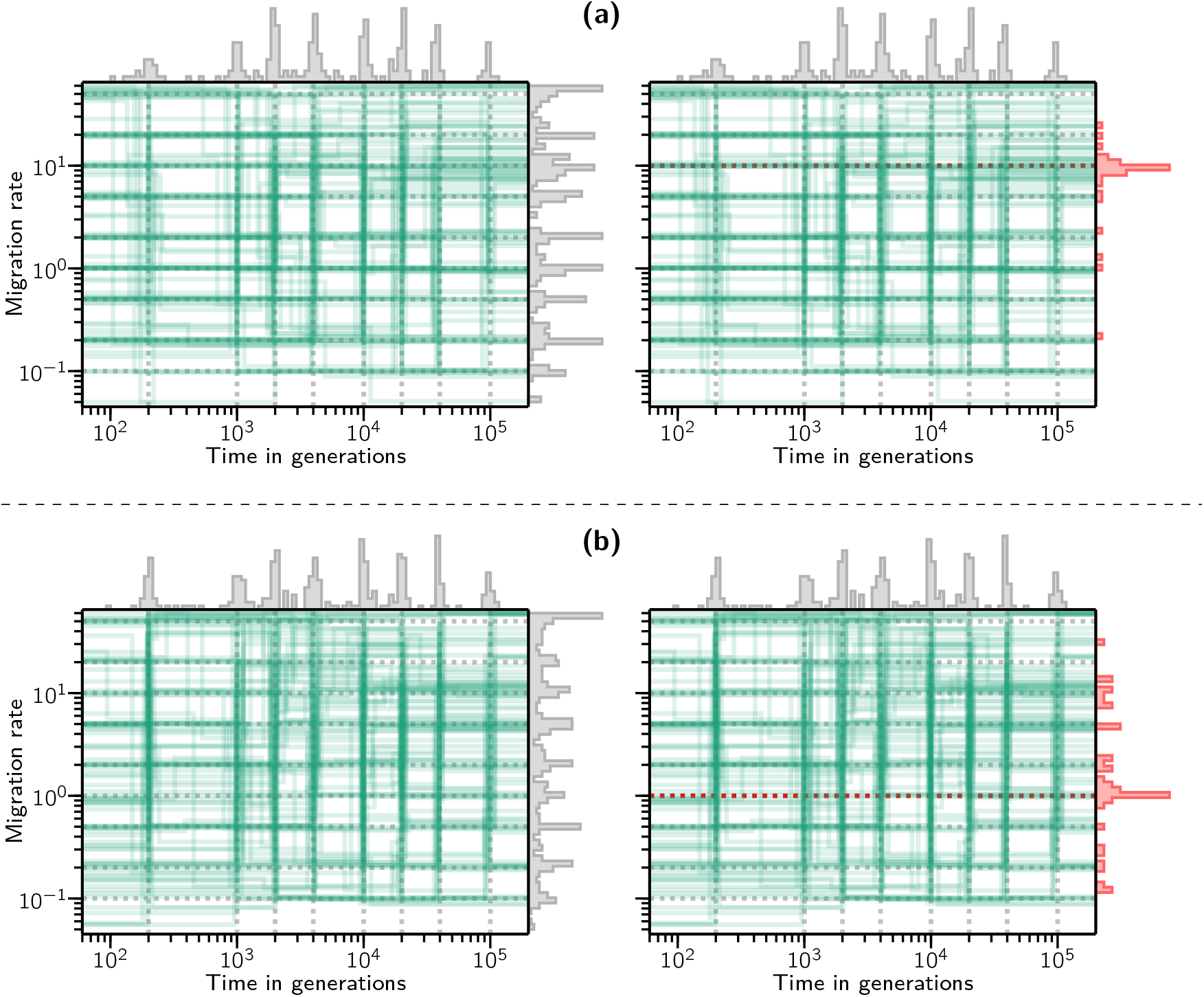
Connectivity graphs of 100 independently inferred histories obtained by sampling for each scenario from the values indicated by the dotted lines. **(a)** scenarios with *c* = 3 components. **(b)** scenarios with *c* = 4 components. The right sub-panels show a side histogram with only the inferred migration rates for those components with a specific simulated migration rate (10 for **(a)** and 1 for **(b)**).

**Figure 8:**
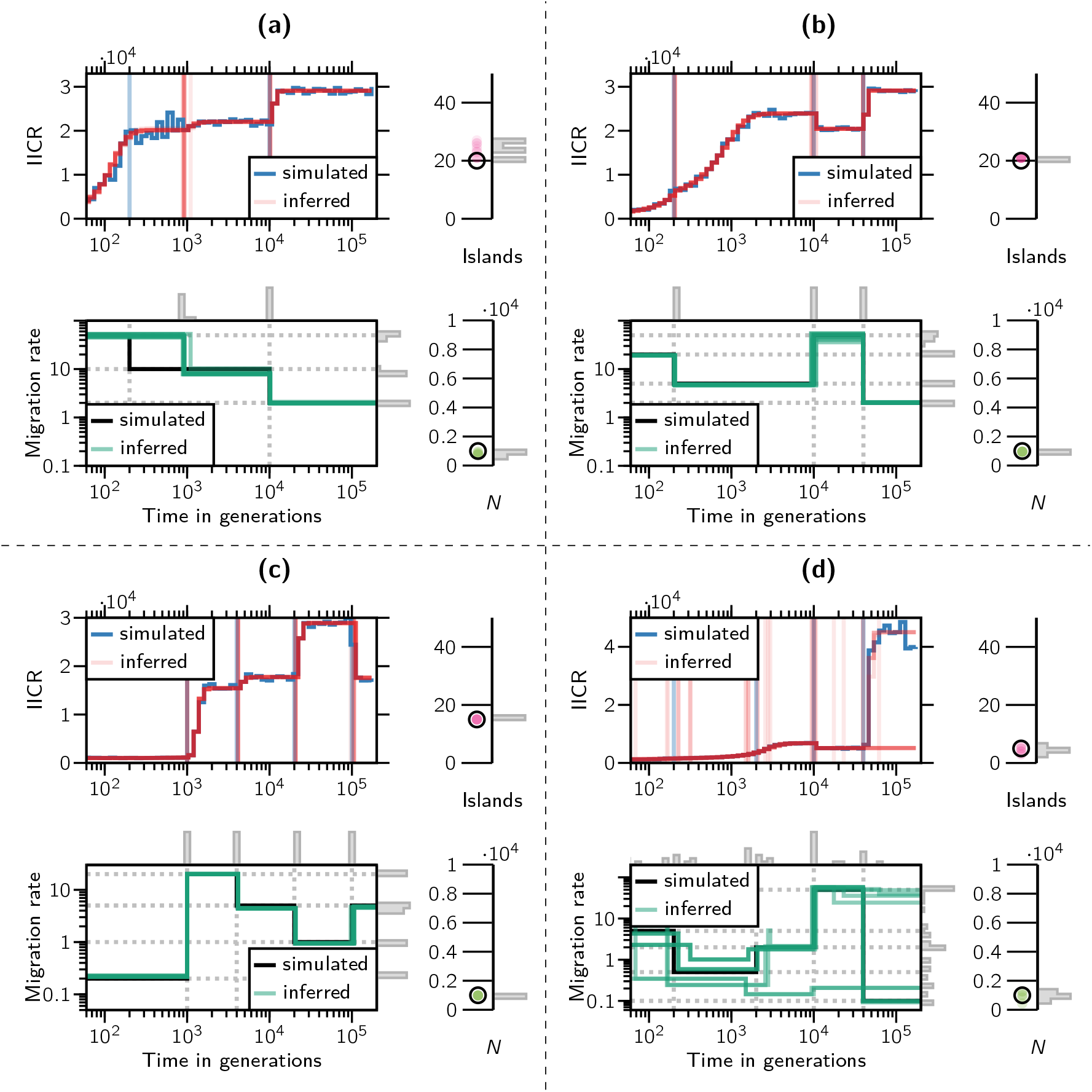
Simulated and inferred IICR plots, connectivity graphs, *N* and *n*. The four panels correspond to four different scenarios. **(a)** a *c* = 3 components scenario **(b)** a *c* = 4 components scenario **(c)** a *c* = 5 components scenario **(d)** *a c* = 5 components scenario. The left part of each panel represents the target and inferred IICRs (top), and the connectivity graphs (down). The right half of each panel shows the simulated and inferred values for *n* (top) and *N* (down). In each IICR graph, the ragged blue line represents the target IICR whereas the red lines represent 10 independently inferred IICRs. The vertical blue and red lines represent the simulated and inferred *t_i_* values, respectively. In the connectivity graphs, the black and green lines represent the simulated and inferred connectivity scenarios, respectively. The simulated *n* and *N* values are represented by black open circles whereas the inferred values for the corresponding parameters are represented by red and green full circles and by grey histogram bars.

**Figure 9:**
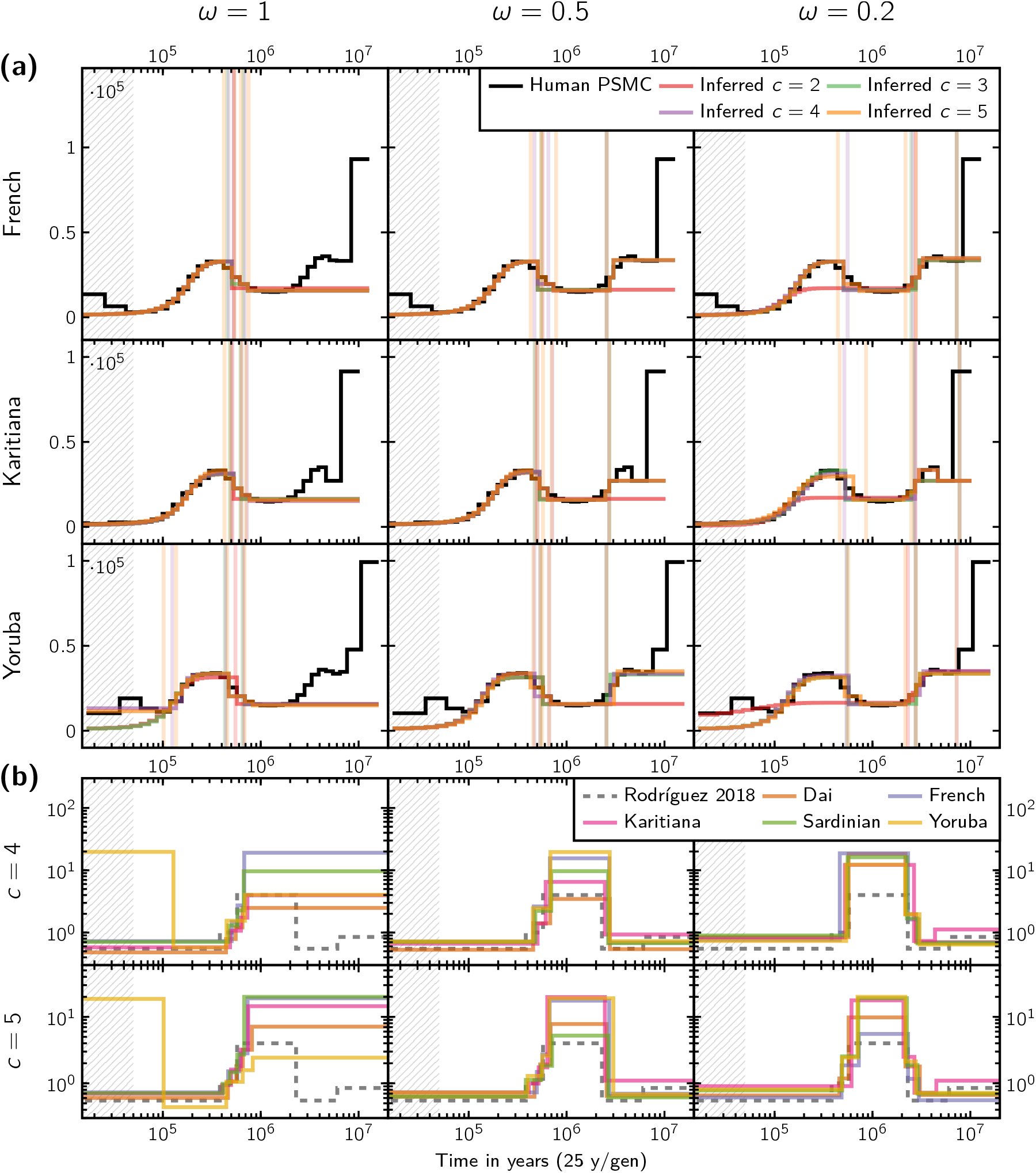
Results of performing demographic inference on three representative human PSMC curves. **(a)** shows the various IICR plots inferred for the different populations, numbers of components c and weight parameters *ω* used, together with the target IICR curves (or PSMC plots) on which these estimations are based. **(b)** shows the connectivity graphs for the same set of inferred scenario. As a reference point, the connectivity graph of the scenario proposed in Rodríguez et al. (2018) is also shown. The vertical axes represent migration rates (*M*).

**Figure 10:**
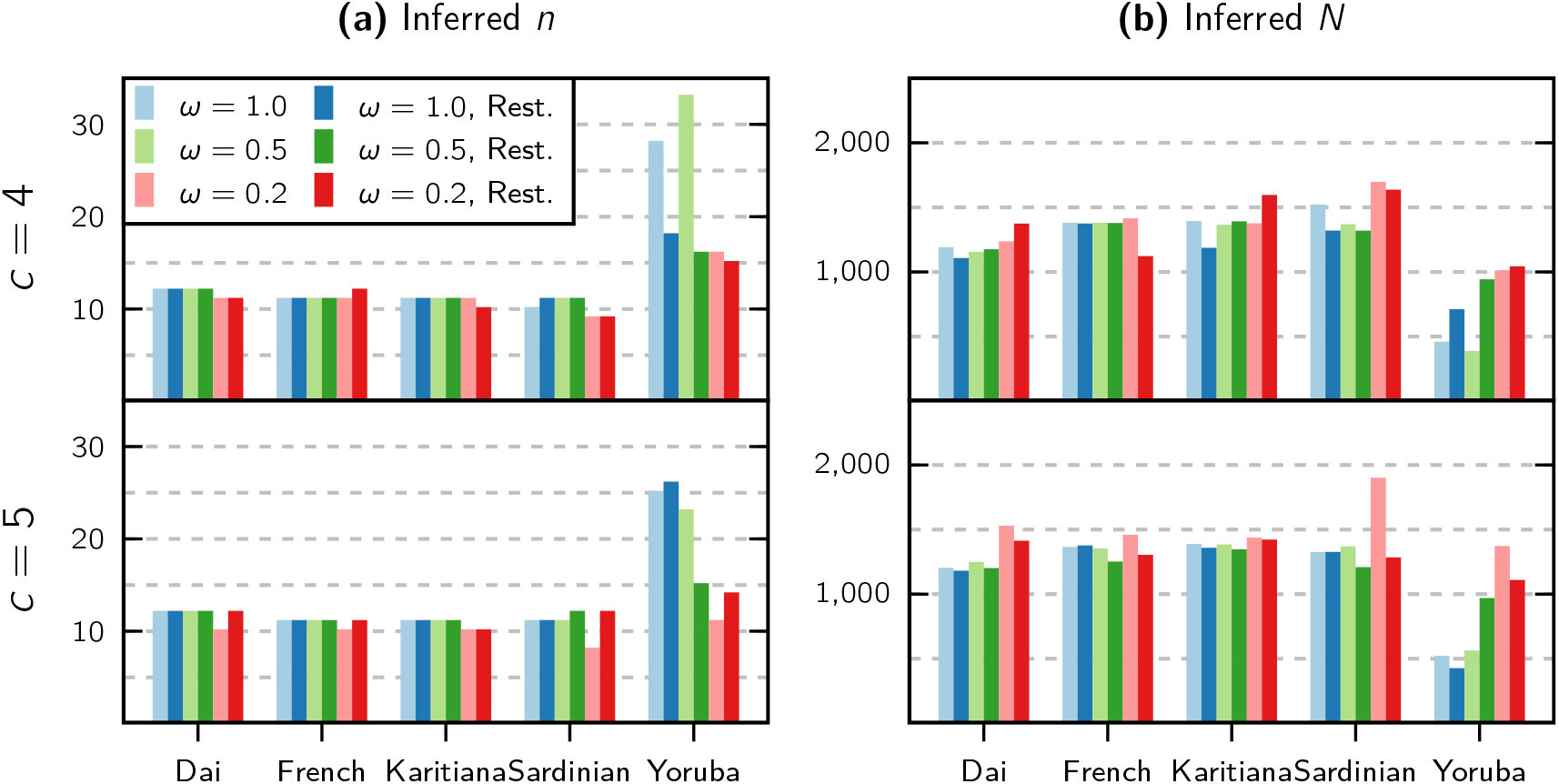
**(a)** Inferred number of islands (n) and **(b)** inferred reference sizes (N) for each human population and each used combination of the weight parameter *ω*, number of components (only 4 and 5 are shown here), and with or without the option of ignoring recent population expansion.

**Figure 11:**
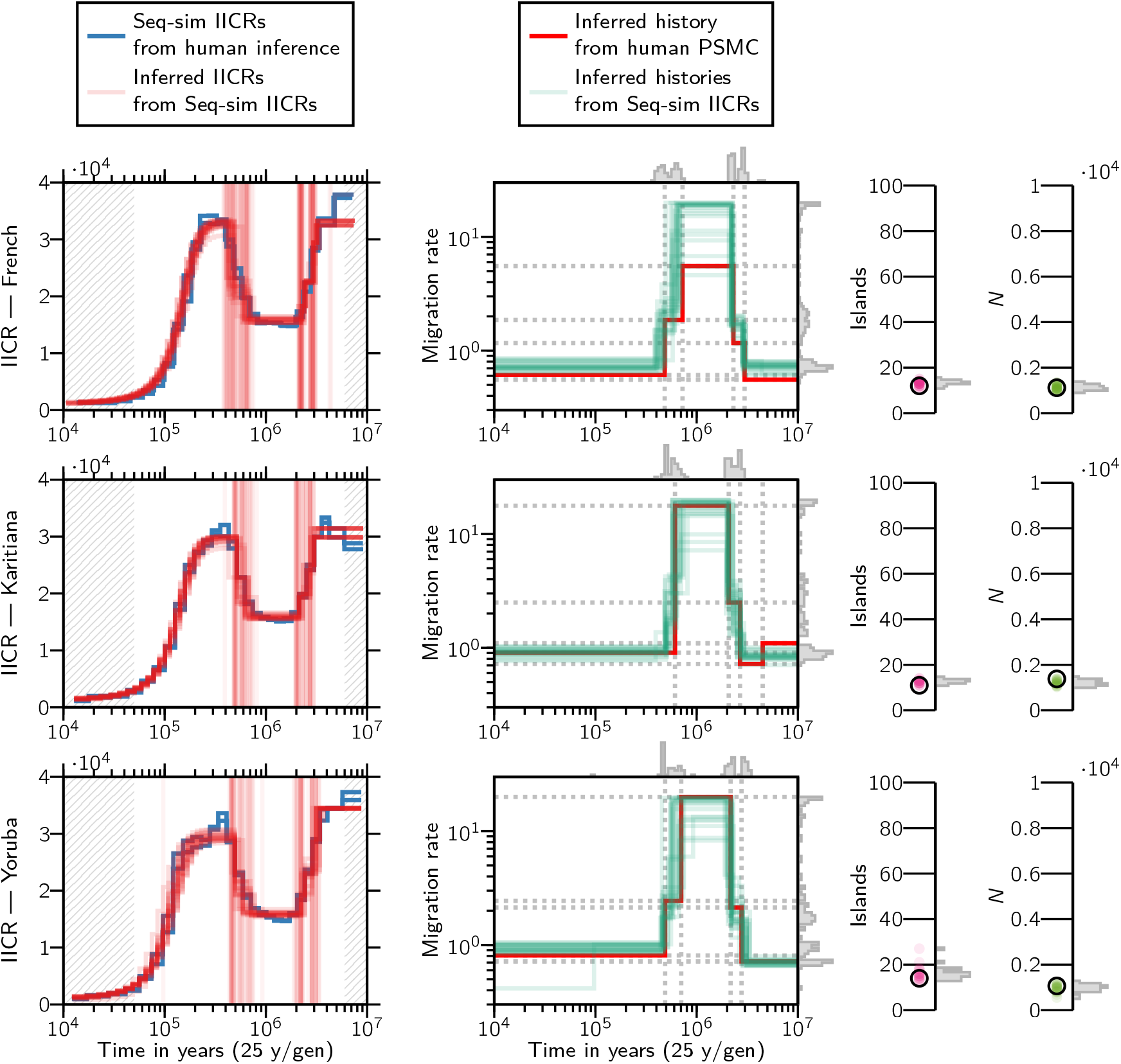
Results of the validation using Seq-sim IICRs. The left panels show two independently simulated seq-sim IICRs (obtained using the demographic scenario inferred with *c* = 5 and *ω* = 0.2 for each of the indicated human population) alongside 10 independent inferred IICRs. The rest of the panels show the connectivity graphs, number of islands, and local deme size of these seq-sim IICRs and their corresponding inferences.

**Figure 12:**
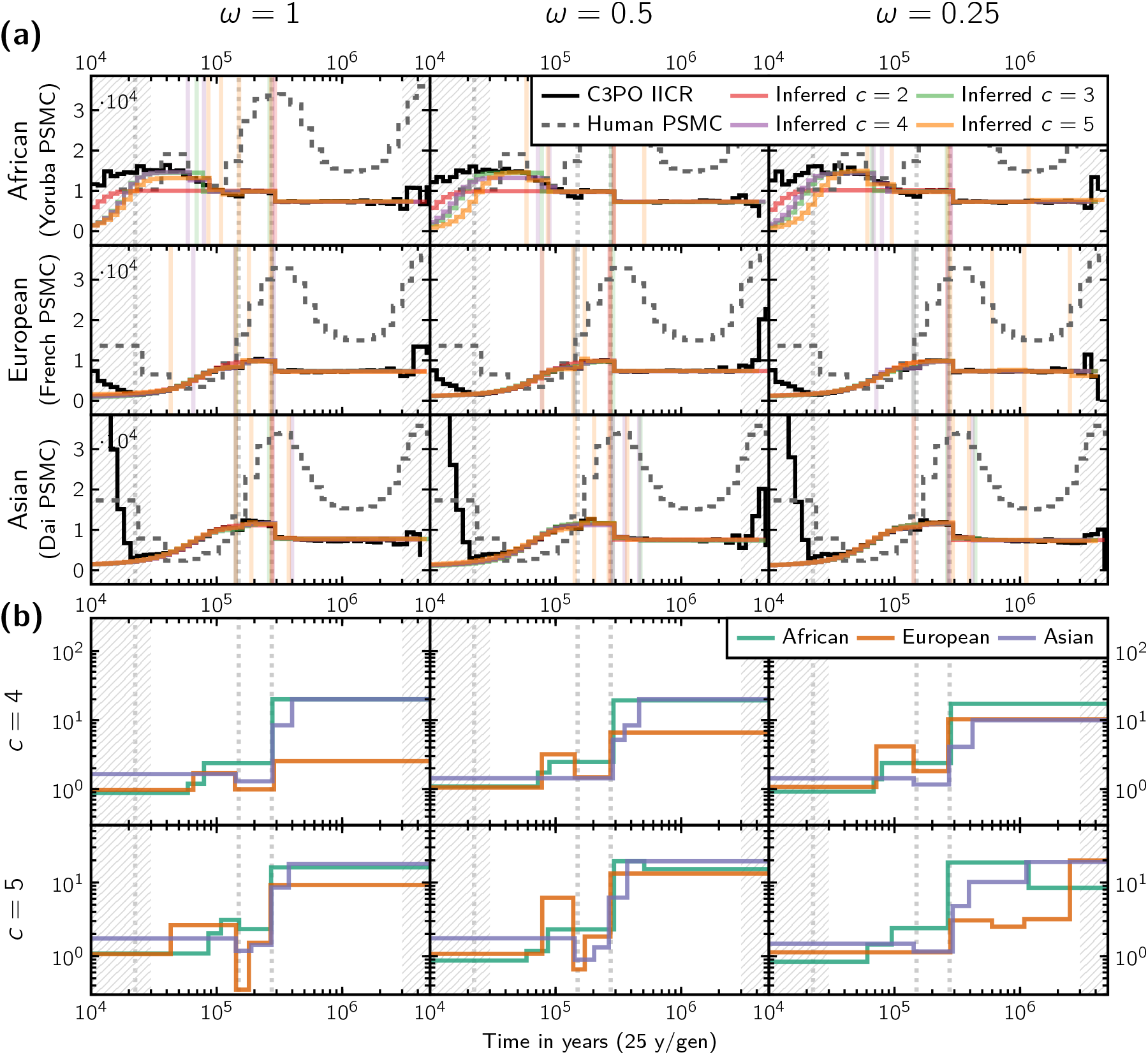
Application of our inference method to a generally-accepted tree-like human demographic scenario with three modern populations. **(a)** IICR plots showing the resulting IICR curves for this model and the inferred IICR curves obtained with our method (where the recent period of human expansion was ignored) for varying number of components c and weight parameters ω. For reference purposes, we also show the real PSMC curve of a human individual for each of the corresponding C3PO populations. **(b)** Connectivity graphs of the inferred scenarios.

**Figure 13:**
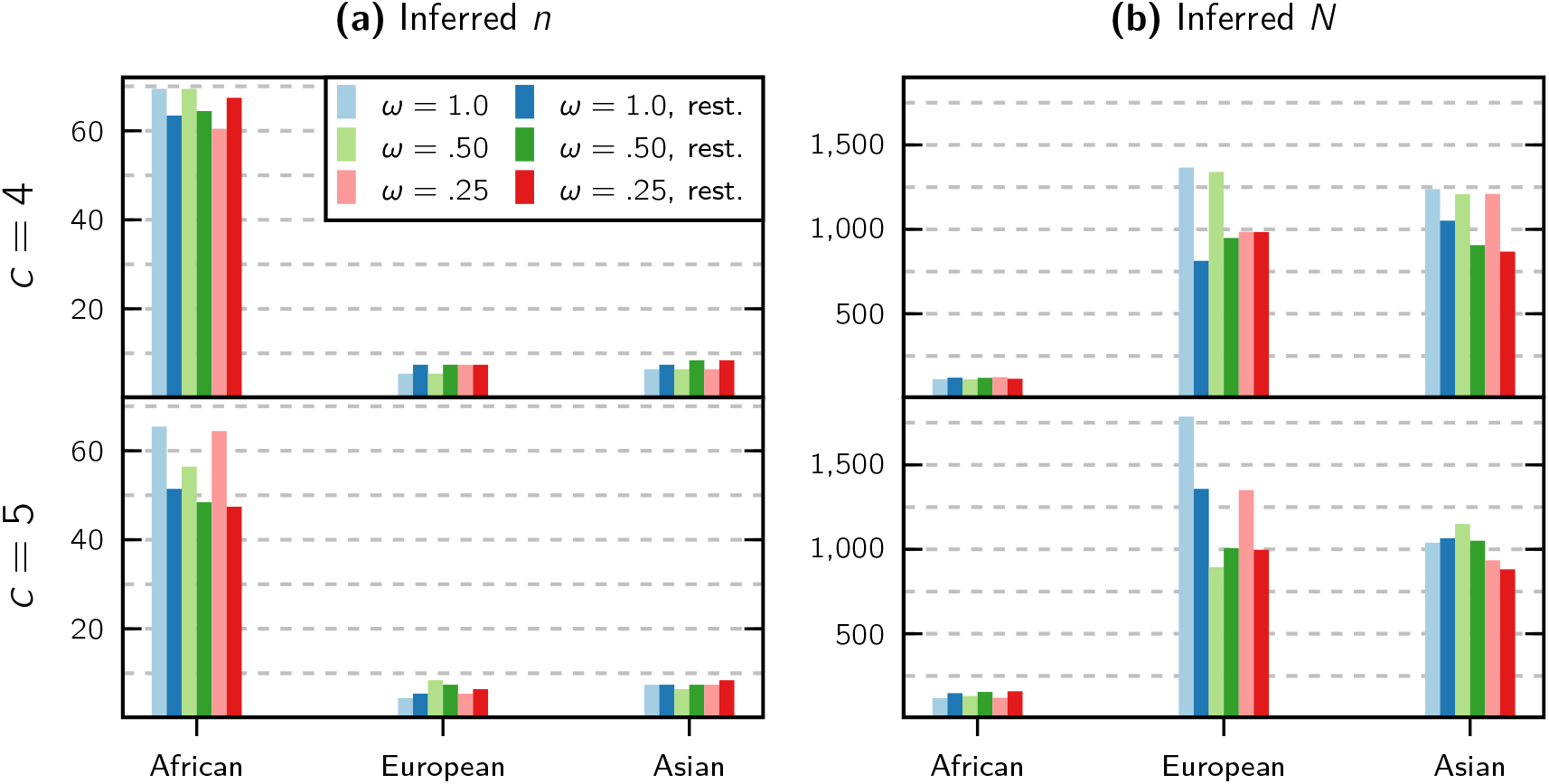
Application of our inference method to a generally-accepted tree-like human demographic scenario with three modern populations. **(a)** Inferred number of islands for each modern population. **(b)** Inferred local size of each island.

### 3.1. Validation using exact target IICRs

A first set of figures (Figure 6 and Figures S3 to S14) represents the simulated and inferred parameter values on the horizontal and vertical axes, respectively, using the continuous sampling strategy. As explained in §2.4.1, the range of possible values in the inference process was always wider than the range used for the simulated values. For instance, we simulated models with a number of islands varying between *n* = 2 and *n* = 40, but the search space for the inferred number of islands varied between *n* = 2 and *n* = 50. Altogether these figures always show a diagonal line corresponding to *y* = *x*, hence suggesting that the inferred parameter was identical or very close to the simulated parameters. This is particularly obvious for all the parameters corresponding to scenarios with up to four components. For instance, Figure 6a shows the results for a model with three components, in which there is a nearly perfect match between the simulated and inferred values for the three migration rates (*M*_0_, *M*_1_, *M*_2_) and the times at which migration rates changed (*t*_1_, *t*_2_). For five- and six-component scenarios the results are still nearly perfect for most of the simulated scenarios but we observe an increasing number of cases (i.e., simulated scenarios) where the parameters are poorly estimated, with the exception of *n*, *M*_0_ (and *N* for scenarios with scaled IICRs) which are almost always well estimated also in such cases. These cases can be identified by the dots scattered in the different panels. They start to appear in scenarios with three components, but their number grows with the number of components.

These poorly estimated scenarios are surprising given the near perfect estimation obtained for most parameter combinations. This is particularly striking because these dots do not seem to be distributed in any clear area of the parameter space. We see at least two possible and non-exclusive interpretations for this result. One is that the search algorithm had not yet *converged* when the maximum number of rounds was reached.

The maximum number of rounds was set to 500 in all simulations because we had found that less than 50 rounds were more than enough in the first tests carried out with one or two components. The search algorithm might however need more than 500 rounds to reach the optimal solution for scenarios of increasing complexity. We thus asked whether the maximum the number of rounds had been reached in the scenarios analysed and whether the proportion of scenarios with 500 rounds increased with the number of components. We found indeed that the proportion of simulations for which that maximum was reached increased with the number of components. For instance, all five- and six-component scenarios stopped their parameter search at 500 rounds, hence suggesting that at least some had not yet reached an optimum solution. For the cases with one- and two-component scenarios, all 800 independent simulations reached convergence in less than 150 rounds (see Figure S1). Again, the choice of the tolerance *ε* plays a role in these results, and selecting larger tolerances will tend to produce earlier convergence in general, but not necessarily better results.

As a test we randomly identified a couple of scenarios with six components that had bad estimates and re-ran the algorithm with 5000 rounds. We found that the distance value consistently decreases with more rounds (see Figure S2), but the inferred parameter values may not converge to the simulated ones because with more components there is a higher probability that two consecutive simulated *M_i_* values are very close, thus making the corresponding event time challenging to infer. Likewise, some simulated components may have a short duration that don’t leave a significant mark on the IICR curve, thus leading them to be “skipped”. We refer to this issue as component miss-identification, which could lead to a particular estimated parameter to be plotted in the wrong panel. For instance, the method may miss the first change in migration rate at *t*_1_ and identify the second change in migration at *t*_2_. In such a case the method will assign the inferred *t*_2_ value to the set of inferred *t*_1_ values and plot it in the *t*_1_ panel. This wrongly assigned *t*_2_ value will thus appear away from the diagonal in the t1 panel. Such miss-assignment cases for one parameter will also have consequences for the *M_i_* plots, and thus will generate several miss-assignments across panels. They are also expected to increase in frequency as the number of components increases and as the t, values become closer to each other (this phenomenon can be observed clearly in the right panels of Figure 7). One way to mitigate the effect of this miss-assignment issue in the analysis of the results is to visualize the simulated and inferred scenarios using what we call a connectivity graph. This connectivity graph represents the times at which migration changes against the values of the migration rates. Such connectivity graphs are featured in the next section.

### 3.2. Validation using T-sim IICRs

The connectivity graphs and IICR plots obtained from simulated *T*_2_ values show that again the scenarios are generally very well reconstructed (Figure 7 and Figures S15 to S29).

In Figure 7 the connectivity graphs obtained for all the scenarios simulated with three and four components show that the inferred times at which migration rates changed (green vertical lines) and the inferred migration rates (green horizontal lines) are generally overlapping close to the simulated values (dotted vertical and horizontal gray lines). In the right panels of this figure, we show a subset of the inferred migration histogram. Namely, we show the distributions of the migration values that were inferred for components with a simulated migration value of *M_i_* = 10 for panel (a) and *M_i_* = 1 for panel (b). This allows us to better appreciate the variance of the inferred migration values in the context of the simulated ones, as well as the component miss-assignment effect mentioned earlier. Indeed, we note here that the incorrectly inferred migration values are clustered around other simulated values, indicating a mismatch in a particular component assignment which does not affect the rest of the inferred demographic history.

For example, consider that in the right sub-panel of (a), most repetitions correctly inferred a value close to *M* =10 for the components with that simulated migration rate. However, there were cases where a given component *i* was simulated with a migration rate of *M_i_* = 10, but it was missed entirely (maybe because it did not generate a very different IICR or because it had a short duration), and thus the inferred migration value for component *i* ultimately reflected either *M*_*i*−1_ or *M*_*i*+1_. In panel (b) we can observe the same effect with higher intensity because with more components it is more likely for them to be miss-aligned or miss-identified during inference.

These connectivity graphs (and the one obtained for five components shown in Figure S27) also show that there are regions of the parameter space where the green lines are more widely distributed. For instance, in the recent past of Figure 7b (*t_i_* < 10^-3^ generations) when the simulated *M_i_* value was 0.1 or 0.2 the inferred values seem to vary between 0.05 and 0.3, suggesting that the method identifies periods with low migration rates but that the exact value is difficult to estimate properly, at least in the recent times. These graphs however summarize extremely different scenarios, including scenarios in which consecutive *M_i_* values may be similar. It is thus important to stress that the quality of the inference is likely dependent on the timing of the changes in migration rates and on the size of the change in *M_i_* values.

Figure 8 shows the results for four different scenarios. In each of the four panels, we represented the inferred and target IICR plots, connectivity graphs, *N* (the size of each the islands) and *n* (the number of islands) for the corresponding model. Panels (a) and (b) correspond to three- and four-component scenarios, whereas panels (c) and (d) show the results for two five-component scenarios, one for which we obtained very good estimates and one for which the estimates are not very good. In panels (b) and (c) the inferred and simulated *M_i_* and *t_i_* values are on top of each other as can be seen in the connectivity graphs. Similarly, *N* and *n* are also well estimated. Here, the IICR plots also overlap, although this does not always guarantee perfect parameter estimation, as is the case in panels (a) and (d). Interestingly, in panel (a) the first change in migration rate (at *t*_1_ = 200 generations) is estimated at around 900 generations due to the stochasticity of the IICR plot. This appears to generate some variance in the estimates of *N* and *n* but the connectivity graph shows the same trend (increasing connectivity) as in the simulations. In the case of panel (d) we can see that the method had some difficulty in estimating several of the changes in *M_i_* values. This is not surprising as some of the randomly simulated changes do not seem to lead to major changes in the IICR curves. This generates again some variance in the *N* and *n* estimates. We also observe a significant variance in the connectivity graph even if several runs overlap nearly perfectly with the simulated connectivity graph.

Altogether the validation tests and figures above suggest that our framework is able to infer changes in connectivity under the *n*-island model, and that some scenarios can be extremely well inferred whereas others may be more difficult depending on their effect on the IICR plots. We also observe that for real data it may be helpful to run the analyses for a varying number of rounds, since too few rounds may negatively affect the quality of the fit. Also, once a scenario has been inferred it is crucial to test whether this scenario could indeed be properly inferred with data simulated under the newly inferred scenario. This is what we do with the real human data in the next section.

### 3.3. Application to humans

Here we show the results of using our inferential method on the human data and compare the obtained demographic scenarios with others recently published in the literature. The demographic stories for the human populations are partially found in Figures 9 and 10 (a more comprehensive set of results can be found in the supplementary materials, Figures S28 to S33).

In Figure 9, each sub-panel of panel (a) shows the PSMC curve of a human individual from a specific “population” together with the best fitting IICRs for models with *c* = 2, 3, 4 and 5 components. The different sub-panels cover the different values for the weight-shifting parameter ω (from left to right: *ω* = 1, 0.5 and 0.2); and the three humans we selected (from top to bottom: French, Karitiana and Yoruba) as explained above. The most striking feature of these plots is the sensitivity of the fit to the value of the weight-shifting parameter *ω*. Smaller values of this parameter allow the optimizer to distribute more of the demographic events towards the ancient past and thus allows this region to be better fitted by the inferred IICR. This functionality (together with being able to ignore certain parts of the plots for the computation of the distance function) can be used to make explicit the knowledge (or beliefs) of the researcher regarding the accuracy of the PSMC curve. We notice that the Yoruba population cannot be well fitted in the recent past for any value of *ω*, even outside of the designated period of recent population expansion. Panel (b) shows the connectivity graphs of the same inferred demographic scenarios. Again, the effects of varying *ω* are readily noticeable. It is also of interest that the pattern of a period of relatively high connectivity between roughly 500 kya and 2 Mya agrees with previous results published in Rodríguez et al. (2018).

Figure 10 shows in panel (a) the number of demes *n* and in panel (b) the reference size *N* that was inferred from each of the five fitted human PSMCs. Of note here is that all individuals except the Yoruba show a consistent value for these inferred parameters across both number of components and value of *ω*. The larger variance of the estimated values for the Yoruba individual suggests that a symmetrical island model may not be enough to explain the patterns of diversity in all five sampled human IICRs.

Figure 11 shows the results of the validations using seq-sim IICRs. For the three chosen representative human populations (French, Karitiana and Yoruba), we selected the SNIF inferences obtained with the parameter values *c* = 5 components and *ω* = 0.2. The selection of these values as the preferred ones was supported by the fact that the *d*_visual_ distance (see Figure S36) was best for them. For each of these inferred models we generated two independent genomic sequences of length 3 × 10^9^ base pairs and applied the PSMC method to obtain two independently estimated target IICRs. These seq-sim IICRs are shown in blue on the left panels of Figure 11. The connectivity graphs associated with these inferred models are represented in the middle panels by the red curves. The corresponding inferred values of *n* and *N* are marked as the black reference circles in the right panels of the figure.

After obtaining the seq-sim IICRs, we applied again our inference method. The goal is to validate whether it would be able to consistently infer the same parameter values of the demographic history regardless of the origin of the source IICR curve. To this end we performed 10 independent inferences from each of the two target IICRs. The inferred IICRs are superimposed on the left panels of Figure 11 (transparent red curves, 20 per population). The inferred connectivity graphs are shown in the middle panels (transparent green curves) and the inferred values of *n* and *N* are presented in the right panels. We observe a general agreement with the previously inferred histories, which suggests that *if* the real history of human evolution were close to piecewise *n*-island models like those used in this work, our method would be able to infer the parameters properly, and they would be similar to those shown in Figures 9 and 10.

Figures 12 and 13 show the results of applying our method to the IICR curves associated with the demographic model for human expansion published by Noskova et al. (2019), which we will refer to as the Classical 3-Populations model—C3PO for short. The C3PO model is a tree-like model with three modern populations that exchange gene flow asymmetrically. It is based on the model of Gutenkunst et al. (2009) and has the same structure but with a higher likelihood and thus can be seen as an improved model with a better fit to the data. The model stipulates the existence of an ancestral population that experienced an increase in size around 275 thousand years ago (Kya), and the a splitting event at about 150 Kya. This split resulted in two populations that exchanged gene flow asymmetrically: a large one that eventually became the modern African population, and a smaller one ancestral to the modern Eurasian population. This ancestral lineage split about 22 Kya into the precursors of the European and Asian populations, which at this point began an exponential increase in size that continued to present day. During this period, all three lineages continued to exchange gene flow asymmetrically. The times for these resize and splitting events are represented as dotted vertical gray lines in Figure 12.

It is clear that the nature of this model does not lend itself to be exactly modelled by a symmetrical *n*-island model, but the piecewise-stationarity of our family of models should still be able to pick up the main demographic events. For example, from an *n*-island perspective, a merger or joining of two populations (going backwards in time) may be represented by an increase in gene flow, although this effect may be confounded by the actual changes in both the sizes of the populations and migration rates taking place during these events. Also of note is the fact that the first merger event is not visible to our method because it marks the start of the recent population expansion and is thus excluded from the distance computation.

As can be seen in panel (a) of Figure 12, these IICRs do not exhibit any major features past the 300 Kya mark, so they don’t agree with the human PSMCs of Figure 5 (of which the representative ones are again shown in Figure 12 in dashed trace for reference), and they also don’t generate significant events in the inferred demographic histories. Particularly, varying the value of the weight-shifting parameter *ω* did not make a great effect in this set of inferences (which is in contrast with the results shown in Figure 9). This inferred demographic history can be roughly summarized from panel (b) as having a period of relative high gene flow followed by a sharp decrease near the 300 Kya mark, which can be very clearly attributed to the size decrease of the ancestral population in the C3PO model.

The inferred number of demes and their relatives sizes for each population can be observed in Figure 13. The numbers for the African population is in sharp contrast with the other two populations. We can also observe that for the three populations there is more variance (compared to the results from Figure 10) in the inferred values of *n* and *N* across the different values of *c* and *ω*. This may indicate a weaker link to an underlying *n*-island model.

In general, there is little agreement between the demographic histories inferred by our method from the PSMC data and the simulated IICRs from the C3PO model. This is expected because of how the two models have fundamentally incompatible structures, not only regarding the island versus tree aspect, but also due to the size changes in the C3PO model that affect the IICR potentially as much as gene flow does. However, we do identify the approximate timings of the two visible demographic events when using *c* = 5 components and the more recent-weighted value of *ω* = 1. These results also serve as additional validation that our method will not return the same parameter values regardless of the source of the data. They also suggest that the C3PO model is unlikely to be a good model to understand questions about ancient human structure and evolution.

## 4. Discussion

Our validations show that the inference framework presented here is able to accurately infer structure parameters (number of islands and their sizes) within a symmetrical island model given an IICR estimate like the PSMC. It is also able to date up to five events of changes in migration rate (i.e., six components) with good precision and consistency, as long as the underlying model is compatible with a symmetrical island model. The migration rates themselves are inferred as well.

### Human evolution

An application of our method to five publicly available human PSMCs suggests that the backwards long term history of the sampled individuals, when accounting for possible recent expansions and the noise introduced by the PSMC method, can be accurately modelled in the framework of a symmetrical island model of approximately 10 to 12 demes with varying levels of connectivity through time. Only one of the five samples (Yoruba) displayed less consistent evidence of this finding, which may indicate that more complex models (possibly including asymmetric gene flow, spatial modelling of the environment, or changes in deme sizes) could be needed to explain the full complexity of the data.

These findings are in agreement with the results of Rodríguez et al. (2018), in which a hand-fitting approach of the IICRs was used to arrive at an estimate of 10 islands with a similar value of *N* and a comparable period featuring a significant increased of gene flow between 600 Kya and 2 Mya.

We also compared our results with the tree model for human evolution published by Noskova et al. (2019) (the C3PO model), which is a revision of the model from Gutenkunst et al. (2009) and represents a widespread view of human evolution (Jouganous et al., 2017; Kamm et al., 2019). The C3PO model proposes an ancestral human population that experiences two splits: an old one that resulted in the current African “population” and another more recent one that resulted in the current European and Asian “populations”. The parameters of this model include the times of these events, the population size history of these populations and their ancestral branches, and the migration rates between them. The summary statistic targeted by these methods is the Allele Frequency Spectrum (AFS), and we see that a fitting AFS does not guarantee a fitting IICR and vice versa (Chikhi et al., 2018). Indeed, the IICRs of the populations from the C3PO model do not resemble those of the real humans. Likewise, the inferred demographic history from the IICRs curves generated from the C3PO model are less consistent across the various runs of our method and they do not resemble those obtained from the human IICRs.

These findings suggest that tree models fitted with the AFS like those considered here do not offer a definitive answer to the question of human evolution and other families of models should be explored (Goldstein and Chikhi, 2002; Scerri et al., 2018, 2019). It remains to be seen however how well do the models inferred with our method fit the real AFS of their respective human populations, but this would be beyond the work presented here.

### Future work

One novel aspect of our approach is that the number of demes gets inferred as one of the model parameters, and it is in fact the best estimated parameter, in the sense that the robustness of the estimated value scales well with increasing model complexity. A similar consistency can be observed with the deme size parameter *N*. We give up some flexibility in the model by keeping the number of demes constant throughout the history of the population, so the timed demographic events cannot represent splits or joining of lineages. Additionally, in the *n*-island model we do not account for possible asymmetrical gene flow or different deme sizes, even when the theoretical framework does allow for such representations. However, it is a more challenging problem to validate due to the fact that during any given component, changing both the migration rate and the deme size have confounding effects on the IICR curve which can be hard to separate. This requires a dedicated study with a different methodology which we will explore in a future work.

Another potential direction is to use multiple IICR curves simultaneously during the inference process. These multiple IICRs may come in the form of more than one IICR sampled from an asymmetrical demographic model (for which the initial sampling deme *does* result in different curves (Chikhi et al., 2018), as opposed to the *n*-island model where demes are by definition indistinguishable). They may also be in the form of multiple IICR_*k*_ curves where *k* is the number of sampled haploid genomes. Indeed, the IICR of Mazet et al. (2016) was defined for *k* = 2, and this is the IICR that we have been studying in our previous works. However, the concept can be extended to more haploid genomes in the same way that the MSMC method (Schiffels and Durbin, 2013) is an extension of the PSMC to multiple genomes, which takes into consideration the distribution of the coalescent time *T_k_*. The precise concept of the IICR_*k*_ is currently being developed in a separate paper (Rodríguez et al., 2020). These approaches may prove beneficial in choosing between structured and non-structured models. Indeed, Grusea et al. (2018) shows that using more than one IICR curve can help discriminate between structured and non structured scenarios in the *n*-island model. Finally, the incorporation of larger samples not only enables exploring more complex scenarios, but it also allows for using other summary statistics to complement the IICR, most notably among them the Allele Frequency Spectrum (AFS), which is widely used for the purposes of demographic inference.

### Conclusion

In summary, we have presented here an inference method for automatically estimating demographic parameters under a piecewise stationary symmetrical island model that uses the IICR as its summary statistic. The underlying methodology consists in quantifying the discrepancy between a target IICR and many simulated IICR curves for a large number of candidate scenarios, and using this metric to drive a global optimization process. With a large number of validations we have shown that the method works accurately and consistently for a diverse range of parameter values, and we additionally showed an application to human data that agrees with and improves upon previously published results using similar approaches.

We believe that despite it’s current scope, our method can be of great value during the initial exploration of the parameter space for simple models, and thus can also provide a starting point for manually fitting the IICR with models that could express spatial structure and varying *N* (Rodríguez et al., 2018).

## 5. Acknowledgements

We would like to thank Simona Grusea, Josué Corujo, Pierre Lacoste and Rémi Tournebize, for their input and suggestions over many lengthy and productive discussions. Armando Arredondo was funded by the Université Fédérale Toulouse Midi Pyrénées (UFTMiP) and the Région Occitanie (formerly Midi-Pyrénées) with PhD grant No. 31I2017M248. Lounès Chikhi was funded by Fundação para a Ciência e Tecnologia (ref. PTDC-BIA-EVL/30815/2017). Olivier Mazet and Lounès Chikhi were funded by the 2015–2016 BiodivERsA COFUND call for research proposals, with the national funders ANR (ANR-16-EBI3-0014) and the Fundação para a Ciência e Tecnologia ref. Biodiversa/0003/2015 and PT-DLR (01LC1617A). This work was also supported by the LABEX entitled TULIP (ANR-10-LABX-41 and ANR-11-IDEX-0002-02) as well as the LIA BEEG-B (Laboratoire International Associé-Bioinformatics, Ecology, Evolution, Genomics and Behaviour). We acknowledge an Investissement d’Avenir grant of the Agence Nationale de la Recherche (CEBA: ANR-10-LABX-25-01). We are grateful to the GenOuest bioinformatics platform for providing computing and storage resources.

## 7. Data Archiving

All software documentation and data required to reproduce our results will be made available upon acceptance of the manuscript.

## References

Beaumont, M. P. (2004). Recent developments in genetic data analysis: what can they tell us about human demographic history? Heredity, 92(5):365–379.

Beaumont, M. A. and Nichols, R. A. (1996). Evaluating loci for use in the genetic analysis of population structure. Proceedings of the Royal Society of London. Series B: Biological Sciences, 263(1377):1619–1626.

Boitard, S., Rodríguez, W., Jay, F., Mona, S., and Austerlitz, F. (2016). Inferring population size history from large samples of genome-wide molecular data-an approximate bayesian computation approach. PLoS Genetics, 12(3):e1005877.

Cavalli-Sforza, L. L. (1966). Population structure and human evolution. Proceedings of the Royal Society of London. Series B. Biological Sciences, 164(995):362–379.

Chikhi, L., Rodríguez, W., Grusea, S., Santos, P., Boitard, S., and Mazet, O. (2018). The IICR (inverse instantaneous coalescence rate) as a summary of genomic diversity: insights into demographic inference and model choice. Heredity, 120:13–24.

Chikhi, L., Sousa, V. C., Luisi, P., Goossens, B., and Beaumont, M. A. (2010). The confounding effects of population structure, genetic diversity and the sampling scheme on the detection and quantification of population size changes. Genetics, 186(3):983–995.

Excoffier, L., Dupanloup, I., Huerta-Sánchez, E., Sousa, V. C., and Foll, M. (2013). Robust demographic inference from genomic and snp data. PLoS Genetics, 9(10):e1003905.

Fenderson, L. E., Kovach, A. I., and Llamas, B. (2020). Spatiotemporal landscape genetics: Investigating ecology and evolution through space and time. Molecular Ecology, 29(2):218–246.

Goldstein, D. B. and Chikhi, L. (2002). Human migrations and population structure: what we know and why it matters. Annual Review of Genomics and Human Genetics, 3(1):129–152.

Goossens, B., Chikhi, L., Ancrenaz, M., Lackman-Ancrenaz, I., Andau, P., Bruford, M. W., et al. (2006). Genetic signature of anthropogenic population collapse in orang-utans. PLoS Biology, 4(2):285.

Grusea, S., Rodríguez, W., Pinchon, D., Chikhi, L., Boitard, S., and Mazet, O. (2018). Coalescence times for three genes provide sufficient information to distinguish population structure from population size changes. Journal of Mathematical Biology, 78(1-2):189–224.

Gutenkunst, R. N., Hernandez, R. D., Williamson, S. H., and Bustamante, C. D. (2009). Inferring the joint demographic history of multiple populations from multidimensional SNP frequency data. PLoS Genetics, 5(10):e1000695.

Hecht, L. B., Thompson, P. C., and Rosenthal, B. M. (2018). Comparative demography elucidates the longevity of parasitic and symbiotic relationships. Proceedings of the Royal Society B, 285(1888):20181032.

Hecht, L. B., Thompson, P. C., and Rosenthal, B. M. (2020). Assessing the evolutionary persistence of ecological relationships: a review and preview. Infection, Genetics and Evolution, page 104441.

Herbots, H. M. J. D. (1994). Stochastic models in population genetics: genealogy and genetic differentiation in structured populations. PhD thesis.

Hey, J. and Machado, C. A. (2003). The study of structured populations–new hope for a difficult and divided science. Nature Reviews. Genetics, 4(7):535.

Hobolth, A., Siri-Jegousse, A., and Bladt, M. (2019). Phase-type distributions in population genetics. Theoretical Population Biology, 127:16–32.

Jouganous, J., Long, W., Ragsdale, A. P., and Gravel, S. (2017). Inferring the joint demographic history of multiple populations: beyond the diffusion approximation. Genetics, 206(3):1549–1567.

Kamm, J., Terhorst, J., Durbin, R., and Song, Y. S. (2019). Efficiently inferring the demographic history of many populations with allele count data. Journal of the American Statistical Association, pages 1–16.

Kaplan, E. L. and Meier, P. (1958). Nonparametric estimation from incomplete observations. Journal of the American Statistical Association, 53(282):457–481.

Kuhlwilm, M., Gronau, I., Hubisz, M. J., de Filippo, C., Prado-Martinez, J., Kircher, M., Fu, Q., Burbano, H. A., Lalueza-Fox, C., de La Rasilla, M., et al. (2016). Ancient gene flow from early modern humans into eastern neanderthals. Nature, 530(7591):429–433.

Li, H. and Durbin, R. (2011). Inference of human population history from individual whole-genome sequences. Nature, 475(7357):493–496.

Liu, X. and Fu, Y.-X. (2015). Exploring population size changes using SNP frequency spectra. Nature Genetics.

Mazet, O., Rodríguez, W., Grusea, S., Boitard, S., and Chikhi, L. (2016). On the importance of being structured: instantaneous coalescence rates and human evolution—lessons for ancestral population size inference? Heredity, 116(4):362.

Noskova, E., Ulyantsev, V., Koepfli, K.-P., O’Brien, S. J., and Dobrynin, P. (2019). Gadma: Genetic algorithm for inferring demographic history of multiple populations from allele frequency spectrum data. bioRxiv.

Poelstra, J., Salmona, J., Tiley, G. P., Schußler, D., Blanco, M. B., Andriambeloson, J. B., Manzi, S., Campbell, C. R., Bouchez, O., Etter, P. D., et al. (2020). Cryptic patterns of speciation in cryptic primates: microendemic mouse lemurs and the multispecies coalescent. Systematic Biology, page 742361.

Prado-Martinez, J., Sudmant, P. H., Kidd, J. M., Li, H., Kelley, J. L., Lorente-Galdos, B., Veeramah, K. R., Woerner, A. E., O’Connor, T. D., Santpere, G., et al. (2013). Great ape genetic diversity and population history. Nature, 499(7459):471–475.

Quéméré, E., Amelot, X., Pierson, J., Crouau-Roy, B., and Chikhi, L. (2012). Genetic data suggest a natural prehuman origin of open habitats in northern Madagascar and question the deforestation narrative in this region. Proceedings of the National Academy of Sciences, 109(32):13028–13033.

Rodríguez, W., Boitard, S., Grusea, S., Arredondo, A., Corujo, J., Mazet, O., and Chikhi, L. (2020). Extending the IICR to multiple genomes to get insights into demographic history of species. Manuscript in preparation.

Rodríguez, W., Mazet, O., Grusea, S., Arredondo, A., Corujo, J. M., Boitard, S., and Chikhi, L. (2018). The IICR and the non-stationary structured coalescent: towards demographic inference with arbitrary changes in population structure. Heredity, 121(6):663.

Rogers, A. R. and Harpending, H. (1992). Population growth makes waves in the distribution of pairwise genetic differences. Molecular Biology and Evolution, 9(3):552–569.

Salmona, J., Heller, R., Quéméré, E., and Chikhi, L. (2017). Climate change and human colonization triggered habitat loss and fragmentation in madagascar. Molecular Ecology, 26(19):5203–5222.

Scerri, E. M., Chikhi, L., and Thomas, M. G. (2019). Beyond multiregional and simple out-of-Africa models of human evolution. Nature Ecology & Evolution, 3(10):1370–1372.

Scerri, E. M. L., Thomas, M. G., Manica, A., Gunz, P., Stock, J. T., Stringer, C., Grove, M., Groucutt, H. S., Timmermann, A., Rightmire, G. P., d’Errico, F., Tryon, C. A., Drake, N. A., Brooks, A. S., Dennell, R. W., Durbin, R., Henn, B. M., Lee-Thorp, J., deMenocal, P., Petraglia, M. D., Thompson, J. C., Scally, A., and Chikhi, L. (2018). Did our species evolve in subdivided populations across Africa, and why does it matter? Trends in Ecology & Evolution.

Schiffels, S. and Durbin, R. (2013). Inferring human population size and separation history from multiple genome sequences. Nature Genetics, 8(46):919–925.

Steinrücken, M., Kamm, J., Spence, J. P., and Song, Y. S. (2019). Inference of complex population histories using whole-genome sequences from multiple populations. Proceedings of the National Academy of Sciences, 116(14):1–6.

Storn, R. and Price, K. (1997). Differential evolution–a simple and efficient heuristic for global optimization over continuous spaces. Journal of Global Optimization, 11(4):341–359.

Tavaré, S. (2004). Part I: Ancestral inference in population genetics. In Lectures on probability theory and statistics, pages 1–188. Springer.

Wakeley, J. (1999). Nonequilibrium migration in human history. Genetics, 153(4):1863–1871.

Wang, K., Mathieson, I., O’Connell, J., and Schiffels, S. (2020). Tracking human population structure through time from whole genome sequences. PLoS Genetics, 16(3):e1008552.

Wright, S. (1931). Evolution in Mendelian populations. Genetics, 16(2):97.

