## Supplementary Materials for "Inferring number of populations and changes in connectivity under the n-island model"

**5 Results of application to human data** **32**

**6 A note on implementation and use cases** **39**

### 1. Proof of Lemma 1

**Lemma 1.** *Given any* $\hat{\varphi} = (N, n, t_1 \dots t_\gamma, M_0 \dots M_\gamma, s_0 \dots s_\gamma) \in \hat{\Phi}_{\gamma,B}$*, then the parameter tuple:*

$$\hat{\varphi}_0 = \Big(\frac{N}{C}, n, Ct_1 \dots Ct_\gamma, \frac{M_0}{C} \cdots \frac{M_\gamma}{C}, Cs_0 \dots Cs_\gamma\Big)$$

*is such that:*

$$\mathrm{sIICR}_{\hat{\varphi}}(g) = \mathrm{sIICR}_{\hat{\varphi}_0}(g) \qquad \forall g > 0,$$

*where* $C$ *is any rescaling factor for which the coordinates of* $\hat{\varphi}_0$ *are within the bounds* $B$*.*

*Proof.* Let us denote by $\pi_{n,M,s}(t)$ the factors $e^{tQ}$ that appear in the definition of $P(t)$ (equations 2.4 and 2.5). It is easy to verify that for any $C > 0$, we have:

$$\pi_{n,M,s}(t) = \pi_{n,\frac{M}{C},Cs}(Ct). \tag{1}$$

Indeed, from the expressions in (2.6) we can see that the parameter $M$ always appears in the factor $Ms$, and the parameter $t$ always appears in the factor $t/s$, which are invariant under the transformation $(M, s, t) \mapsto (\frac{M}{C}, Cs, Ct)$.

Next, given any:

$$\varphi = (n, t_1 \dots t_\gamma, M_0 \dots M_\gamma, s_0 \dots s_\gamma),$$
$$\varphi_0 = (n, Ct_1 \dots Ct_\gamma, \frac{M_0}{C} \cdots \frac{M_\gamma}{C}, Cs_0 \dots Cs_\gamma),$$

and any $t > 0$, we can write $P(t)$ from (2.4) as:

$$\begin{aligned}
P_\varphi(t) &= \Big(\prod_{k=1}^{i} \pi_{n,M_{k-1},s_{k-1}}(t_k - t_{k-1})\Big)\pi_{n,M_i,s_i}(t - t_i) \\
&= \Big(\prod_{k=1}^{i} \pi_{n,\frac{M_{k-1}}{C},Cs_{k-1}}(Ct_k - Ct_{k-1})\Big)\pi_{n,\frac{M_i}{C},Cs_i}(Ct - Ct_i) \\
&= P_{\varphi_0}(Ct).
\end{aligned} \tag{2}$$

where $i$ is the largest index such that $t_i \leqslant t$ and subsequently $Ct_i \leqslant Ct$. We denote now by $F_\varphi(t)$ any of $F_{\mathrm{same}}(t) = P_\varphi(t)_{(1,3)}$ or $F_{\mathrm{diff.}}(t) = P_\varphi(t)_{(1,2)}$. From (2) we have $F_\varphi(t) = F_{\varphi_0}(Ct)$. In order to introduce scaling, we consider an arbitrary effective size $N$ and perform the corresponding change of variable $t = g/2N$. Thus, by having $\hat{\varphi} = (N, \varphi)$ and $\hat{\varphi}_0 = (N/C, \varphi_0)$, we get:

$$\begin{aligned}
& F_\varphi\Big(\frac{g}{2N}\Big) = F_{\varphi_0}\Big(\frac{Cg}{2N}\Big) \\
\Rightarrow \quad & \frac{1}{2N} f_\varphi\Big(\frac{g}{2N}\Big) = \frac{C}{2N} f_{\varphi_0}\Big(\frac{Cg}{2N}\Big) \\
\Rightarrow \quad & N\,\frac{1 - F_\varphi(g/2N)}{f_\varphi(g/2N)} = \frac{N}{C}\,\frac{1 - F_{\varphi_0}(Cg/2N)}{f_{\varphi_0}(Cg/2N)} \\
\Rightarrow \quad & \mathrm{sIICR}_{\hat{\varphi}}(g) = \mathrm{sIICR}_{\hat{\varphi}_0}(g)
\end{aligned}$$

■

### 2. Comparison of optimization parameters

Here we explore the effect of various parameters of the optimization algorithm on the speed of convergence of the inference process.

The most important parameters that affect the search process and convergence criteria are:

- `strategy`: it can be one of 12 possible values (see Table 1) and controls how each new generation of solutions are computed from the previous one. Default is '`best1bin`'.
- `maxiter`: maximum number of iterations the algorithm will perform before forcing convergence. Default is 5000.
- `popsize`: number of simultaneous solutions during any given generation. Default is 15.
- `tol`: relative tolerance for search convergence. The convergence criteria is met when the standard deviation of the solutions within a generation is smaller than `tol` times the average energy (in our case, distance) within that generation. Default is $10^{-2}$.
- `mutation`: per-generation mutation rate of the solutions. The default behaviour is to draw a random value from a uniform distribution in $[0.5, 1]$ each generation.
- `recombination`: per-generation recombination rate of the solutions. Default is $0.7$.

We selected 10 random simulated scenarios with unscaled IICR and $c = 4$ components from the set of exact IICR validations that had not converged before 500 rounds (these would correspond to scenarios off the diagonals in Figure S6). For each one of them, and for each of the 12 possible values for the `strategy` parameter, we attempted another 100 rounds. Table 1 shows the number of rounds it took for each of the 10 scenarios to converge in each case (a value of 100 means that convergence was not reached).

| Strategy | #1 | #2 | #3 | #4 | #5 | #6 | #7 | #8 | #9 | #10 | Total |
|---|---|---|---|---|---|---|---|---|---|---|---|
| best2exp | 17 | 1 | 100 | 2 | 100 | 22 | 3 | 100 | 57 | 22 | 424 |
| best2bin | 6 | 35 | 100 | 2 | 11 | 93 | 100 | 4 | 33 | 100 | 484 |
| currenttobest1exp | 2 | 80 | 100 | 5 | 100 | 69 | 12 | 100 | 8 | 26 | 502 |
| best1exp | 3 | 76 | 100 | 21 | 4 | 100 | 10 | 100 | 62 | 100 | 576 |
| rand1exp | 4 | 100 | 100 | 10 | 100 | 14 | 100 | 100 | 100 | 38 | 666 |
| randtobest1exp | 32 | 100 | 100 | 9 | 100 | 78 | 100 | 30 | 100 | 100 | 749 |
| currenttobest1bin | 21 | 100 | 100 | 27 | 100 | 100 | 100 | 94 | 63 | 88 | 793 |
| best1bin | 27 | 100 | 100 | 28 | 54 | 100 | 100 | 100 | 100 | 100 | 809 |
| rand2exp | 40 | 100 | 100 | 100 | 100 | 24 | 100 | 100 | 100 | 100 | 864 |
| rand1bin | 100 | 100 | 100 | 3 | 100 | 100 | 100 | 100 | 100 | 100 | 903 |
| rand2bin | 96 | 100 | 100 | 23 | 100 | 100 | 100 | 100 | 100 | 100 | 919 |
| randtobest1bin | 92 | 100 | 100 | 100 | 100 | 100 | 100 | 100 | 100 | 100 | 992 |

**Table 1:** Results of varying the `strategy` parameter of the differential evolution algorithm on the speed of convergence of 10 difficult demographic scenarios with $c = 4$ components.

As can be seen, strategy '`best2exp`' was best with a combined number of 424 rounds for the 10 scenarios.

Next, using this optimal strategy parameter, we tried one alternative value for the rest of the optimization parameters at a time, again allowing a maximum of 100 rounds. The alternative values were as follows:

- `maxiter` was changed from 5000 to 10000.
- `popsize` was changed from 15 to 50.
- `mutation` was changed from random sample in $[0.5, 1]$ to random sample in $[0.5, 1.7]$.
- `recombination` was changed from $0.7$ to $0.9$.

The results, shown in Table 2, suggest that these parameters should be left at their default values.

| Parameter | no. 1 | no. 2 | no. 3 | no. 4 | no. 5 | no. 6 | no. 7 | no. 8 | no. 9 | no. 10 | Total |
|---|---|---|---|---|---|---|---|---|---|---|---|
| popsize | 100 | 100 | 100 | 100 | 100 | 100 | 100 | 100 | 100 | 100 | 1,000 |
| recombination | 100 | 100 | 100 | 100 | 100 | 100 | 100 | 100 | 100 | 100 | 1,000 |
| mutation | 100 | 100 | 100 | 100 | 100 | 100 | 100 | 100 | 100 | 100 | 1,000 |
| max-iter | 100 | 100 | 100 | 100 | 100 | 100 | 100 | 100 | 100 | 100 | 1,000 |

**Table 2:** Results of varying the `popsize`, `recombination`, `mutation` and `max-iter` parameters of the differential evolution algorithm on the speed of convergence of 10 difficult demographic scenarios with $c = 4$ components.

### 3. Results of validation using exact simulated IICRs

In figures S1 and S2 we explore the question of how many rounds of inference are needed in order to achieve optimal distance. These results are only of theoretical interest since in real applications, the target IICR is not known exactly (just approximated) therefore the optimal distance of 0 cannot be achieved in any meaningful way.

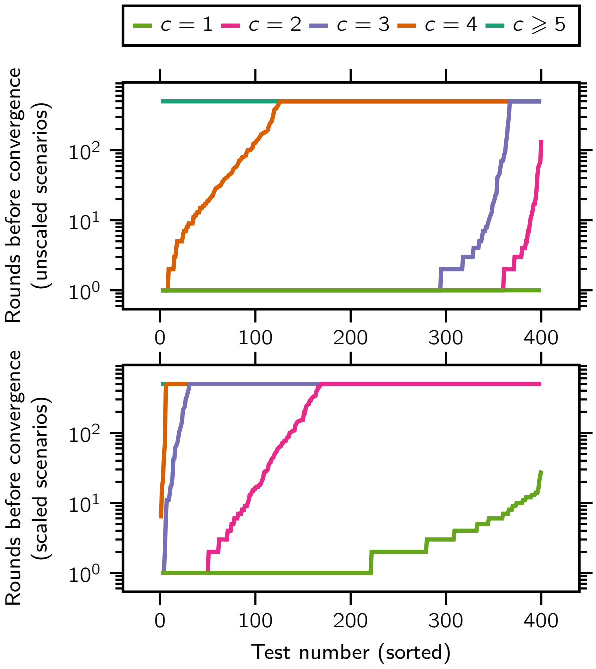

**Figure S1:** Number of performed rounds during validations with exact IICRs. The top panel shows the number of rounds (up to a maximum of 500) that the method required before converging to a scenario with a distance value smaller than the tolerance $\varepsilon = 10^{-10}$ for the unscaled case. The bottom panel shows the same information for scaled scenarios and a tolerance of $\varepsilon = 10^{-7}$. We represent in different colors the curves corresponding to scenarios with different simulated (and inferred) number of components, ranging from $c = 2$ up to $c = 6$; the same data-set represented in figures S3 to S8 (top panel) and figures S9 to S14 (bottom panel). Higher number of rounds is not correlated with worse fit when the maximum number of rounds is not reached; and when it is reached it is not necessarily an indication of bad fit either, although all instances of bad fit stem from inferences that reached the maximum allowed number of rounds.

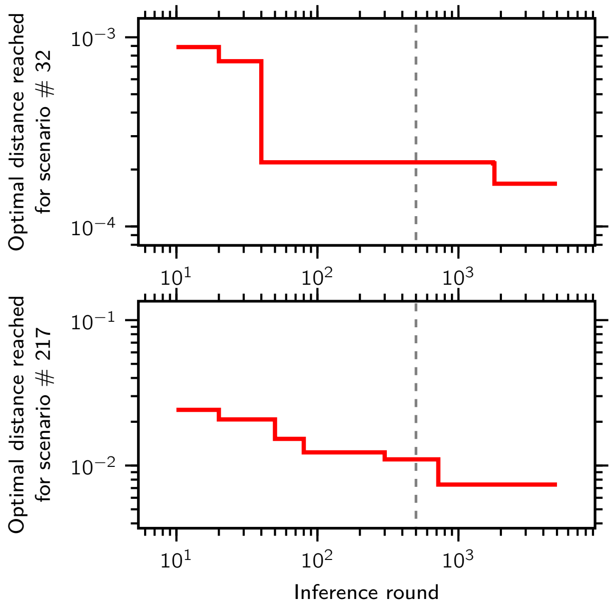

**Figure S2:** Convergence of the inferred IICR with the progression of the inference rounds. We measured the best distance achieved ($\omega = 1$) as a function of the completed rounds for up to 5000 rounds. The two curves correspond to two $c = 6$ component scenarios that didn't previously converge ($\varepsilon = 10^{-10}$) in 500 rounds (marked with a vertical dashed line). We note that there is a clear trend for the distance between the IICR curves to decrease with more inference rounds, but there is also diminishing returns to performing a very large number of rounds, since the issue of component misidentification may be an insurmountable difficulty for some scenarios (see main text for a more detailed discussion).

#### 3.1. Unscaled IICR

In this section we show the validation results of simulating and then inferring from 400 randomly generated demographic scenarios with unscaled IICRs and varying number of components $c$. Each figure consists of as many sub-panels as there are free parameters for that model, and in each one the simulated values are in the horizontal axis and the inferred ones in the vertical axis.

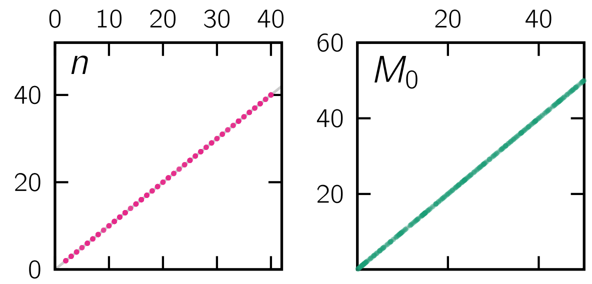

**Figure S3:** Scatter plots of all the simulated (horizontal axis) versus inferred (vertical axis) parameter values for scenarios of $c = 1$ component.

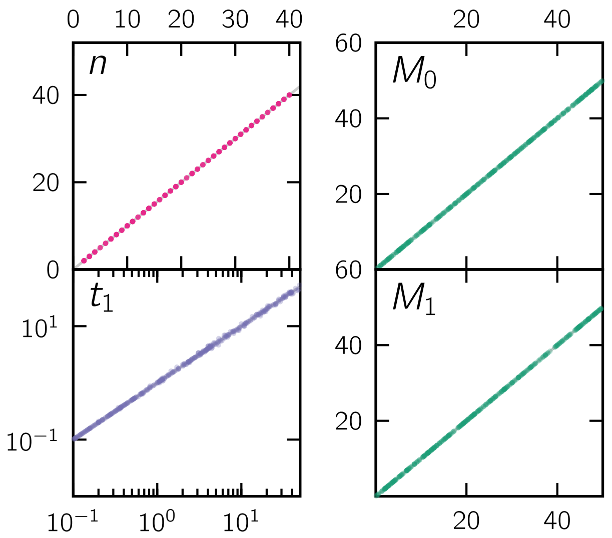

**Figure S4:** Scatter plots of all the simulated (horizontal axis) versus inferred (vertical axis) parameter values for scenarios of $c = 2$ components.

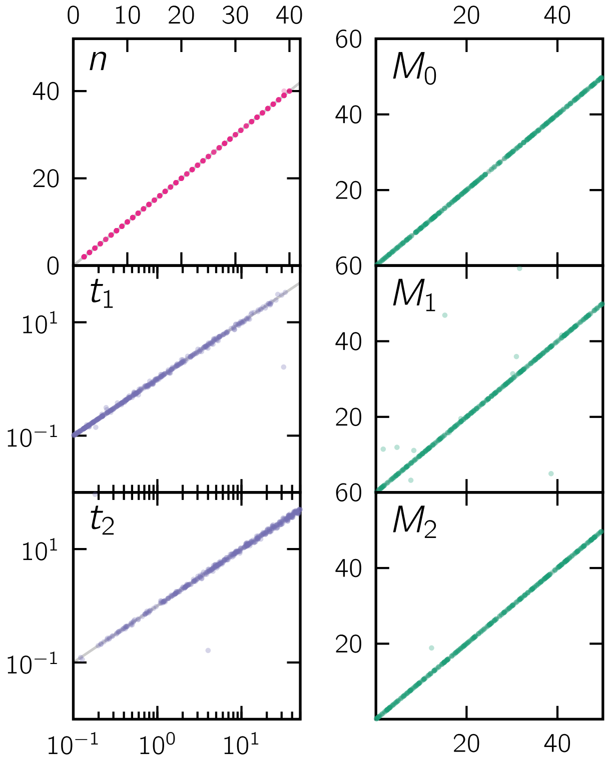

**Figure S5:** Scatter plots of all the simulated (horizontal axis) versus inferred (vertical axis) parameter values for scenarios of $c = 3$ components.

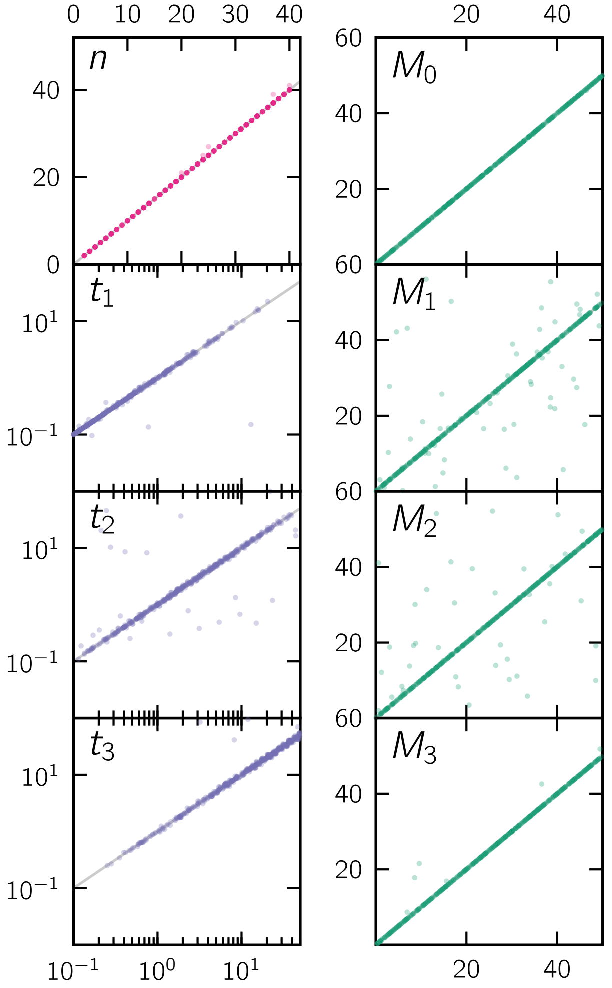

**Figure S6:** Scatter plots of all the simulated (horizontal axis) versus inferred (vertical axis) parameter values for scenarios of $c = 4$ components.

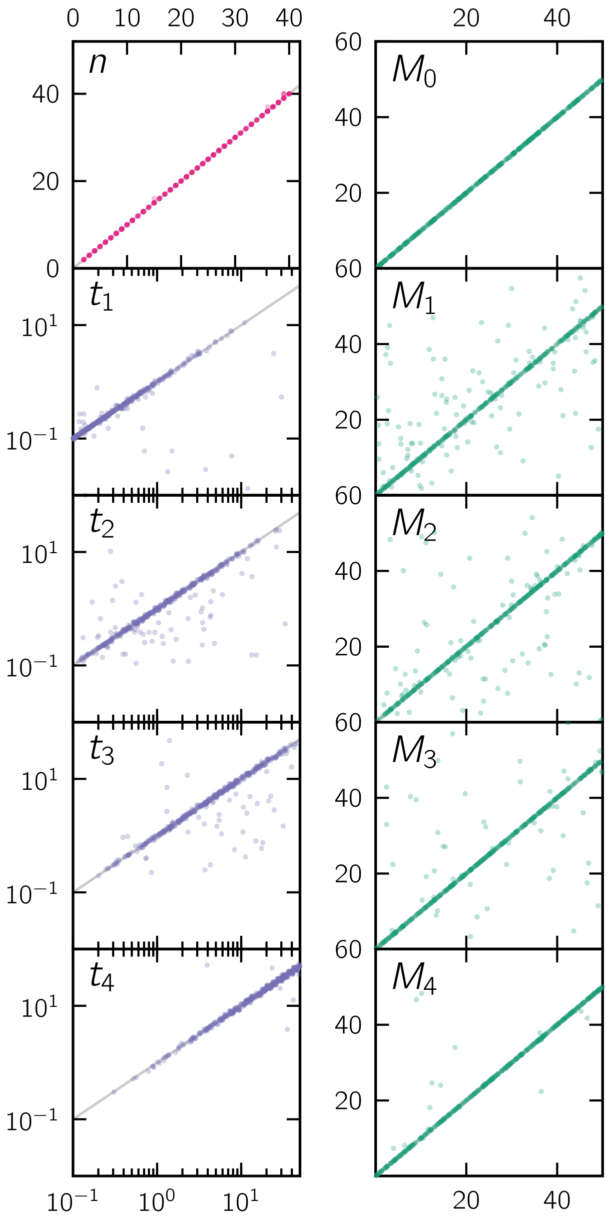

**Figure S7:** Scatter plots of all the simulated (horizontal axis) versus inferred (vertical axis) parameter values for scenarios of $c = 5$ components.

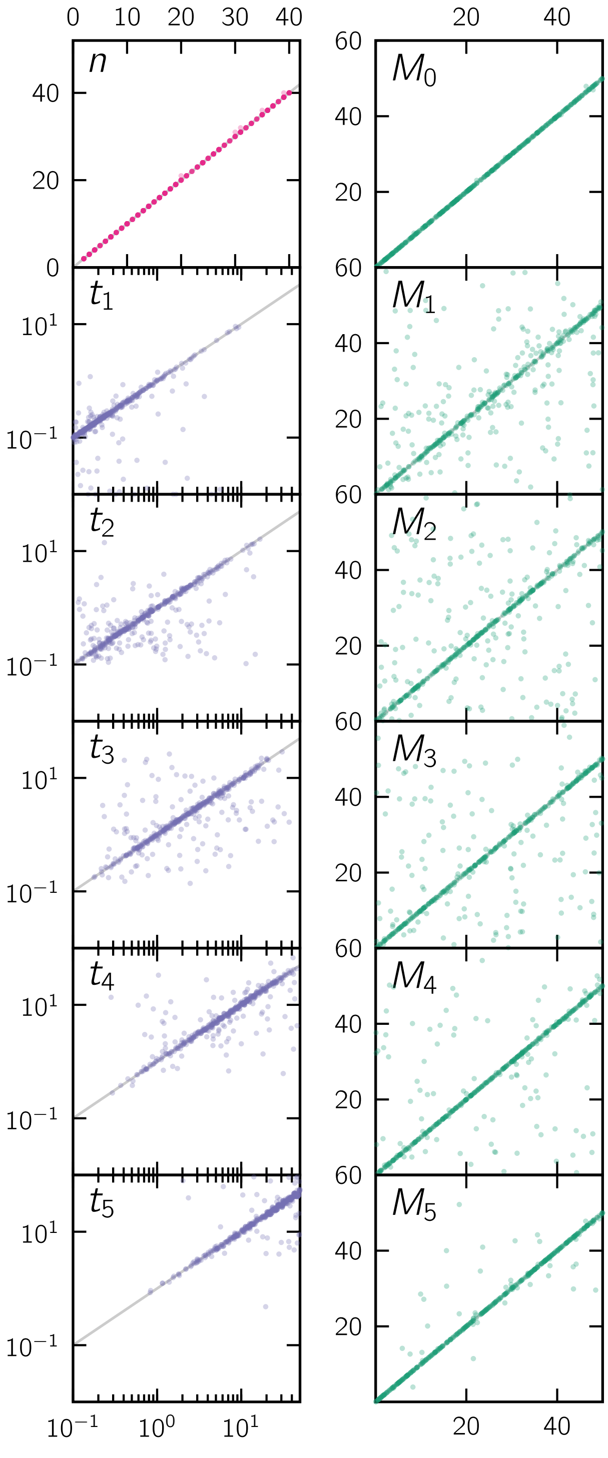

**Figure S8:** Scatter plots of all the simulated (horizontal axis) versus inferred (vertical axis) parameter values for scenarios of $c = 6$ components.

#### 3.2. Scaled IICR

In this section we show the validation results of simulating and then inferring from 400 randomly generated demographic scenarios with scaled IICRs and varying number of components $c$. Each figure consists of as many sub-panels as there are free parameters for that model, and in each one the simulated values are in the horizontal axis and the inferred ones in the vertical axis. The deme size panel ($N$) is different because the simulated values for $N$ was always $N = 1000$, therefore the horizontal axis indicates the test number (1 to 400) and the vertical axis the inferred value.

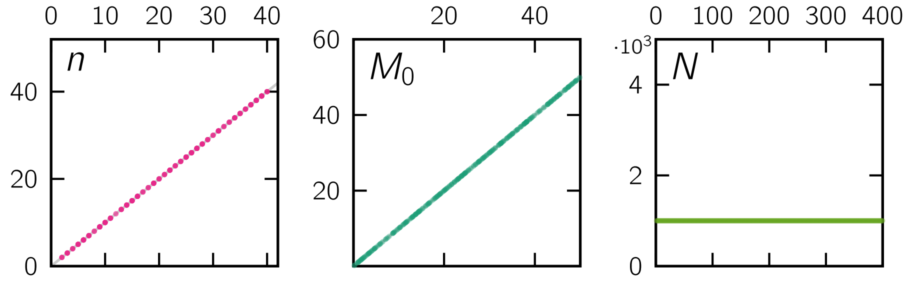

**Figure S9:** Scatter plots of all the simulated (horizontal axis) versus inferred (vertical axis) parameter values for scenarios of $c = 1$ component.

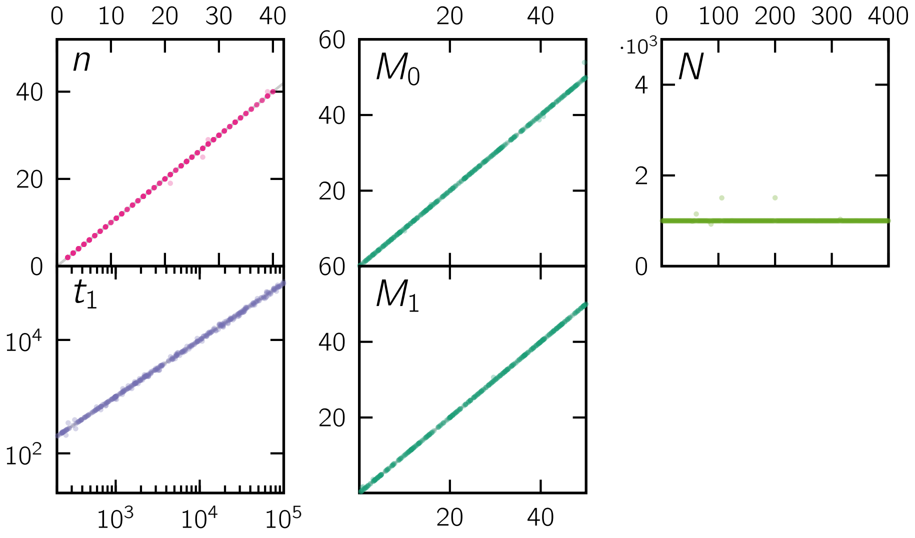

**Figure S10:** Scatter plots of all the simulated (horizontal axis) versus inferred (vertical axis) parameter values for scenarios of $c = 2$ components.

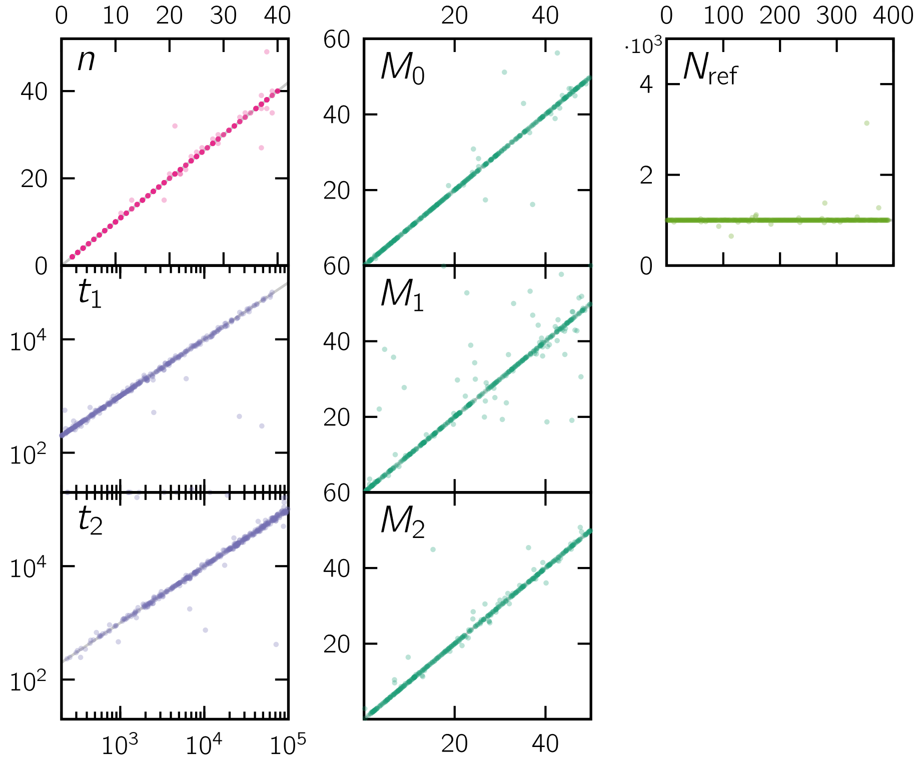

**Figure S11:** Scatter plots of all the simulated (horizontal axis) versus inferred (vertical axis) parameter values for scenarios of $c = 3$ components.

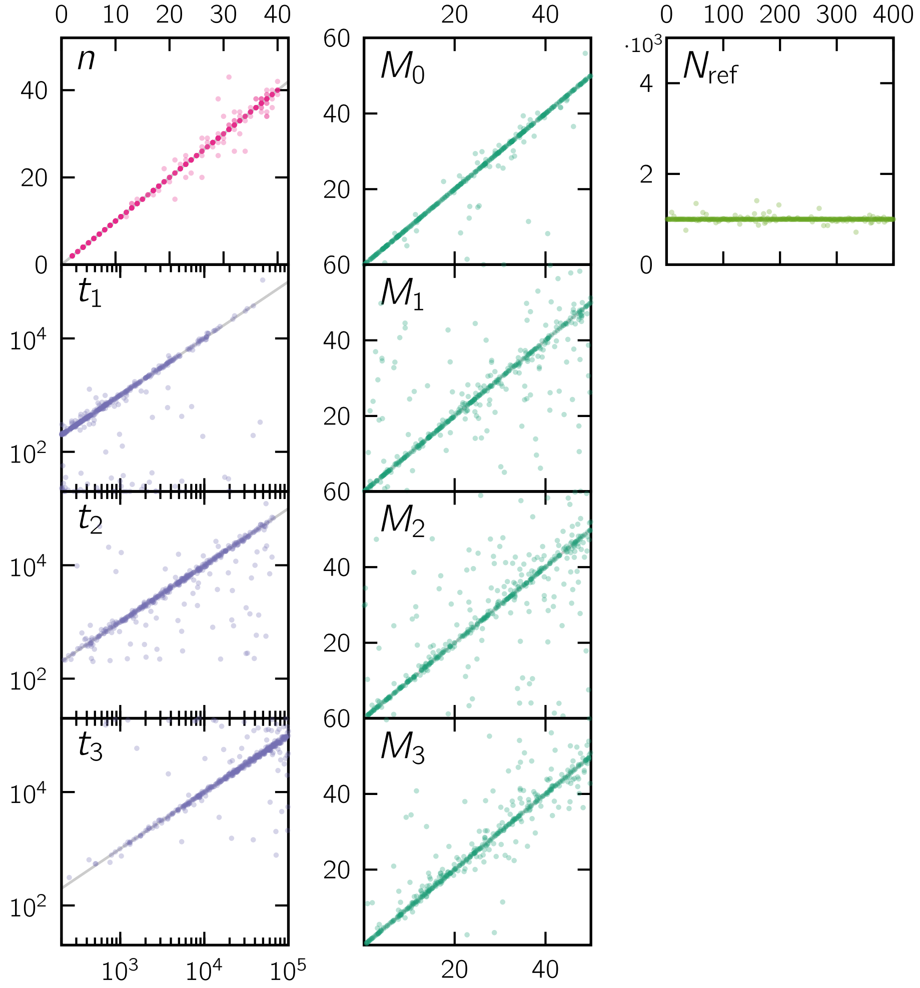

**Figure S12:** Scatter plots of all the simulated (horizontal axis) versus inferred (vertical axis) parameter values for scenarios of $c = 4$ components.

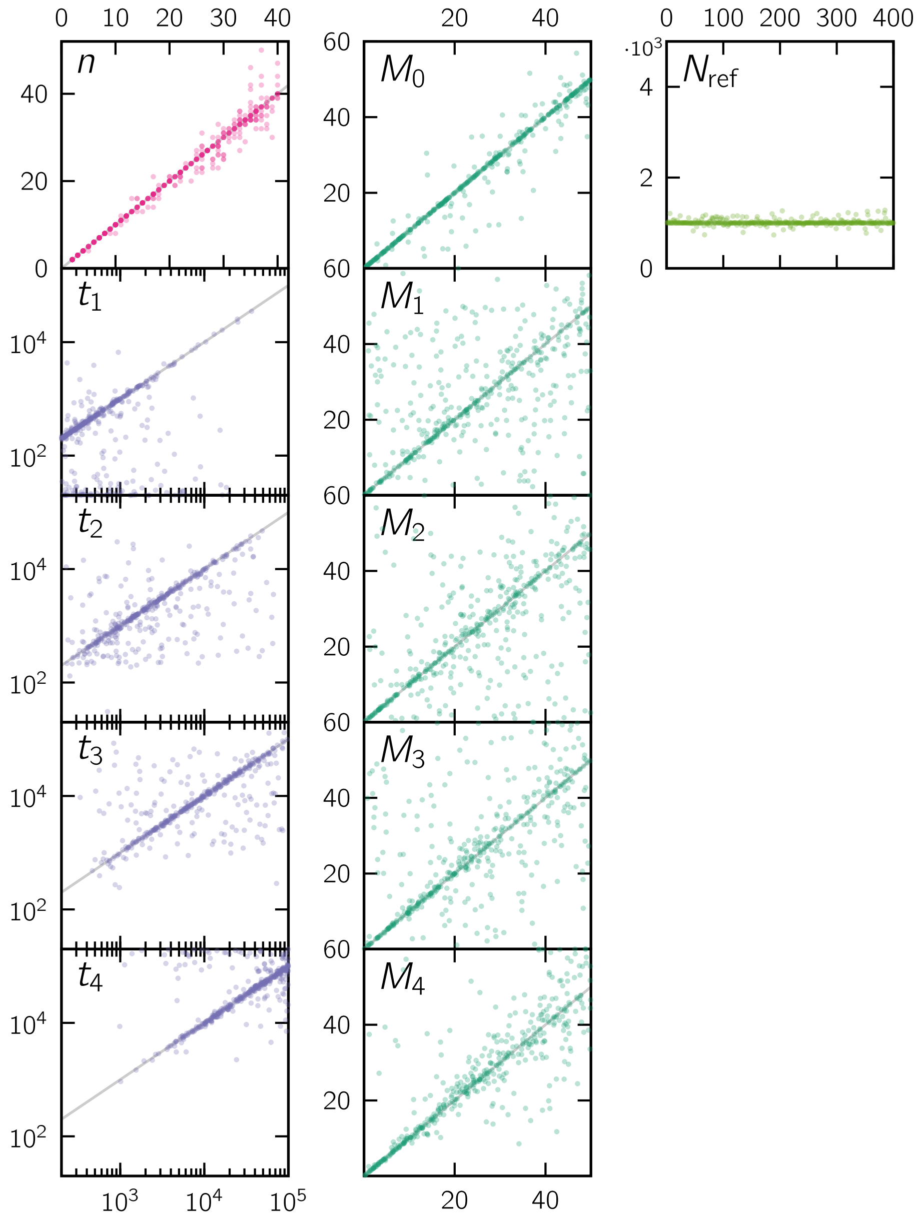

**Figure S13:** Scatter plots of all the simulated (horizontal axis) versus inferred (vertical axis) parameter values for scenarios of $c = 5$ components.

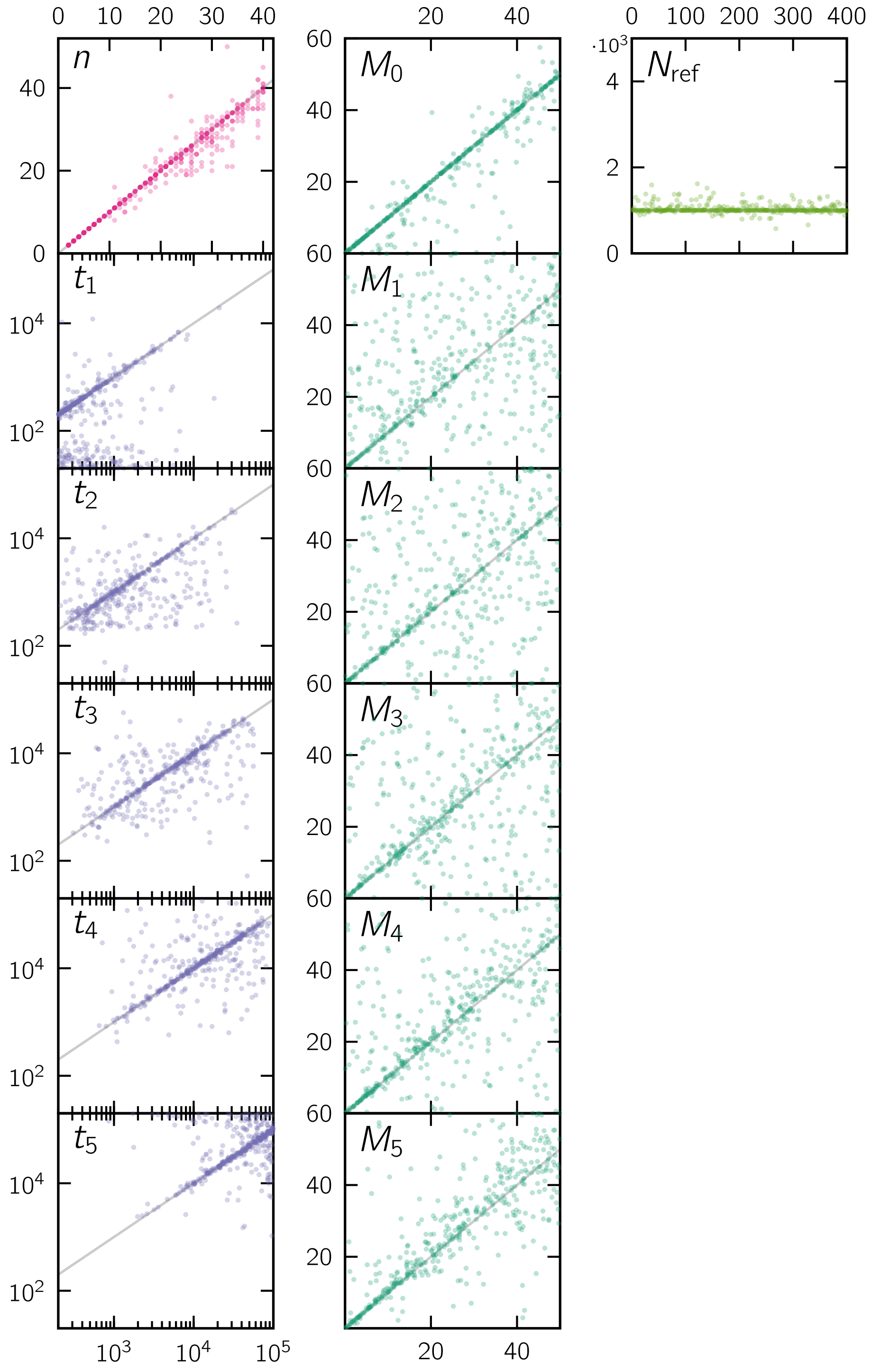

**Figure S14:** Scatter plots of all the simulated (horizontal axis) versus inferred (vertical axis) parameter values for scenarios of $c = 6$ components.

### 4. Results of validation using T-sim IICRs

In this section we show the validation results of simulating and then inferring from 100 randomly generated demographic scenarios with scaled IICRs and varying number of components $c$. In all cases, the scenario parameters were drawn from the following finite sets:

$$\begin{aligned} n &\in \{2, 5, 10, 15, 20\}, \\ t_i &\in \{0.1, 0.5, 1, 2, 5, 10, 20, 50\} \quad \forall i, \\ M_i &\in \{0.1, 0.2, 0.5, 1, 2, 5, 10, 20, 50\} \quad \forall i, \\ s_i &= 1 \quad \forall i, \\ N &= 1000. \end{aligned} \tag{3}$$

Unlike in the previous sections, the simulated IICRs are T-sim IICRs, meaning that the values are not exact due to the stochastic nature of the underlying ms simulation. For each value of $c$ from $c = 1$ to $c = 5$ we show the aggregate connectivity graph for all the simulations as well as the IICR and parameters of two individual scenarios from the set.

#### 4.1. Scenarios with 1 component

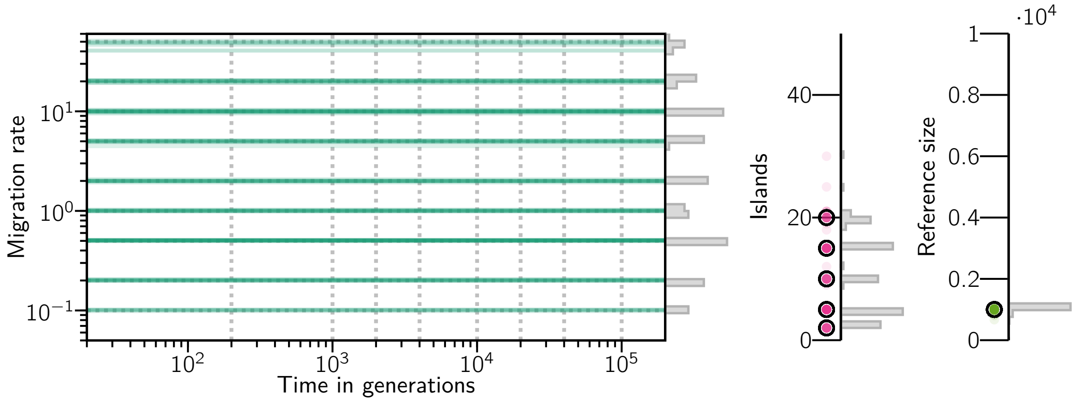

**Figure S15:** Connectivity graph of 100 inferred demographic histories using simulated T-sim IICRs of $c = 1$ components with simulated parameters randomly drawn from (3) and represented here by the dashed gray lines in the connectivity graph and the bold black circles in the islands and reference size plots.

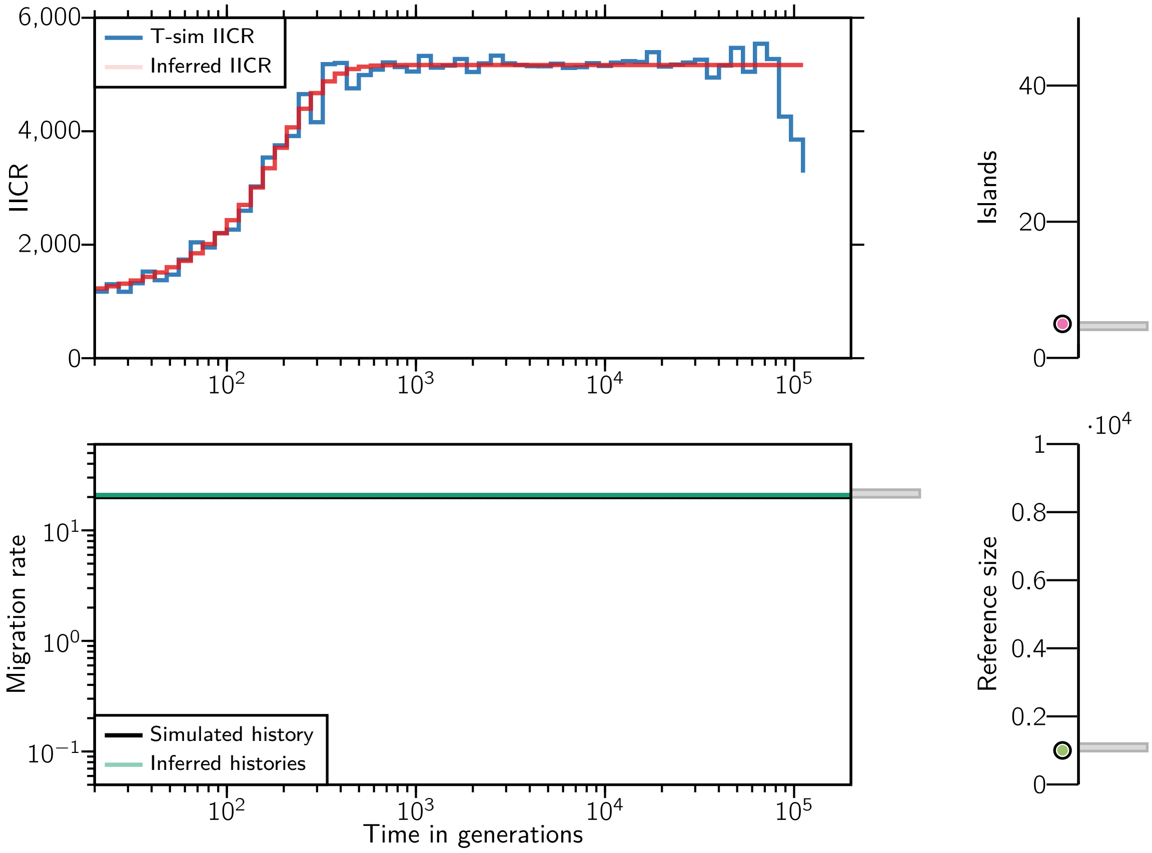

**Figure S16:** IICRs, connectivity graph, number of islands and reference size for one of the 100 simulated scenarios with $c = 1$ component and 10 independent inferences. The inferred scenario corresponds to $n = 5$, $M = 20$ and $N = 1000$.

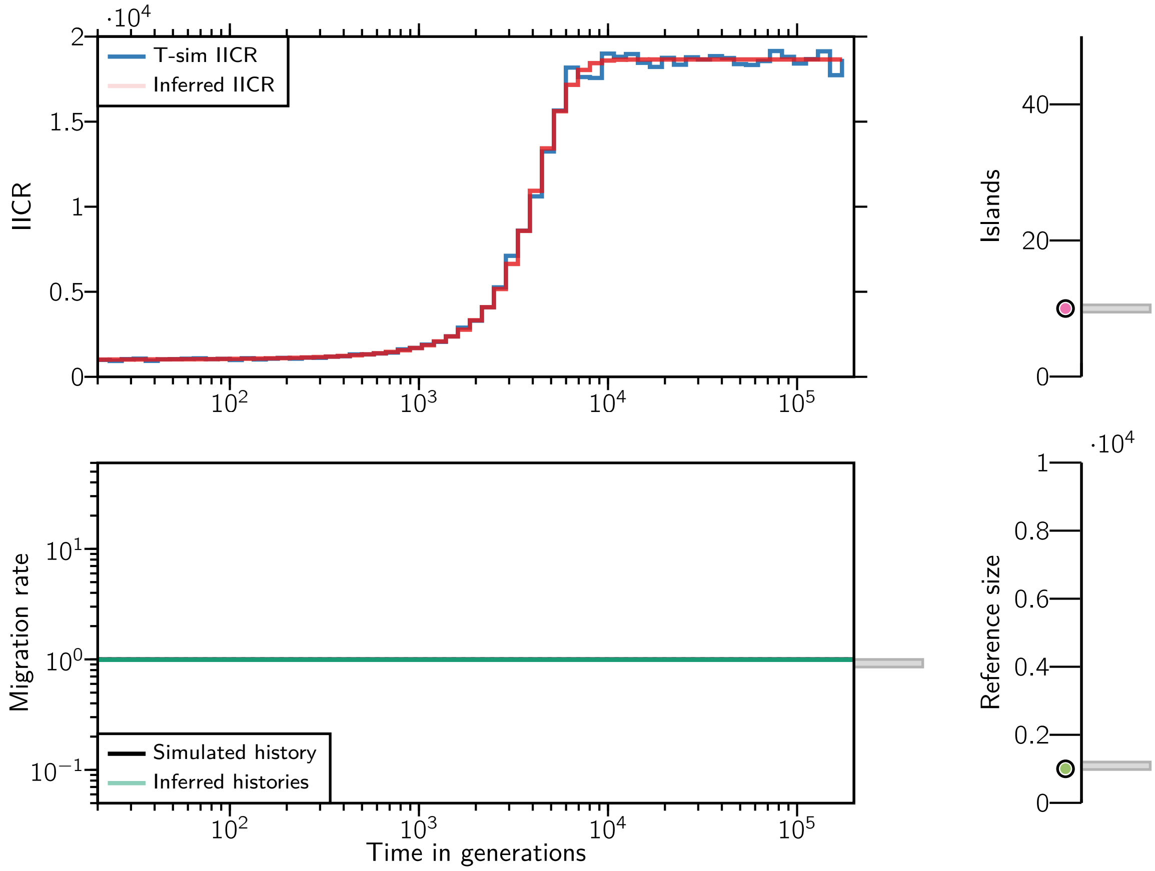

**Figure S17:** IICRs, connectivity graph, number of islands and reference size for one of the 100 simulated scenarios with $c = 1$ component and 10 independent inferences. The inferred scenario corresponds to $n = 10$, $M = 1$ and $N = 1000$.

#### 4.2. Scenarios with 2 components

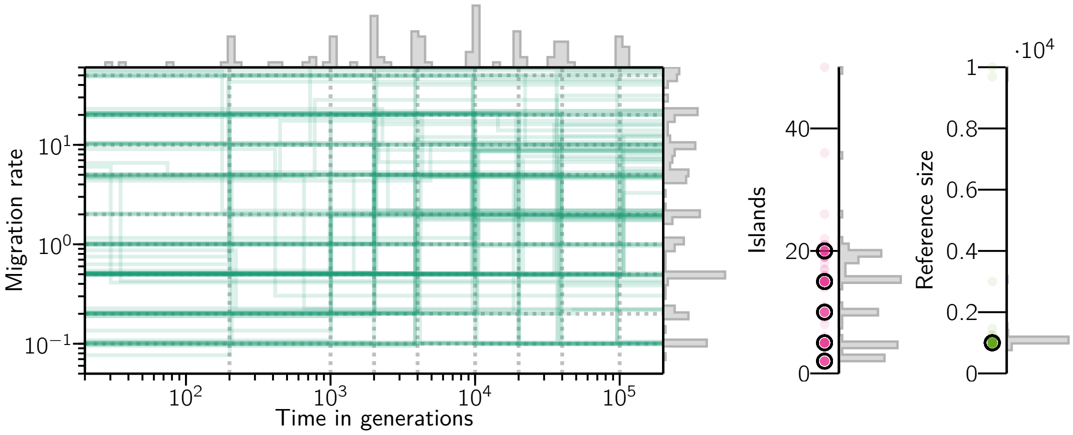

**Figure S18:** Connectivity graph of 100 inferred demographic histories using simulated T-sim IICRs of $c = 2$ components with simulated parameters randomly drawn from (3) and represented here by the dashed gray lines in the connectivity graph and the bold black circles in the islands and reference size plots.

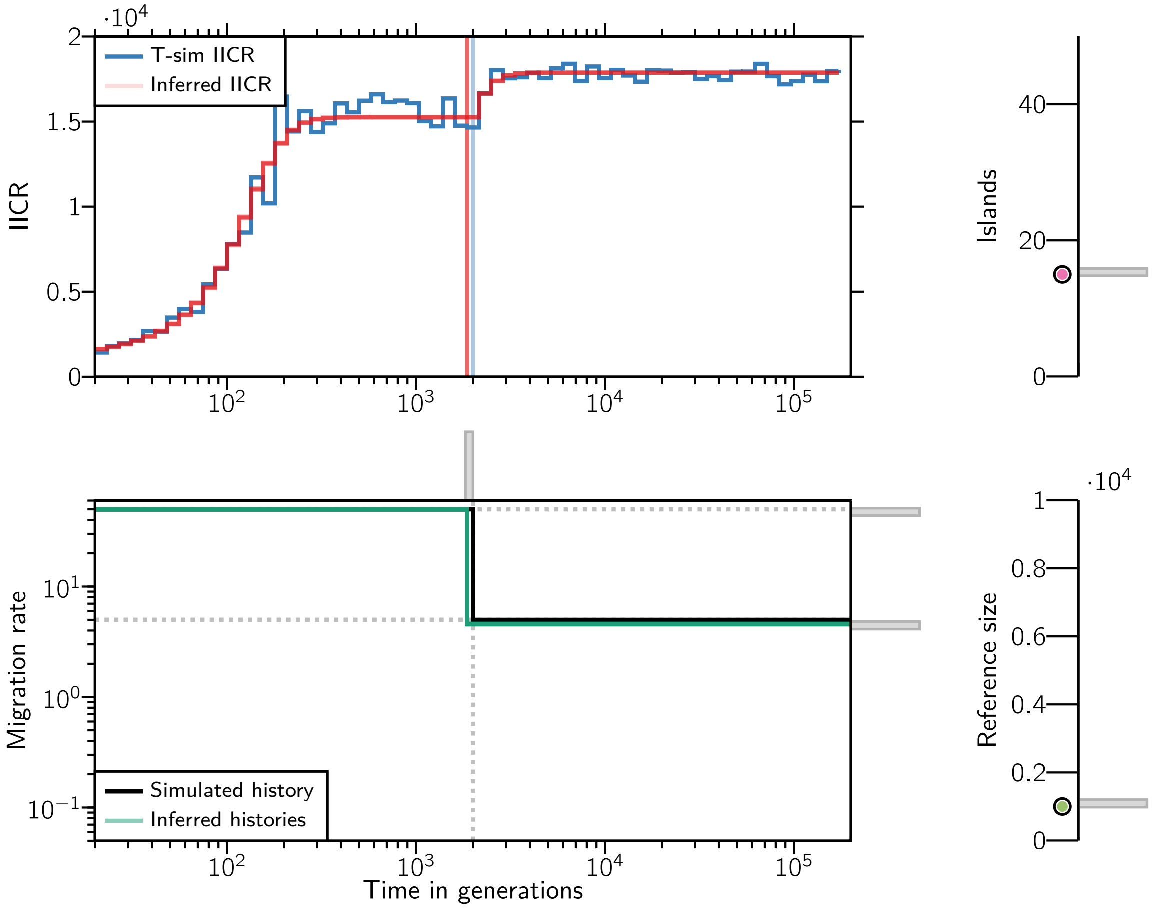

**Figure S19:** IICRs, connectivity graph, number of islands and reference size for one of the 100 simulated scenarios with $c = 2$ components and 10 independent inferences. The inferred scenario corresponds to $n = 15$, $t = 1$, $M = (50, 5)$ and $N = 1000$.

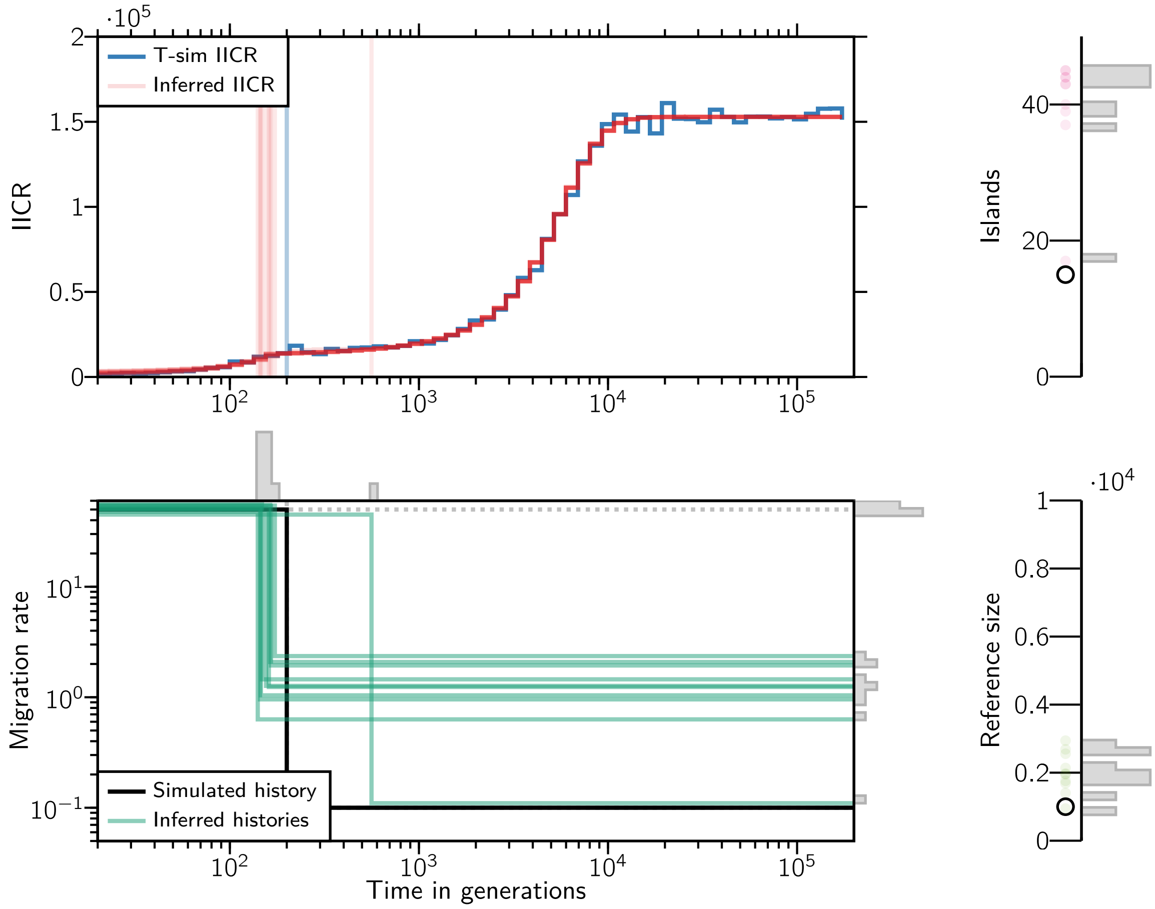

**Figure S20:** IICRs, connectivity graph, number of islands and reference size for one of the 100 simulated scenarios with $c = 2$ components and 10 independent inferences. The inferred scenario corresponds to $n = 15$, $t = 0.1$, $M = (50, 0.1)$ and $N = 1000$.

#### 4.3. Scenarios with 3 components

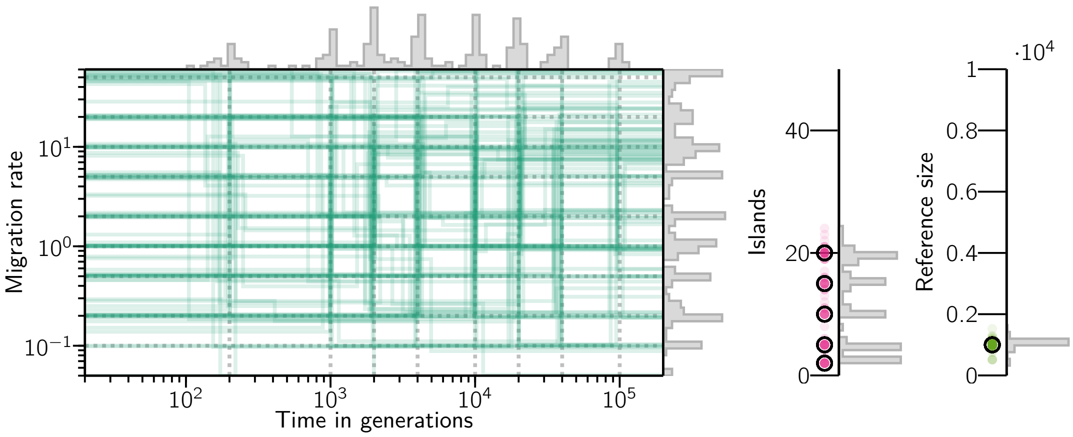

**Figure S21:** Connectivity graph of 100 inferred demographic histories using simulated T-sim IICRs of $c = 3$ components with simulated parameters randomly drawn from (3) and represented here by the dashed gray lines in the connectivity graph and the bold black circles in the islands and reference size plots.

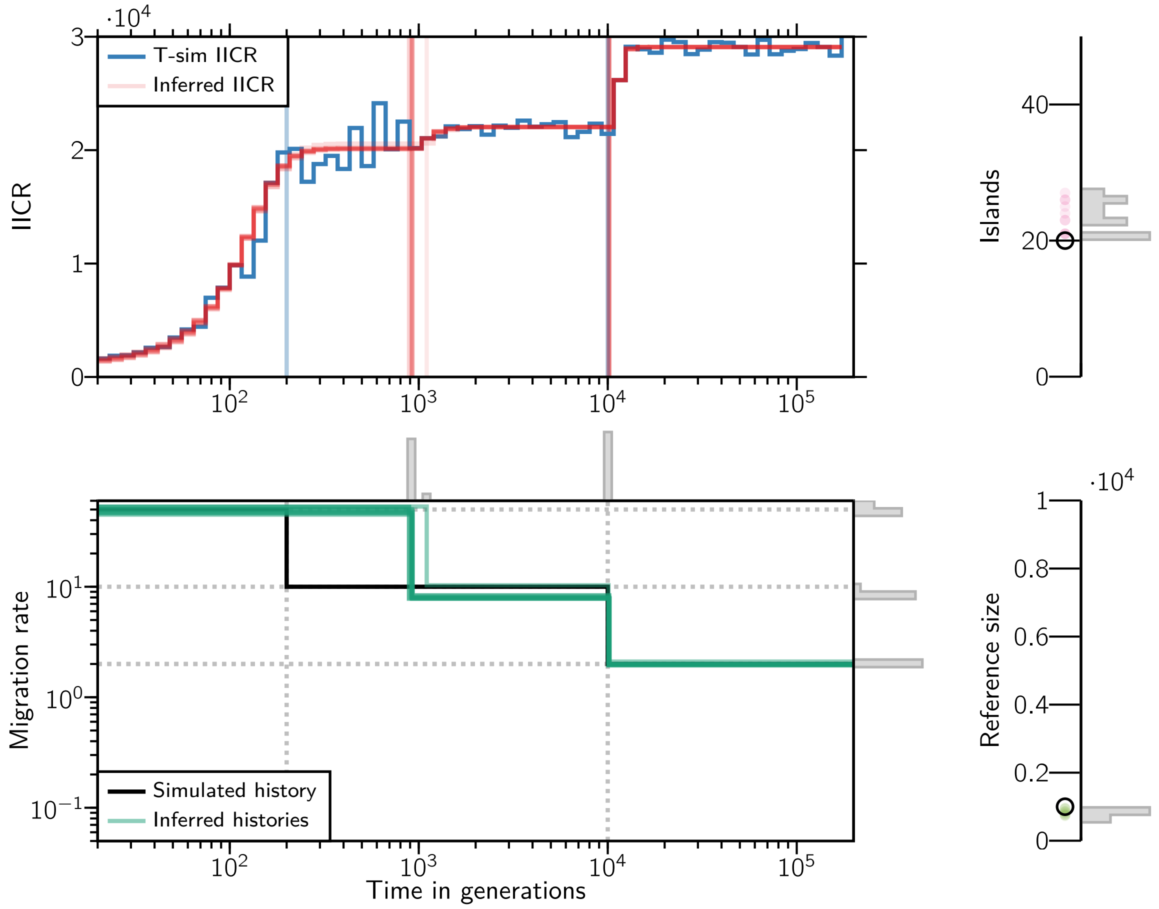

**Figure S22:** IICRs, connectivity graph, number of islands and reference size for one of the 100 simulated scenarios with $c = 3$ components and 10 independent inferences. The inferred scenario corresponds to $n = 20$, $t = (0.1, 5)$, $M = (50, 10, 2)$ and $N = 1000$.

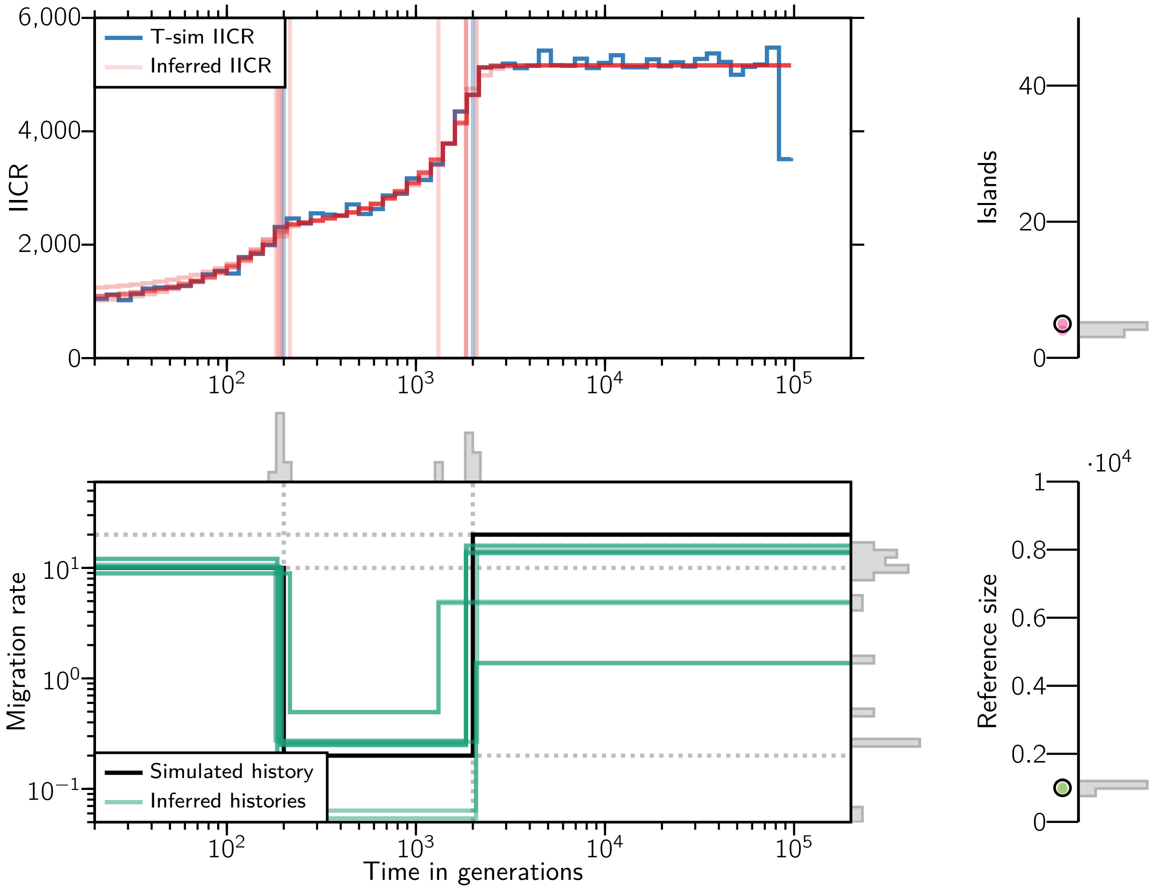

**Figure S23:** IICRs, connectivity graph, number of islands and reference size for one of the 100 simulated scenarios with $c = 3$ components and 10 independent inferences. The inferred scenario corresponds to $n = 5$, $t = (0.1, 1)$, $M = (10, 0.2, 20)$ and $N = 1000$.

#### 4.4. Scenarios with 4 components

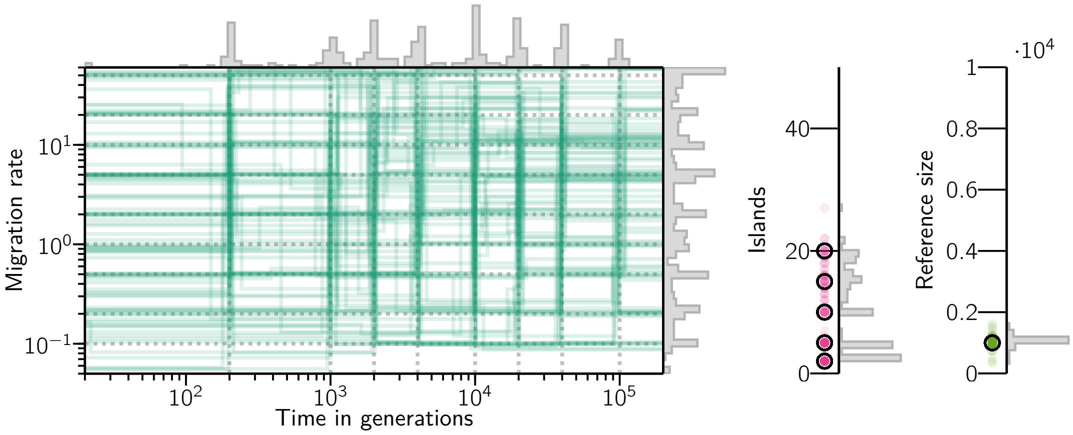

**Figure S24:** Connectivity graph of 100 inferred demographic histories using simulated T-sim IICRs of $c = 4$ components with simulated parameters randomly drawn from (3) and represented here by the dashed gray lines in the connectivity graph and the bold black circles in the islands and reference size plots.

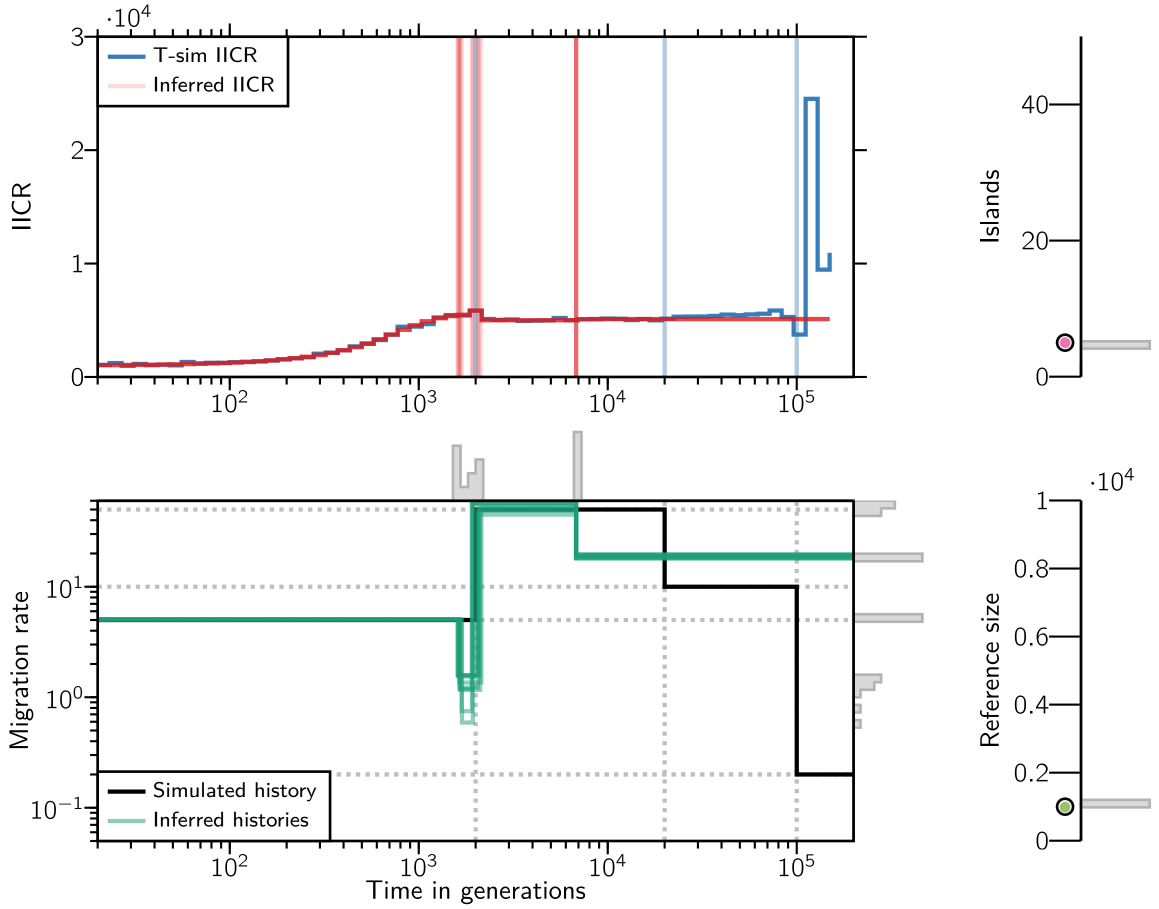

**Figure S25:** IICRs, connectivity graph, number of islands and reference size for one of the 100 simulated scenarios with $c = 4$ components and 10 independent inferences. The inferred scenario corresponds to $n = 5$, $t = (1, 10, 50)$, $M = (5, 50, 10, 0.2)$ and $N = 1000$.

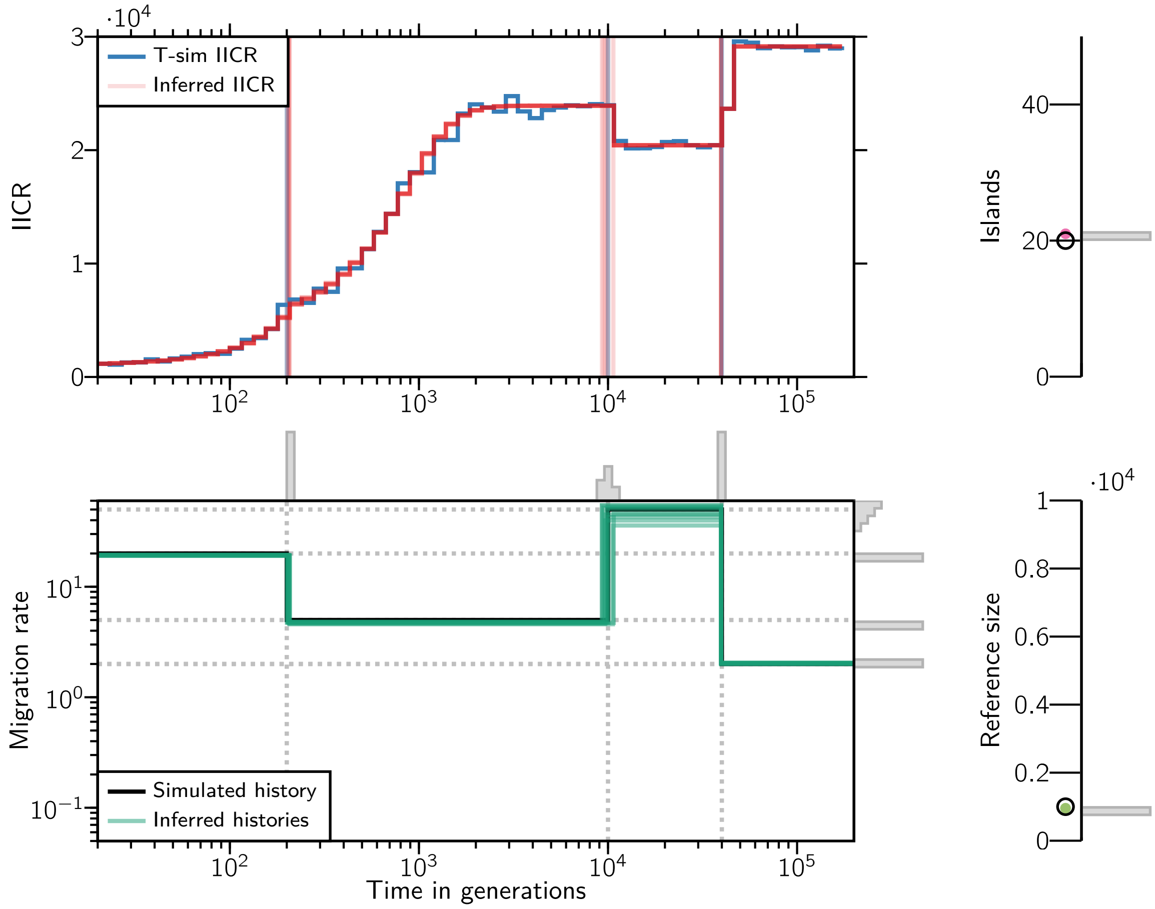

**Figure S26:** IICRs, connectivity graph, number of islands and reference size for one of the 100 simulated scenarios with $c = 4$ components and 10 independent inferences. The inferred scenario corresponds to $n = 20$, $t = (0.1, 5, 20)$, $M = (20, 5, 50, 2)$ and $N = 1000$.

#### 4.5. Scenarios with 5 components

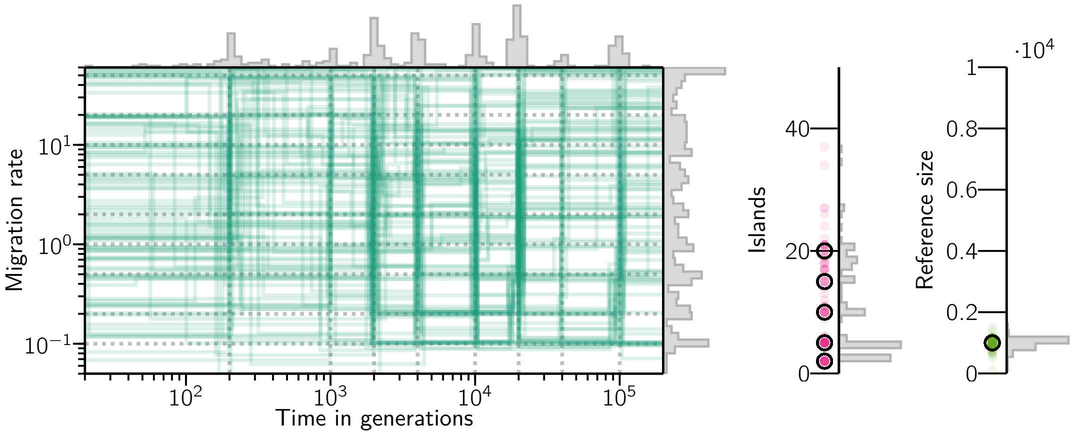

**Figure S27:** Connectivity graph of 100 inferred demographic histories using simulated T-sim IICRs of $c = 5$ components with simulated parameters randomly drawn from (3) and represented here by the dashed gray lines in the connectivity graph and the bold black circles in the islands and reference size plots.

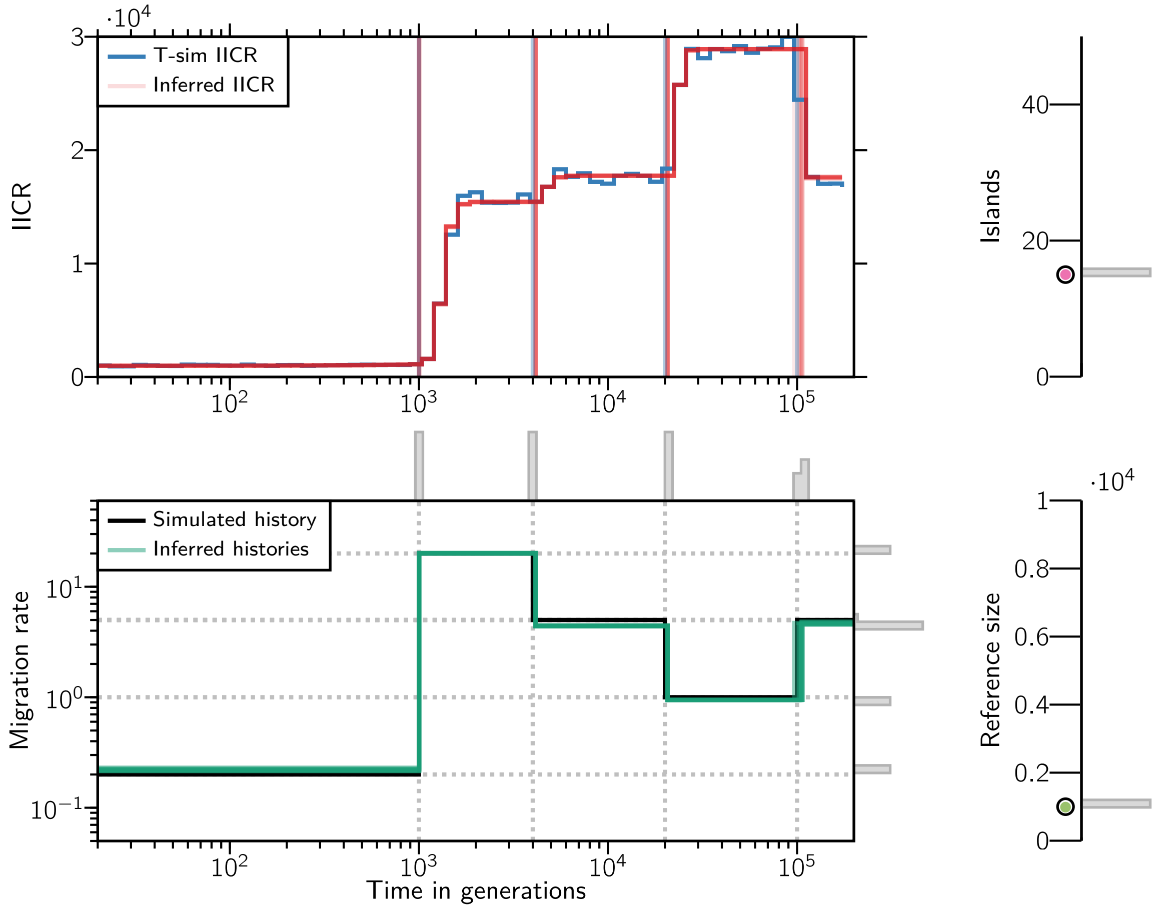

**Figure S28:** IICRs, connectivity graph, number of islands and reference size for one of the 100 simulated scenarios with $c = 5$ components and 10 independent inferences. The inferred scenario corresponds to $n = 15$, $t = (0.5, 2, 10, 50)$, $M = (0.2, 20, 5, 1, 5)$ and $N = 1000$.

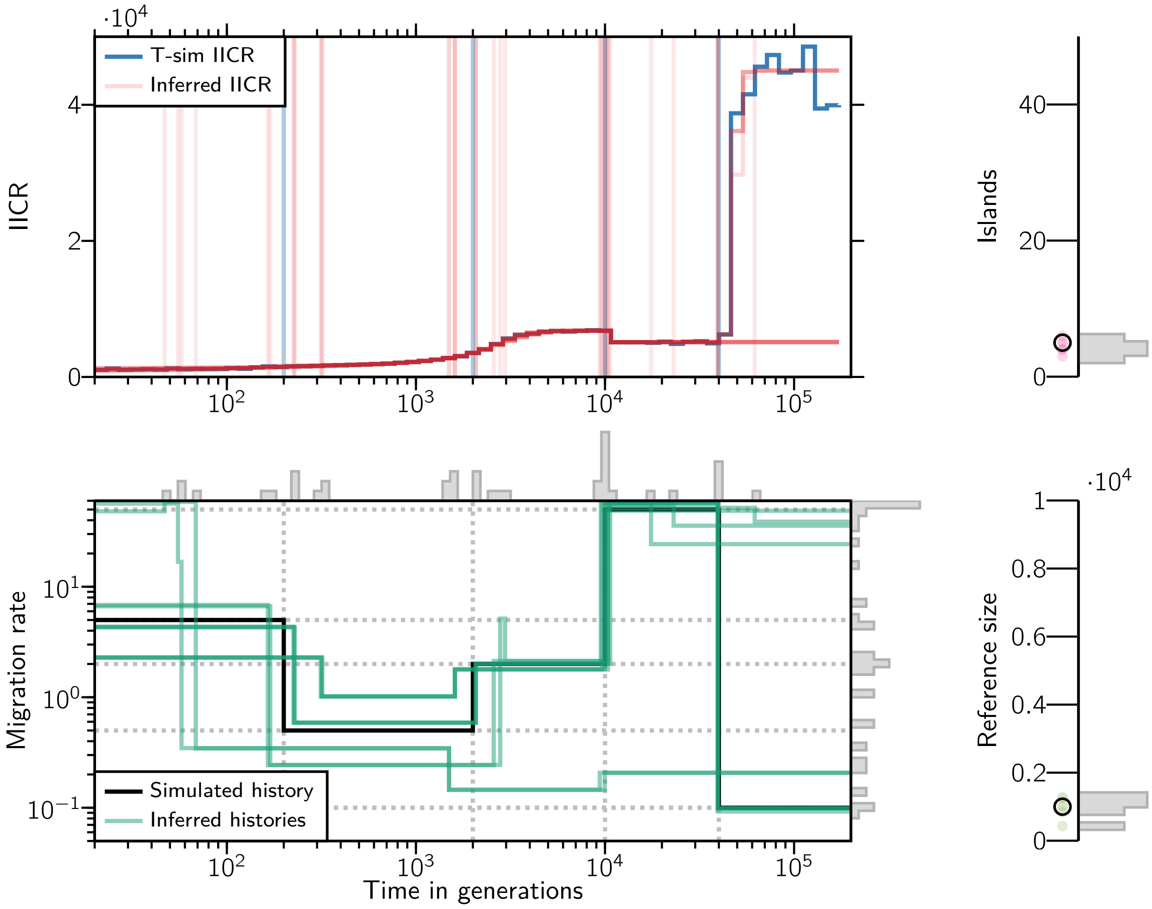

**Figure S29:** IICRs, connectivity graph, number of islands and reference size for one of the 100 simulated scenarios with $c = 5$ components and 10 independent inferences. The inferred scenario corresponds to $n = 5$, $t = (0.1, 1, 5, 20)$, $M = (5, 0.5, 2, 50, 0.1)$ and $N = 1000$.

### 5. Results of application to human data

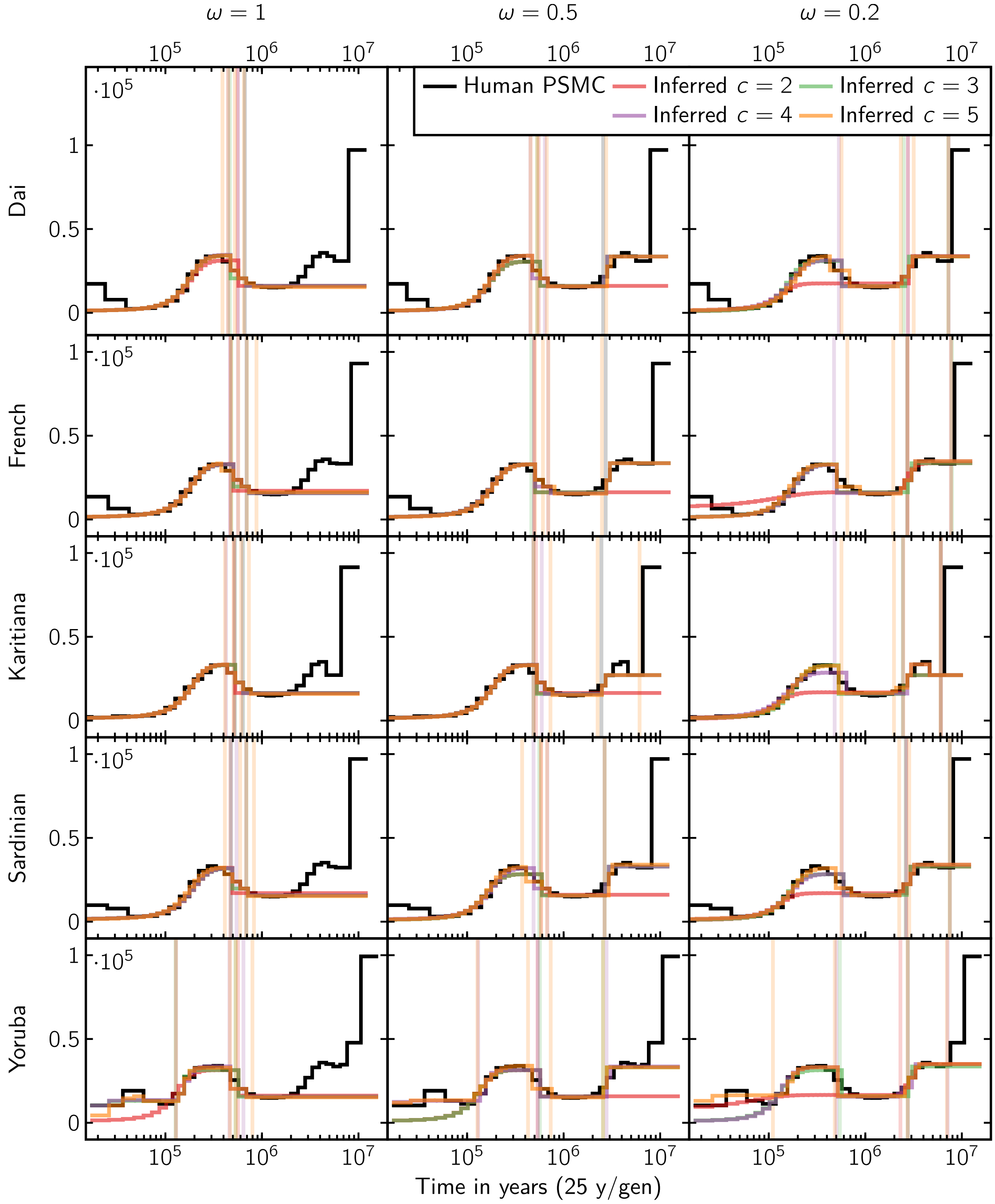

**Figure S30:** IICRs of the inferred human scenarios.

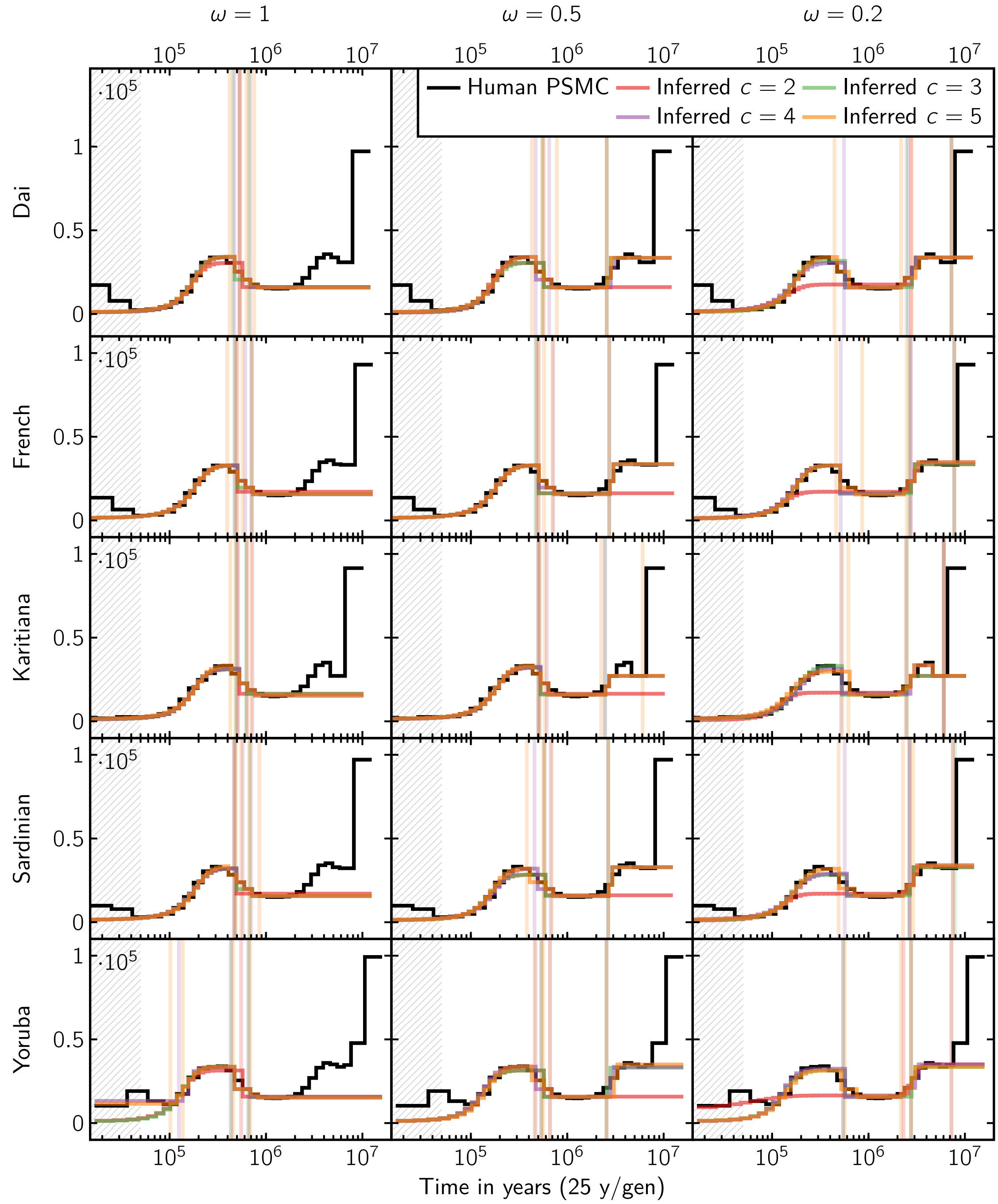

**Figure S31:** IICRs of the inferred human scenarios with restricted inference range.

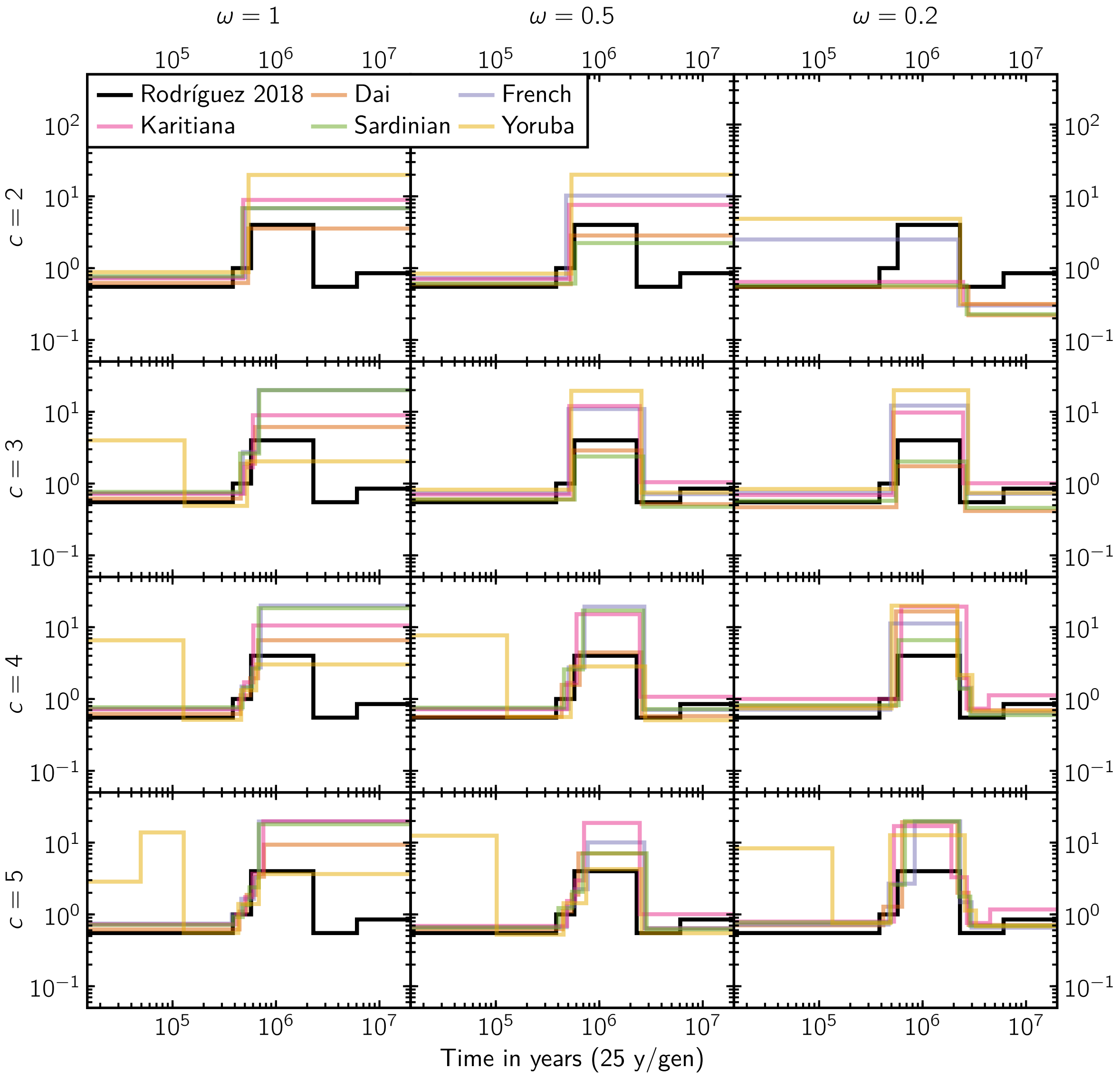

**Figure S32:** Connectivity graphs of the inferred human scenarios.

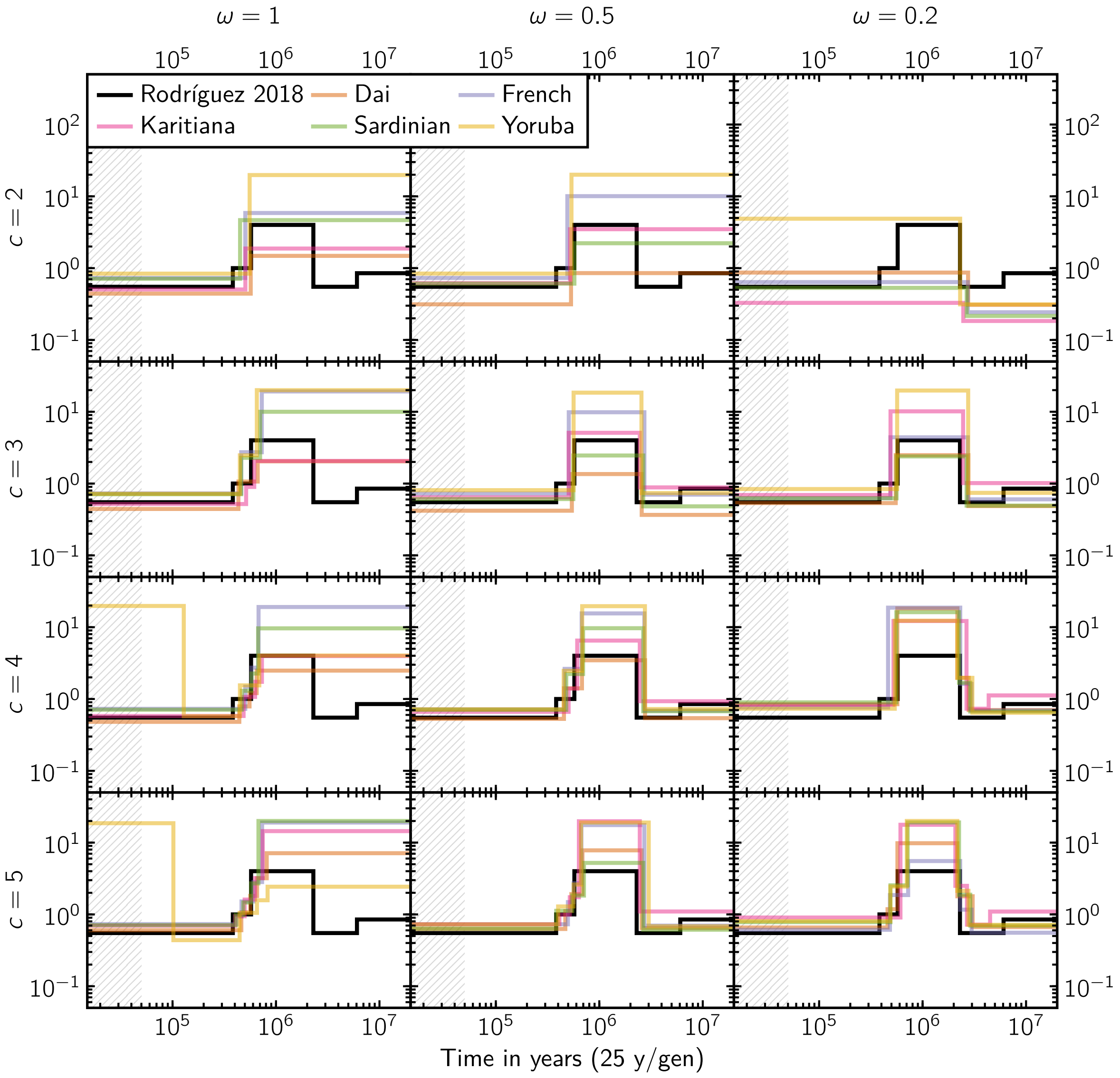

**Figure S33:** Connectivity graphs of the inferred human scenarios with restricted inference range.

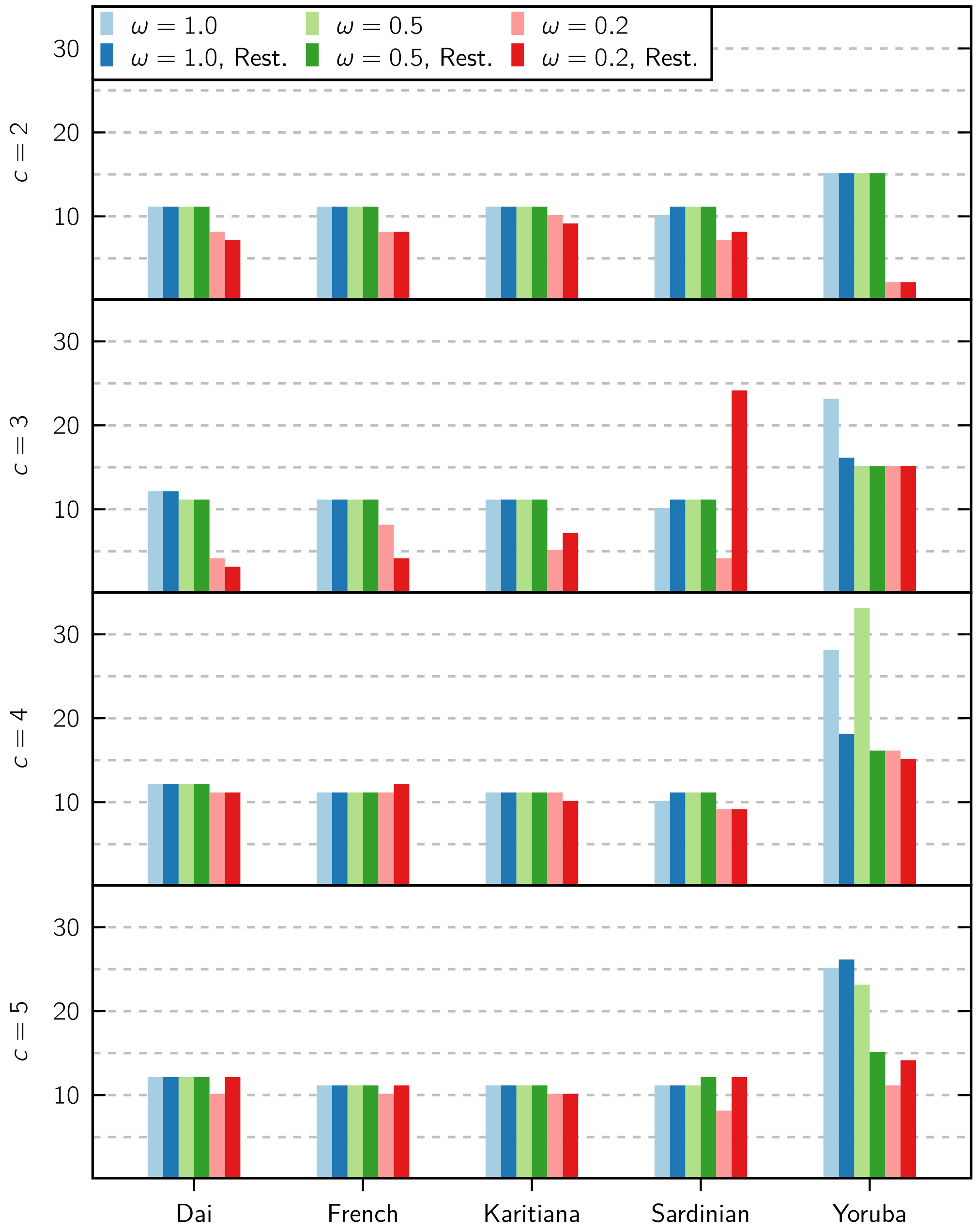

**Figure S34:** Inferred number of islands for the human populations.

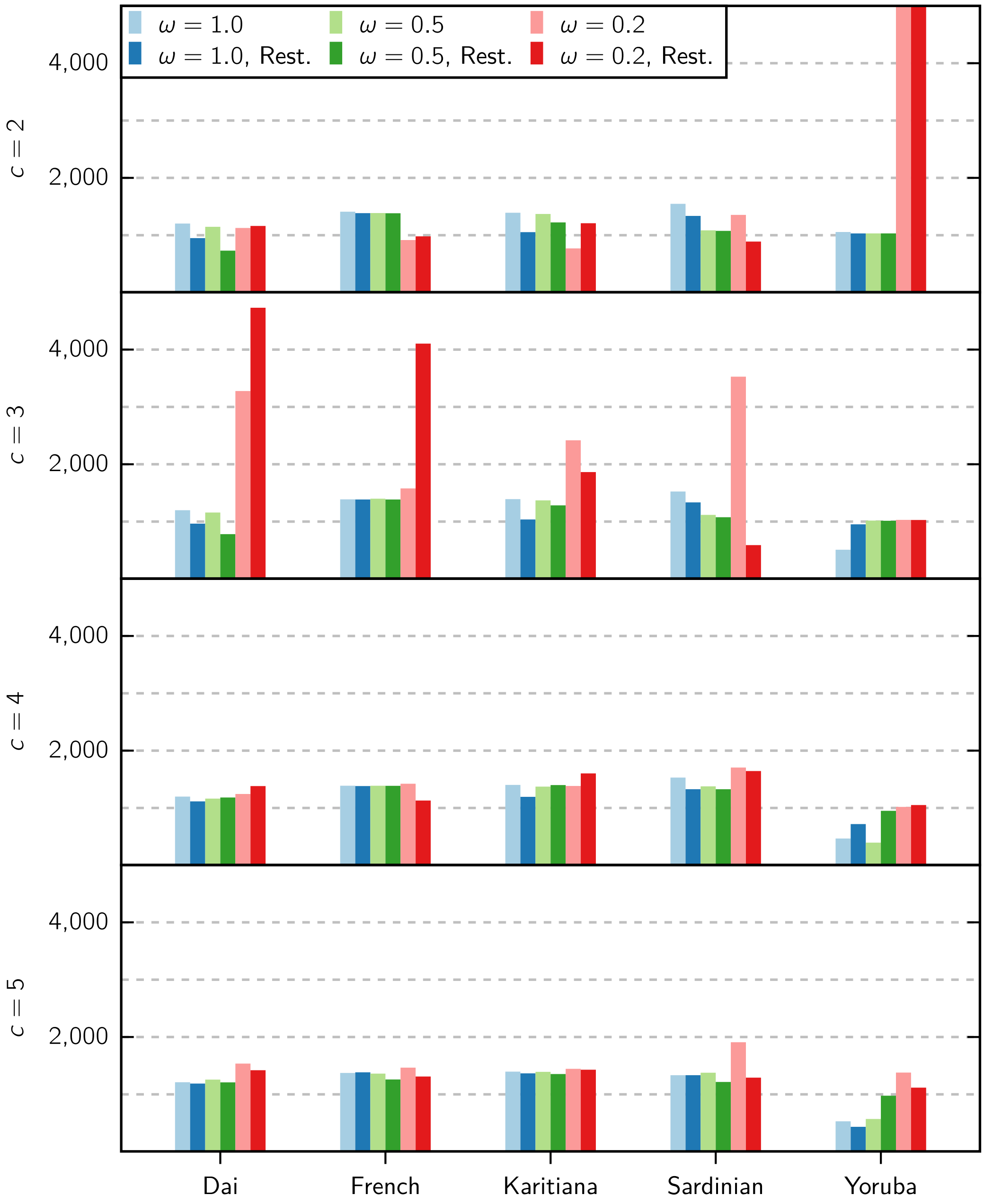

**Figure S35:** Inferred reference sizes for the human populations.

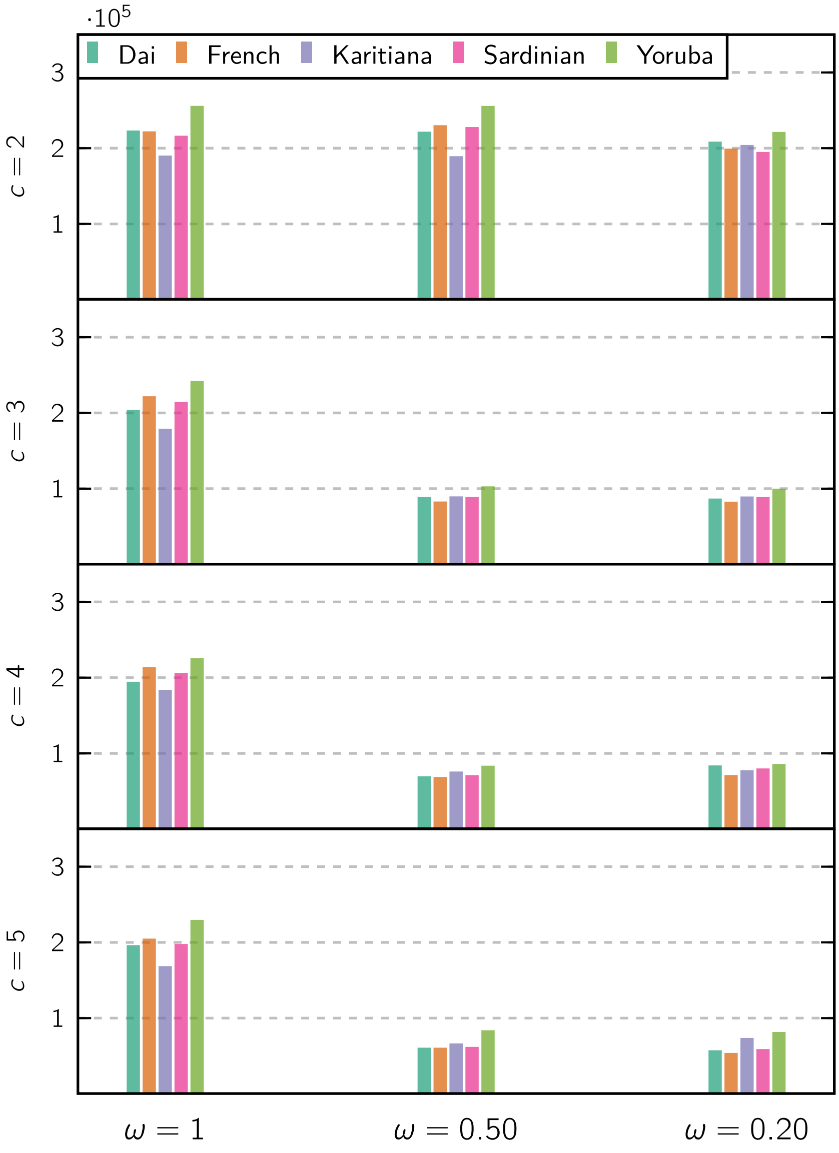

**Figure S36:** Visual distance ($d_{\text{visual}}$) of the best fitting scenario as a function of $\omega$ for various number of components.

### 6. A note on implementation and use cases

Our method is implemented in a program named SNIF (Structured Non-stationary Inference Framework) which will be freely available upon acceptance of the manuscript. The method produces parameter estimates and connectivity graphs in order to determine whether there is consistency across individuals or species. We focused on models in which the population size was maintained constant but the method can in principle infer changes in population size. This should be part of an extension of the present work. At this stage we stress that more work is needed before changes in connectivity and population size can be estimated together with confidence.

The intended use case for our method consists of several inference cycles, each preceded by adjustments to the many available parameters which include: the bounds $B$ for the estimated variables, the number of components $c$, the distance parameter $\omega$, the time interval where the distance is to be computed, the number of inference rounds along with the tolerance $\varepsilon$, some parameters of the search algorithm, and other minor options. This cycle emerges naturally from the fact that the inference process itself runs fairly quickly (a few seconds per round), so it is feasible to prepare a script that generates inferences under several combinations of these parameters and later do a general assessment of what are the most plausible scenarios for the data depending on the visual fit, the consistency of the inferred demographic histories, the distribution of the distances, etc. The SNIF software already includes a number of auxiliary scripts that may be used in this later analysis stage, including for instance the automatic generation of figures similar to the sub-panels of figures 9 and 10, where the target and inferred IICRs can be compared, and the nature of the best fitting scenarios can be understood using the connectivity graphs or the $n$ and $N$ plots.

As for the number of components $c$, specifying a higher value will generally result in a better fit and a lower distance, but is also incurs in longer analysis times and diminishing returns on the new information present in the resulting inferred histories. These aspects can be balanced by increasing the value of $c$ up to the point where the inferred demographic histories start converging (for instance, as more components are added, the connectivity graphs stop revealing new major events and any additional new degrees of freedom are only used to refine already existing details).

As part of the user inference cycles mentioned above, the bounds system can be configured to infer parameters for more strict or specialized symmetrical island models. For instance, it is possible to fix the number of inferred islands to be exactly 3 by setting $n_{\text{min}} = n_{\text{max}} = 3$ in the inference bounds $B$. Likewise, given an independent estimate of the reference size $N$ for the data, this information may be used to process the corresponding scaled target IICR as an unscaled one using matching inference bounds for the parameter $N$.

An important point concerning scaling is that the supplied value of the mutation rate $\mu$ must be accurately specified when inferring demographic parameters from a PSMC curve. This value is not used during the inference, but it is used in order to properly scale the PSMC curve and convert it into a target IICR, therefore it's important to provide the same value of $\mu$ that was used during the PSMC analysis. Otherwise, the only parameter that can be correctly inferred is $n$, since the rest of the parameters would only be accurate up to a scaling factor. This follows from a similar logic to that of Lemma 1.

We also suggest to use the hand-fitting Python script developed in Chikhi et al. (2018) as a complement to the automated inference process proposed in this work. You may want to compare the results obtained this way with the output of SNIF. Doing this will help make sense of the data, or help set the bounds for the many SNIF parameters.

### References

Chikhi, L., Rodríguez, W., Grusea, S., Santos, P., Boitard, S., and Mazet, O. (2018). The IICR (inverse instantaneous coalescence rate) as a summary of genomic diversity: insights into demographic inference and model choice. *Heredity*, 120:13–24.
